# qcGEM: a graph-based molecular representation with quantum chemistry awareness

**DOI:** 10.1101/2025.11.02.686183

**Authors:** Haoyu Wang, Haipeng Gong

## Abstract

The advancement of artificial intelligence (AI) has reshaped drug discovery. AI-based models typically rely on molecular representations for prediction. However, the absence of physically grounded information in mainstream molecular representations not only limits the model performance in practical applications, but also hinders the mechanistic understanding and exploitation by human. To overcome this issue, we introduce qcGEM, a quantum-chemistry-aware graph-based embedding of molecules that incorporates physical priors into molecular representation learning. By integrating quantum chemistry knowledge with a physics-inspired architecture, qcGEM provides a compact, physics-informed molecular representation that supports a diverse range of downstream applications. Particularly, qcGEM demonstrates the state-of-the-art performance across a broad range of molecule-related benchmarks, as evidenced by comprehensive evaluations on 71 tasks including molecular property prediction, activity cliff detection, protein-ligand interaction modeling and opioid drug classification, and simultaneously offers strong interpretability at multiple representation levels. We additionally propose a simplified variant, qcGEM-Hybrid, with substantially accelerated embedding generation and robust performance. Overall, our method provides an advanced molecular representation that will benefit molecule-related modeling and prediction, supporting further progress in AI-aided drug discovery.

## Introduction

Drug molecules exert therapeutic effects by modulating the activity of macromolecular targets. Therefore, drug discovery typically begins with target identification, followed by exploring the large chemical space for potential binders^1^. Yet, most early hits fail during development, due to assay artifacts, promiscuous reactivity, unacceptable toxicity, poor pharmacokinetics, and/or inadequate potency^2^. Consequently, early triage strategies that incorporate physicochemical properties, ADME (*i.e*. absorption, distribution, metabolism and excretion) profiles, safety assessments as well as binding-mode predictions are essential for enriching truly drug-like candidates^3^. Driven by advances in deep learning, numerous computational approaches have emerged for filtering drug candidates, enormously accelerating the entire process^4–10^. Compared to traditional physics-based approaches^11–13^, deep learning models offer markedly faster inference once training is complete, and support more flexible and more richly defined prediction tasks dictated by the available training labels.

Since deep learning models are fundamentally data-driven, their performance depends critically on the quality and relevance of input data and labels^14^. In the past, numerous molecular representations have been developed and employed as input for deep learning models in the field of drug discovery, which generally fall into two categories: handcrafted features and embeddings learned from neural networks. SMILES^15^ and fingerprints^16^ encode a molecule as a linear text string defined by explicit syntax rules, yielding a compact 1D representation that inherently fits in large language models and thus holding a ubiquitous status in molecular machine learning^5,17^. Based on advances in computer vision, several studies convert molecules into images and feed them to convolution-based models for downstream property prediction^6,7^. In addition, molecules are often naturally represented as atom-bond graphs and processed by a range of graph neural network (GNN) architectures^8–10^. Albeit widely used, these handcrafted representations usually suffer from information redundancy, non-unique representations, insufficient structural context, and slow convergence at downstream training. In recent years, an increasing number of deep-learning-based molecular representation approaches have been proposed^18–33^, which reportedly achieve superior performance over traditional methods in various applications. These methods automatically distill higher-order semantic information from the pre-trained model and transfer with outstanding accuracy across downstream tasks. However, their performance is highly contingent on both the volume and quality of pre-training data, as well as on the design of pre-training strategy^34,35^. Despite the progress, rudimentary learning features and simplistic self-supervised frameworks in these methods result in high data demands and representational dimensionality, limited generalizability, incomplete structure encoding, and more importantly, the lack of physical interpretability.

Compared to conventional learning objectives, quantum chemical calculations offer more physically grounded targets for the pre-training process^36,37^. By solving the Schrödinger equation, complete quantum information of electrons can be obtained, which prescribes interactions between nuclei, including non-local internuclear communications that are of critical importance in conjugate molecular systems but are often neglected by traditional methods^38^. In principle, electronic wave functions contain full quantum information for a specified configuration of atomic nuclei. However, deficits like the high dimensionality, numerical instability and continuous distribution hinder their direct integration with deep learning models^39^. The electron density, a distribution function directly defined in the three-dimensional Cartesian space, contains equivalent quantum information with substantially reduced dimensionality. However, the continuous nature and the lack of rotational invariance jointly hinder its effective modeling in mainstream network architectures^40,41^. Localization of wave functions, as a further simplification approach, provides a separation of electronic orbitals into those residing on atomic centers and those falling in-between^42,43^, perfectly aligning with the node and edge inputs of GNNs. Additionally, the corresponding quantum descriptors also provide informative learning objectives to guide the model in generating more compact and physically meaningful molecular representations. In a prior work, we have constructed a quantum chemically annotated database qcMol^44^, which incorporates abundant quantum chemical descriptors and three-dimensional molecular configurations for over 1.2 million chemical compounds, laying the foundation for quantum-chemistry-based molecular representation learning.

In this study, we propose qcGEM, a <u>q</u>uantum-<u>c</u>hemistry-aware <u>g</u>raph-based <u>e</u>mbedding of <u>m</u>olecules. Through pre-training over a subset of molecules in the qcMol database, qcGEM learns to encode electronic quantum information and molecular structural features into a compact graph that preserves rich physicochemical and complete geometrical details. At the architectural level, the global-node-edge hierarchy in the encoder-decoder framework of our method enables the generation of a multiscale molecular embedding applicable for diverse prediction and molecular design tasks. To validate the advantage of our representation, we compared it with 29 widely adopted molecular embeddings. Results suggest that qcGEM comprehensively achieves the best performance on 71 datasets covering versatile molecular property and protein-ligand related tasks. Subsequently, we developed a simplified variant, qcGEM-Hybrid, by taking the trade-off between accuracy and speed. Allowing enhanced batch embedding generation at an acceptable speed for practical applications, this hybrid version maintains a comparably robust performance in downstream tasks. Finally, our analysis shows that the representation space of qcGEM is highly interpretable, presenting a promising potential for molecular design.

## Results

### Overview of framework

As shown in Figure 1a, the qcGEM framework generates representations for drug molecules in three steps: 1) searching from the qcMol library for rapid quantum chemical annotation, 2) pre-training an encoder-decoder model to learn a graph-based representation space from initial features, and 3) transferring the representation to practical molecules for downstream applications with frozen or fine-tuned settings. Viewing a chemical molecule as a system composed of atomic nuclei that interact through the potential energy enforced by electrons, qcGEM substitutes traditional sparse graph representations with a fully connected graph to better capture all pairwise atomic interactions. Through specialized designs in model architecture (Figure 1b), information exchange (Figure 1c) and pre-training strategy, qcGEM constructs a multi-level molecular representation (Figure 1d) that delivers robust generalization across diverse tasks while preserving strong physical interpretability.

**Figure 1.**
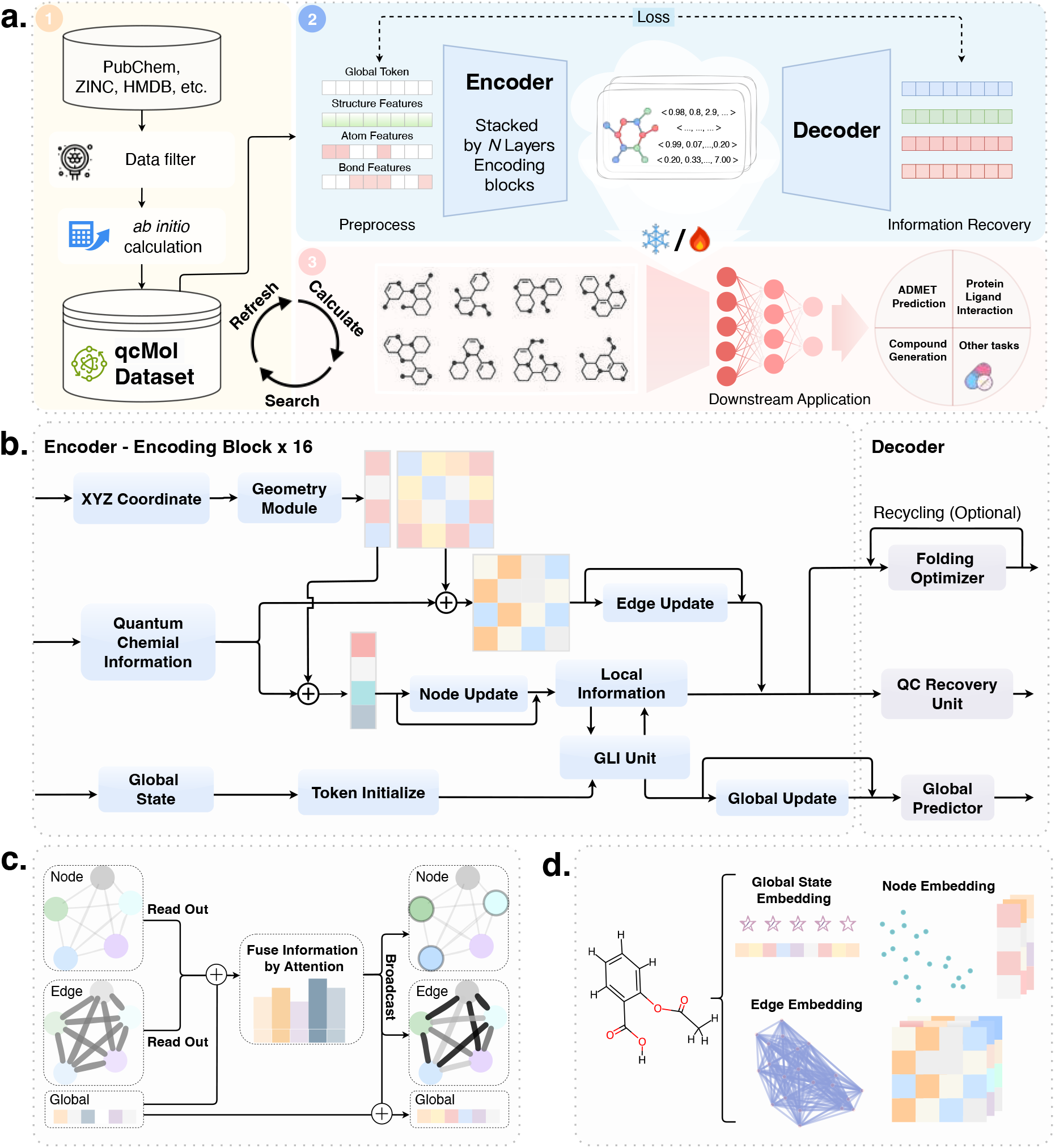
The model architecture of qcGEM. **a)** Schematic overview of the qcGEM framework. The compound is first queried from the qcMol database or computed via quantum chemical calculations. The qcGEM model encodes the initial features into a unified molecular representation, from which all input information could be reconstructed after decoding. The encoded representation is finally transferred to downstream tasks for diverse applications. **b)** The encoder-decoder architecture of the qcGEM model. The encoder consists of 16 repeated encoding blocks, and allows sufficient information exchange between the structural, quantum chemical and global features. The decoder is composed of a folding optimizer, a QC recovery unit and a global predictor, for the reconstruction of three-dimensional configuration, quantum chemical descriptors and global features, respectively. **c)** Model design of the GLI unit in panel (**d**) for effective representation learning of molecular information at the global, node and edge levels. **d)** Schematic show of the compact and multi-level graph representation generated by qcGEM for an exemplar molecule.

### Physics-inspired architecture and pre-training strategy

The model architecture of qcGEM consists of an encoder and a decoder (Figure 1b). The encoder employs stacked encoding blocks to embed quantum chemical information, three-dimensional molecular structure and other global properties into a compact, multi-level representation. The decoder leverages this learned representation to reconstruct molecular structure, quantum descriptors and global characteristics via the folding optimizer, the QC recovery unit and the global predictor, respectively. The information flow in qcGEM is physics-inspired, allowing iterative exchanges between information of various sources. First, the geometric information of atomic nuclei is encoded into nodes and edges from the complete molecular structure in a rotranslation-invariant manner. Next, these structure-aware features are fused into the localized electronic quantum information for node and edge updating. Finally, the global state and local node-edge information are exchanged in the global-local interaction (GLI) unit (Figure 1c) to produce a condensed global representation for the molecule. These designs enable efficient multi-scale learning and fast convergence, yielding a quantum-chemistry-aware representation in a compact form.

The pre-training tasks include geometry denoising, electronic feature recovery and correction, and global state condensation. For geometry denoising, qcGEM is specially designed with a geometry module for complete geometry encoding, a folding optimizer for effective structure decoding (Supplementary Figure 1a), and a tailored FAPE loss^45^ for constraining the structural reconstruction of small molecules (Supplementary Figure 1b). To capture electronic quantum information, qcGEM employs corruption and masking of node and edge features, followed by a correction and reconstruction process. The global state condensation enhances the permutation-invariant readout capability of graph embedding by feature decoupling^33,46^. These components jointly endow qcGEM the capacity to capture high-level semantics, while supporting multi-level molecular prediction and design at the same time.

qcGEM is pre-trained on only 300 thousand samples, requiring substantially fewer resources than many recently proposed methods (Supplementary Table 4), and supports applications in both parameter freezing and rapid fine-tuning settings. The fixed representations generally achieve sufficiently strong performance, and require only a single-pass inference for repeated uses. Pre-calculated representations are available for download through the qcMol database, while the embeddings for unseen molecules could be computed from scratch from the online server of qcGEM.

### Benchmark on molecular property prediction with qcGEM

To evaluate the performance of qcGEM, we compared it against mainstream hardcraft features and model-based embeddings (Supplementary Table 4) across a series of downstream tasks using a probe network of the multiple layer perceptron (MLP) architecture (Figure 2a). We first tested on molecular property prediction tasks, involving 30 widely used benchmarks collected from MoleculeNet^47^ and the TDC dataset^48^. These datasets cover molecular properties of multiple aspects including biophysics, physiology and physical chemistry, and are categorized into prediction tasks of absorption (8 sets), distribution (4 sets), metabolism (8 sets), excretion (3 sets), and toxicity (7 sets), which are abbreviated in conglomerate as ADMET properties (Supplementary Table 5).

**Figure 2.**
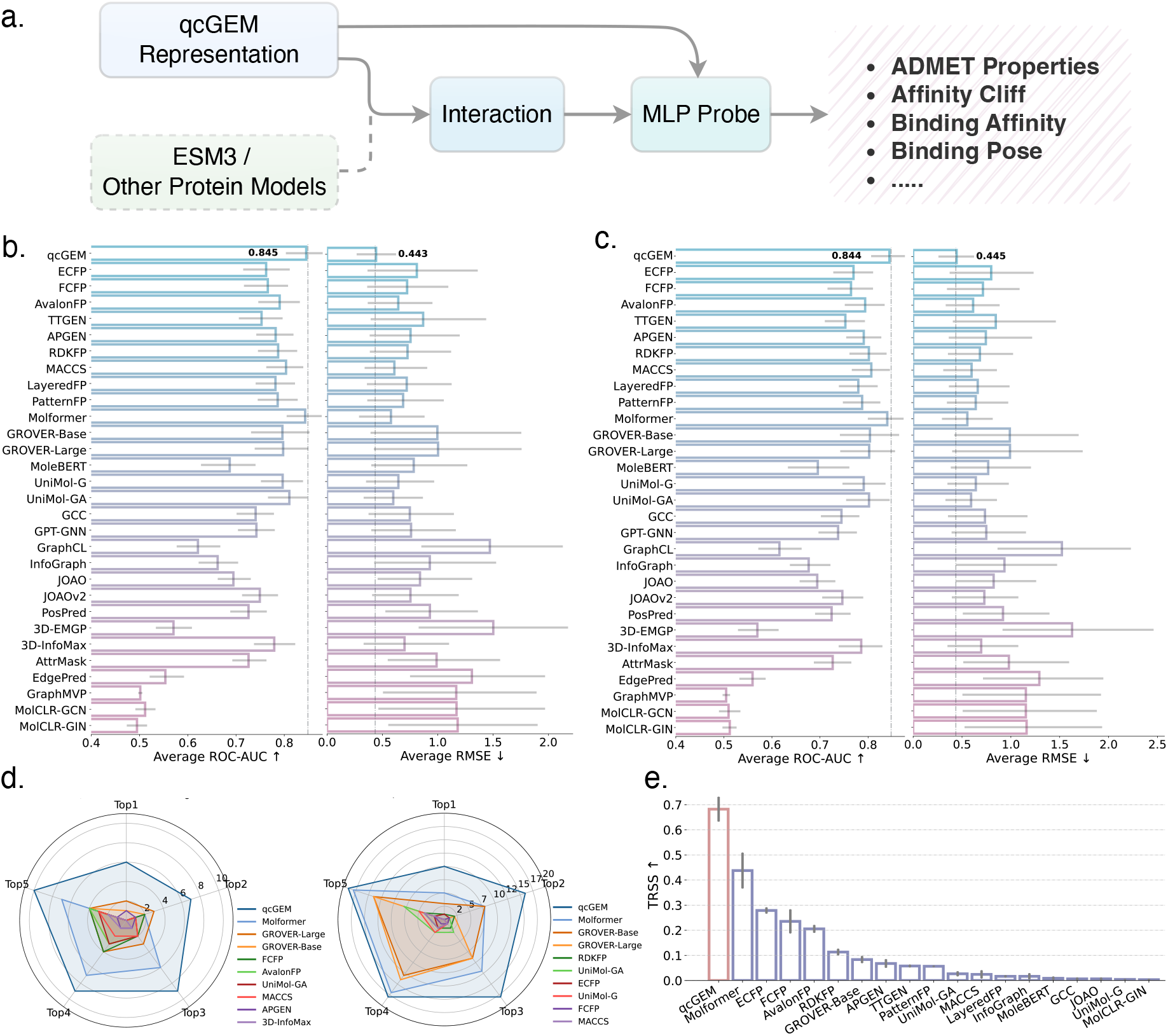
Performance of qcGEM on molecular properties prediction. **a)** In downstream evaluations, qcGEM and/or rival representations are directly utilized for prediction through an MLP probe network for compound-only tasks, but are first integrated with protein representation models via an interaction module for protein-related tasks. **b)** Comparison of the 30 molecular representations on 30 molecular property prediction tasks in the scaffold split setting. ROC-AUC is shown for classification tasks (left), while RMSE is reported for regression tasks (right), with arrows indicating the desired directions. Vertical dash-dotted lines indicate the best-performing results. **c)** Comparison of the 30 molecular representations on 30 molecular property prediction tasks in the random split setting. The results are presented in the same convention as in panel (**b**). **d)** Radar plots of the overall performance profiles for the top methods in the scaffold split setting. Here, occurring counts of each method in the top 1-5 rankings are shown for the regression tasks (left) and classification tasks (right), respectively. **e)** The overall TRSS scores of the top methods. Here, TRSS scores are calculated across split settings and task categories, and their mean values and standard deviations are shown in the bar plot.

The evaluation was performed in both the standard random split setting (*i.e*. random splitting of training and testing data) and the more challenging scaffold split setting (*i.e*. scaffold-based splitting of training and testing data), using the metric of root mean square error (RMSE) for regression tasks and area under curve (AUC) of the receiver operating characteristic (ROC) curve for classification tasks. As shown in Figure 2b&c, qcGEM achieves the state-of-the-art performance in both regression and classification tasks across both split settings. Particularly, our representation leads the other methods by a large margin in regression tasks. Its advantage in classification tasks is comparatively modest, likely due to the influence of toxicity tasks, which often require organism-specific omics data for accurate prediction. Detailed ranking information of qcGEM in all individual tasks and across different AMDET task categories could be found in Supplementary Figures 6 and 7, respectively. The radar plots in Figure 2d summarize the occurring counts of tested methods in the top *k* (*k* = 1, 2, 3, 4, 5) rankings in the scaffold split setting for regression and classification tasks, respectively. Clearly, qcGEM consistently outperforms the other methods in this respect (see Supplementary Figure 8 for detailed statistics). Here, we propose the top-ranking stability score (TRSS), a metric that mitigates the impact of occasional outliers by rewarding consistent top-tier performances (Equation 8), to comprehensively evaluate the performance and robustness of tested methods. Figure 2e presents the distribution of TRSS scores across four evaluation scenarios (two split settings and two task types). Again, qcGEM markedly outperforms the other methods in this metric.

Results on the ADMET datasets demonstrate that qcGEM offers a powerful molecular representation, leading to superior performance in molecular property prediction. Detailed per dataset results could be found in Supplementary Tables 9-20.

### Benchmark on activity cliff prediction with qcGEM

In drug screening, subtle structural modifications can lead to substantial changes in binding affinity^49,50^, as exemplified by the activity cliff molecules. To assess qcGEM’s sensitivity to such variations, we curated 30 tasks from the MoleculeACE dataset^51^, which incorporates experimental cases where minor structural changes result in pronounced differences in bioactivity (Supplementary Table 6). Following the convention of MoleculeACE, all ligands are categorized into common molecules and activity cliff molecules to evaluate the accuracy and generalization of different molecular representations.

As shown in Figure 3a, qcGEM on average achieves the lowest prediction error (in RMSE) across the 30 datasets, regardless of whether the evaluation involves common molecules or only focuses on activity cliff compounds. To assess performance across task types, we further categorized all testing data into 30 specific protein targets, 6 protein classes and 2 experimental categories, respectively (Supplementary Table 6). qcGEM exhibits strong performance across all task subdivisions among the 30 molecular representations, achieving the highest ranking in 70%, 83% and 100% of the corresponding task categories, respectively, for all tested molecules (Figure 3d). This superiority persists when the focus is shifted to activity cliff molecules (Supplementary Figure 9a). Moreover, either for all molecules or activity cliff molecules, qcGEM maintains a consistent advantage over the other methods in respect of the occurring counts within the top 1-5 rankings (Supplementary Figure 9b; see Supplementary Figure 10 for detailed statistics). We further calculated the TRSS score across all task types. As expected, qcGEM achieves the highest overall score, indicating its superior robustness (Figure 3c).

**Figure 3.**
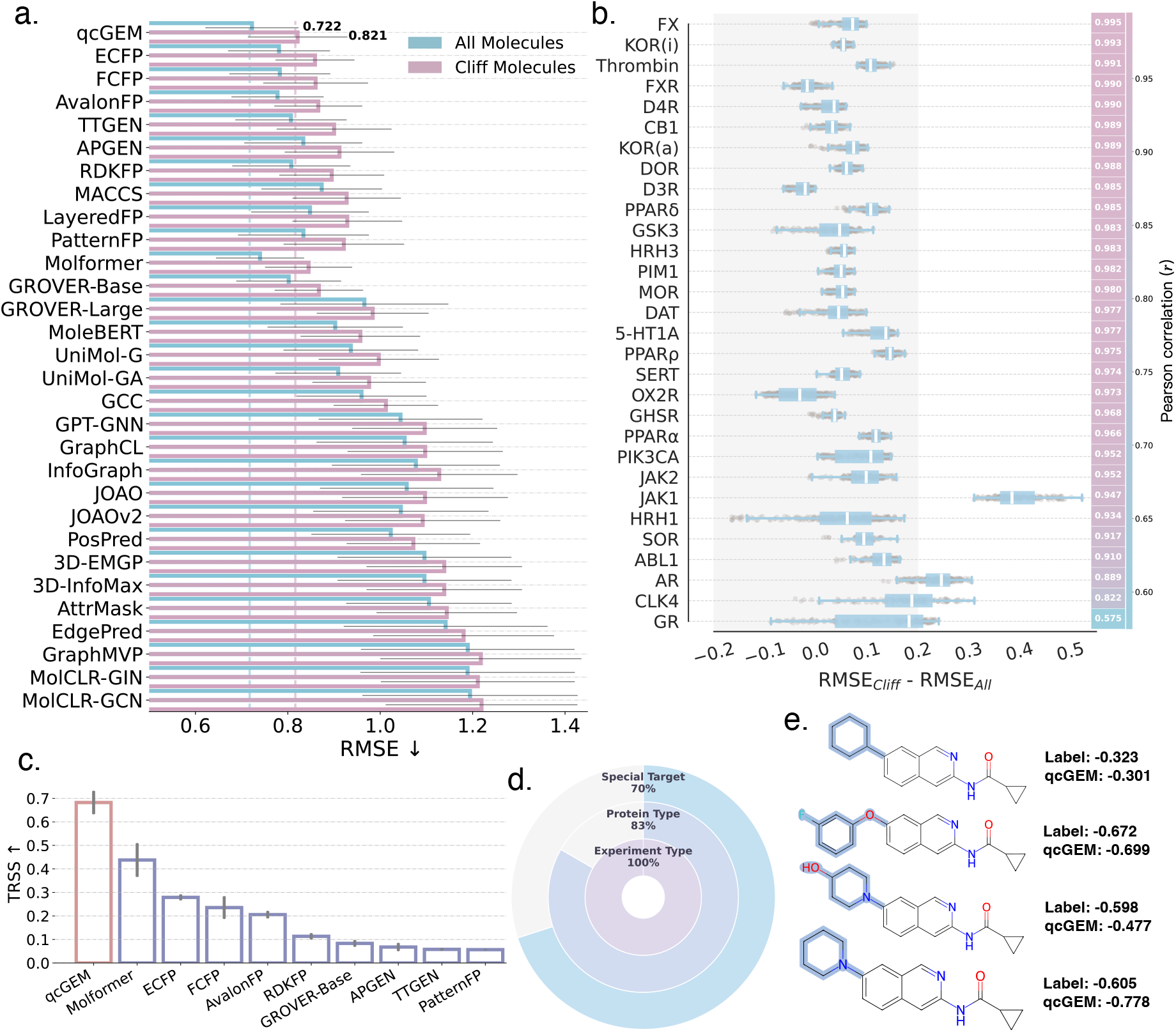
Performance of qcGEM in capturing subtle structural variations. **a)** Comparison of prediction errors for all molecules (blue) and activity cliff molecules (pink) across 30 molecular representation methods. The prediction error is evaluated by RMSE, with the arrow aside indicating desired direction. Vertical dash-dotted lines indicate the best-performing results. **b)** The box plot (left) shows the distribution of differences in prediction errors (using qcGEM) between activity cliff molecules and all molecules (*i.e*. RMSE_Cliff_ − RMSE_All_) across 30 protein targets, where box limits correspond to quantiles with whiskers extending to 1.5 inter-quartile range. The heatmap (right) illustrates the Pearson correlation coefficient *r* between inference values of activity cliff molecules and all molecules during the training of MLP module. **c)** The overall TRSS scores of the top methods. Here, TRSS scores are calculated with five random seeds across activity cliff molecules and all molecules, respectively, and their mean values and standard deviations are shown in the bar plot. **d)** The multi-layer pie chart illustrates proportions of qcGEM achieving the best ranking in 30 tasks under three task classification criteria: individual protein targets (outer), protein classes (middle), and experimental categories (inner). **e)** An example of the identification of activity cliff molecules for the protein kinase ABL1. For each ligand, the difference in chemical structure is highlighted in blue shadow, while the ground-truth binding affinity and prediction based on qcGEM are listed aside.

Following the MoleculeACE^51^ work, we evaluated the difference of prediction error between activity cliff molecules and all molecules to assess the generalizability of qcGEM on bioactivity prediction in the presence of activity cliff molecules. In most protein targets, prediction errors of the two sets of molecules only exhibit a minor deviation, falling between -0.2 and 0.2, laterally indicating good generalization (Figure 3b, left; Supplementary Figure 11a). Consistently, during the optimization of MLP parameters (see Figure 2a), models trained on activity cliff molecules and all molecules present a strong Pearson correlation coefficient for most protein targets, precluding biased training (Figure 3b, right; Supplementary Figure 11b). In addition, qcGEM also demonstrates consistent performance across different protein types (Supplementary Figure 12), achieving the leading position in most types. Furthermore, qcGEM effectively captures the minor structural modifications that lead to notable changes in bioactivity. In case studies on protein targets of various classes, despite the high similarity among candidate ligands, qcGEM successfully distinguishes them and yields reliable predictions on binding affinity (Figure 3e and Supplementary Figure 13).

These results demonstrate the capacity of qcGEM in effectively capturing the subtle structural variations within chemical compounds. The strong robustness and generalization across diverse application scenarios underscore its potential in high-throughput screening of active drug candidates. Detailed per dataset results could be found in Supplementary Tables 21-32.

### Benchmark on protein-ligand binding prediction with qcGEM

To comprehensively assess the representational capability of qcGEM, we examined it against the other methods on 11 protein-ligand interaction prediction tasks by integrating the molecular representation with the output of protein language model ESM-3^52^ (Figure 2a). The datasets^53–57^ cover 5 general protein-ligand interaction tasks (Supplementary Table 7) and 6 opioid-type tasks (Supplementary Table 8).

In the general protein-ligand interaction prediction, qcGEM achieves the best overall performance across both the 3 classification tasks and the 2 regression ones (Figure 4a). Specifically, it ranks first in one task and is consistently located within the top 5 in all the others (Figure 4c, upper; see Supplementary Figure 14a for detailed statistics). In cases that qcGEM fails to achieve the highest rank, absolute inferiority in the according metric is frequently small (Supplementary Table 33). In contrast to other methods that occasionally excel on a single task, qcGEM exhibits the highest TRSS score, demonstrating its robust and consistent performance across all evaluated tasks (Figure 4c, lower). Subsequently, we validated the predicted positive ligands by molecular docking. The resultant ligand-protein structures reveal plausible interaction patterns, indicating that qcGEM effectively facilitates the identification of protein-ligand binding modes (Figure 4e and Supplementary Figure 15).

**Figure 4.**
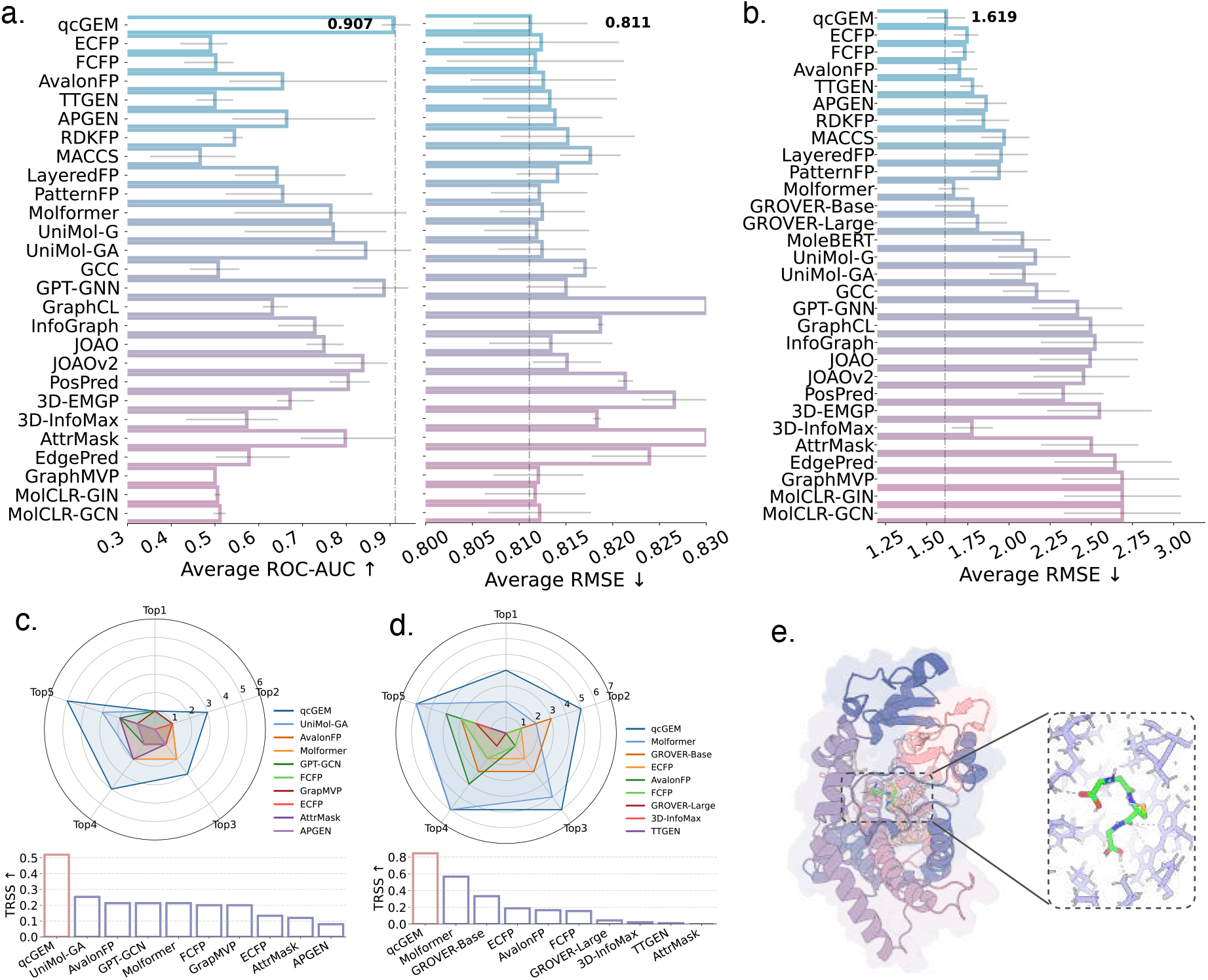
Performance of qcGEM on protein-ligand prediction. **a)** Performance comparison of qcGEM and other methods on the general protein-ligand interaction prediction tasks. ROC-AUC is shown for classification tasks (left), while RMSE is reported for regression tasks (right), with arrows indicating the desired directions. Vertical dash-dotted lines indicate the best-performing results. **b)** Performance comparison of qcGEM and other methods on the opioid-type molecule-receptor interaction prediction tasks. The prediction error is evaluated by RMSE, with the arrow indicating the desired direction. The vertical dash-dotted line indicates the best-performing result. **c)** The radar chart (upper) shows the occurring counts of each method in the top 1-5 rankings on the general protein-ligand interaction prediction tasks, while the bar plot (lower) presents the corresponding TRSS scores. Unlike previous evaluations (Figures 2e & 3c), the TRSS score is evaluated here only in one scenario, disallowing the calculation of standard deviation. **d)** Results of the opioid-type molecule-receptor interaction prediction tasks, presented in the same convention as in panel (**c**). **e)** The docking result of the predicted positive ligand for an exemplar protein target. The inset shows the atomic-level interaction details, with the ligand colored by elemental types and protein side-chains colored in purple.

We also evaluated qcGEM on opioid-receptor interaction prediction. Unlike previous tasks involving diverse interaction types, this setting focuses exclusively on interactions with 6 opioid receptor subtypes. Again, qcGEM achieves the best overall performance (Figure 4b) and demonstrates strong robustness in ranking (Figure 4d; see Supplementary Figure 14b for detailed statistics), which further validates its generalizability to domain-specific scenarios.

These results confirm that qcGEM can be effectively integrated with external models to deliver stable performance across a wide range of protein-ligand related tasks. In practice, the protein encoder (in Figure 2a) can be easily replaced with other high-quality backbones for seamless adaptation to task-specific applications. Detailed per dataset results could be found in Supplementary Tables 33-34.

### Scalable and robust performance of qcGEM-Hybrid

Previous evaluations have demonstrated that qcGEM abstracts essential molecular information into a compact form through pre-training and achieves strong performance across a broad spectrum of downstream tasks. However, the standard qcGEM relies on the full B3LYP-D3/def2-SV(P) pipeline for geometric optimization of candidate molecules, which is slow and thus hinders efficient feature derivation for molecules without pre-calculated quantum chemical annotations. We attempted to accelerate the computational process by leveraging the semiempirical GFN2-xTB^58^ model to achieve more rapid, albeit less accurate, geometric optimization, under the assumption that qcGEM has learned to handle suboptimal structures through the pre-training task of geometry denoising (Supplementary Figure 3). To this end, we proposed a hybrid variant qcGEM-Hybrid, by simply replacing the full B3LYP-D3/def2-SV(P) quantum chemistry calculation with B3LYP-D3/def2-SV(P)//GFN2-xTB while keeping the remainder of the pipeline unchanged.

In terms of quantum feature calculation, qcGEM-Hybrid offers substantial time savings over qcGEM (Figure 5a). Its advantage becomes more pronounced with the increase of molecular size, as evidenced by the reduction of power-law scaling exponent from 2.58 to 1.76. Specifically, for a large molecule of 100 atoms, the time consumption of quantum chemistry calculation on a single CPU core drops from 18 hours to 28 minutes, and will be further reduced to within 2 minutes if 16 CPU cores are employed (Figure 5b). Considering the nearly negligible time for model inference by the encoder, the overall computational efficiency of qcGEM-Hybrid is fully acceptable in practical embedding generation for unseen molecules.

**Figure 5.**
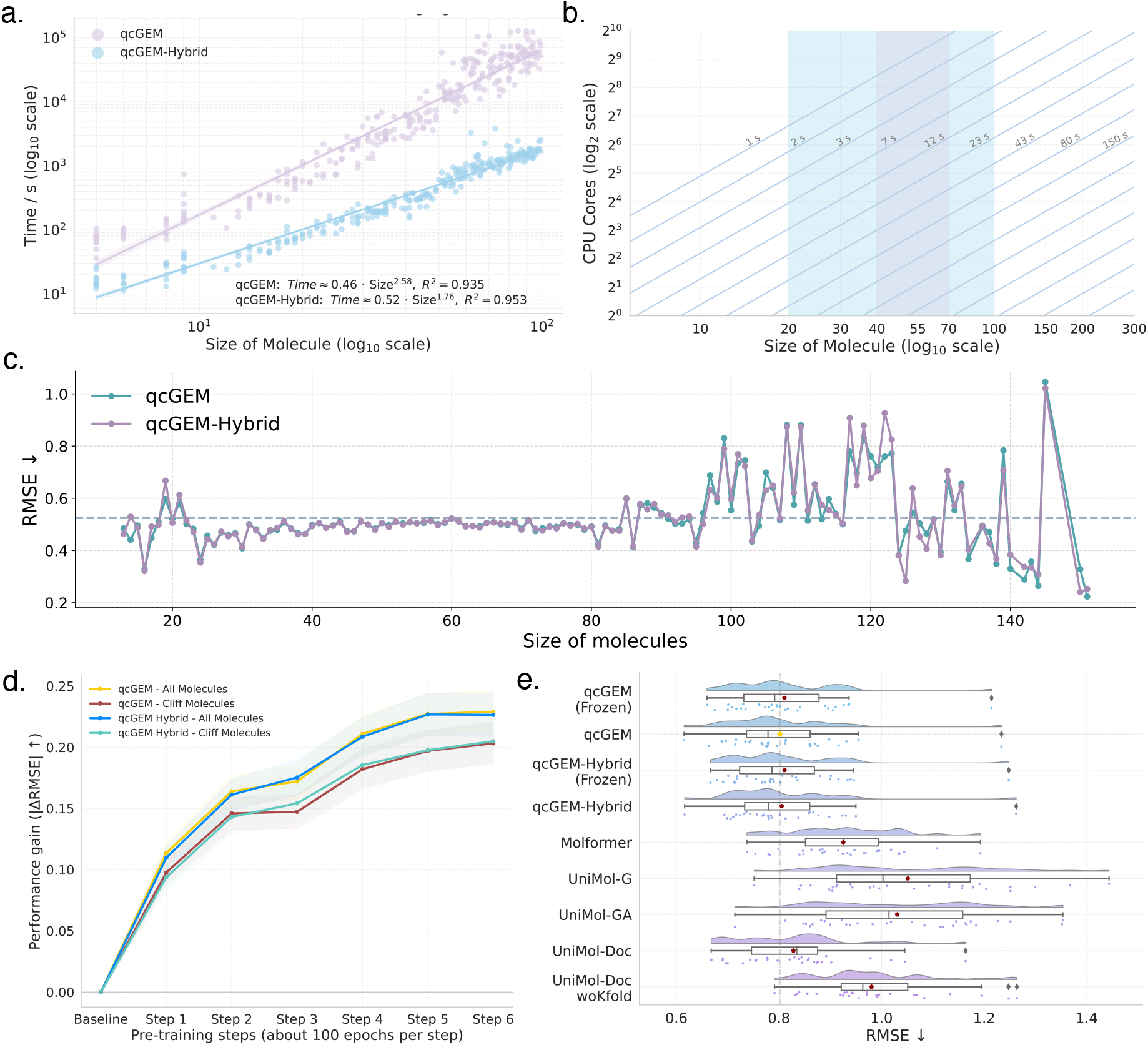
Evaluation of the efficiency and accuracy of qcGEM-Hybrid. **a)** Computation time vs. molecular size for qcGEM (purple) and qcGEM-Hybrid (blue) on the double logarithmic scale. The optimal parameters for the power-law fitting are shown in the legend, with *R*^2^ values indicated aside. **b)** Dependence of the time consumption by qcGEM-Hybrid on molecular size and CPU cores. The time cost is presented in the format of isolines, with the horizontal and vertical axes representing the molecular size and CPU core number, respectively, in the logarithmic scale. The shaded areas colored in blue and purple highlight ranges of molecular sizes for molecules of common interest and of high frequency in practical use, respectively. **c)** Prediction errors of qcGEM (green) and qcGEM-hybrid (purple) on molecules of different sizes in activity cliff prediction. **d)** Evaluation of qcGEM and qcGEM-Hybrid in activity cliff prediction based on various model checkpoints collected in the pre-training stage. The performance gain (in the vertical axis) is quantified by the reduction in prediction error (*i.e*. |ΔRMSE| ), when compared to the baseline (the 20th epoch). **e)** Comparison of the fixed (*i.e*. frozen) and fine-tuned qcGEM/qcGEM-Hybrid representations with various fine-tuned versions of Molformer and UniMol in activity cliff tasks. Here, Molformer was fine-tuned following the official guideline, while the fine-tuning of UniMol was attempted in a number of strategies, including global-token-based fine-tuning (UniMol-G), global-atom-based fine-tuning (UniMol-GA), fine-tuning according to the official documentation (UniMol-Doc), and fine-tuning without *K*-fold cross-validation (UniMol-Doc woKfold). In each box plot, the central bar indicates the median, the box limits correspond to the lower and upper quartiles, the whiskers extend to 1.5 inter-quartile range, and the star denotes the mean value. The vertical dash-dotted line indicates the best performance in terms of the mean value.

In activity cliff tasks, the encoder shows a consistent trend when using the two pipelines for quantum chemistry calculation, though the qcGEM-Hybrid variant exhibits a slightly higher variation in prediction error for larger molecules (Figure 5c). In addition, when evaluated on model checkpoints collected from various stages in the pre-training phase of the original qcGEM, qcGEM-Hybrid also attains similar gains of performance (Figure 5d). These observations demonstrate that the pre-trained qcGEM encoder is robust upon potential structural noises introduced by low-precision geometry optimization procedures like GFN2-xTB, further supporting our prior assumption.

Owing to its markedly enhanced efficiency, we evaluated the qcGEM-Hybrid embedding under the fine-tuning scenario against two state-of-the-art baselines (Molformer^18^ and UniMol^19^) that also support encoder fine-tuning (Figure 5e). In activity cliff tasks, the fixed qcGEM-Hybrid with frozen parameters has already surpassed fine-tuned Molformer and UniMol, reaching a comparable performance to qcGEM. Moreover, with only minimal fine-tuning on the encoder parameters, the performance of qcGEM/qcGEM-Hybrid further improves, underscoring the flexible application modes of our representation in downstream tasks. Detailed information of fine-tuning experiments could be found in Supplementary Tables 35-36.

Overall, qcGEM-Hybrid achieves a favorable trade-off between efficiency and accuracy, ensuring both scalability and reliability for downstream applications. For applications requiring high-precision predictions, qcGEM is recommended to generate high-quality representations for repeated reuse. In contrast, qcGEM-Hybrid provides a more efficient and reliable alternative in the large-scale screening and/or fine-tuning scenarios. By supporting representation reuse and rapid inference, qcGEM and its hybrid variant offer a practical and scalable solution as an efficient large-scale molecular filter.

### Interpretability and quantum chemistry awareness of qcGEM

After the previous systematic evaluations, in this section, we further examined the latent representation space of qcGEM to assess its interpretability.

We employed t-distributed stochastic neighbor embedding (t-SNE)^59^ to visualize the landscape of the qcGEM representation space (Figure 6a). Although molecular weight predominantly determines the global distribution, molecules of high similarities in chemical formulas tend to cluster locally, with observable gradual transitions between adjacent clusters. The atomic-level representations also show notable clustering: atoms of different types occupy distinct regions, and atoms of the same kind are further divisable based on their chemical environments (Figure 6b).

**Figure 6.**
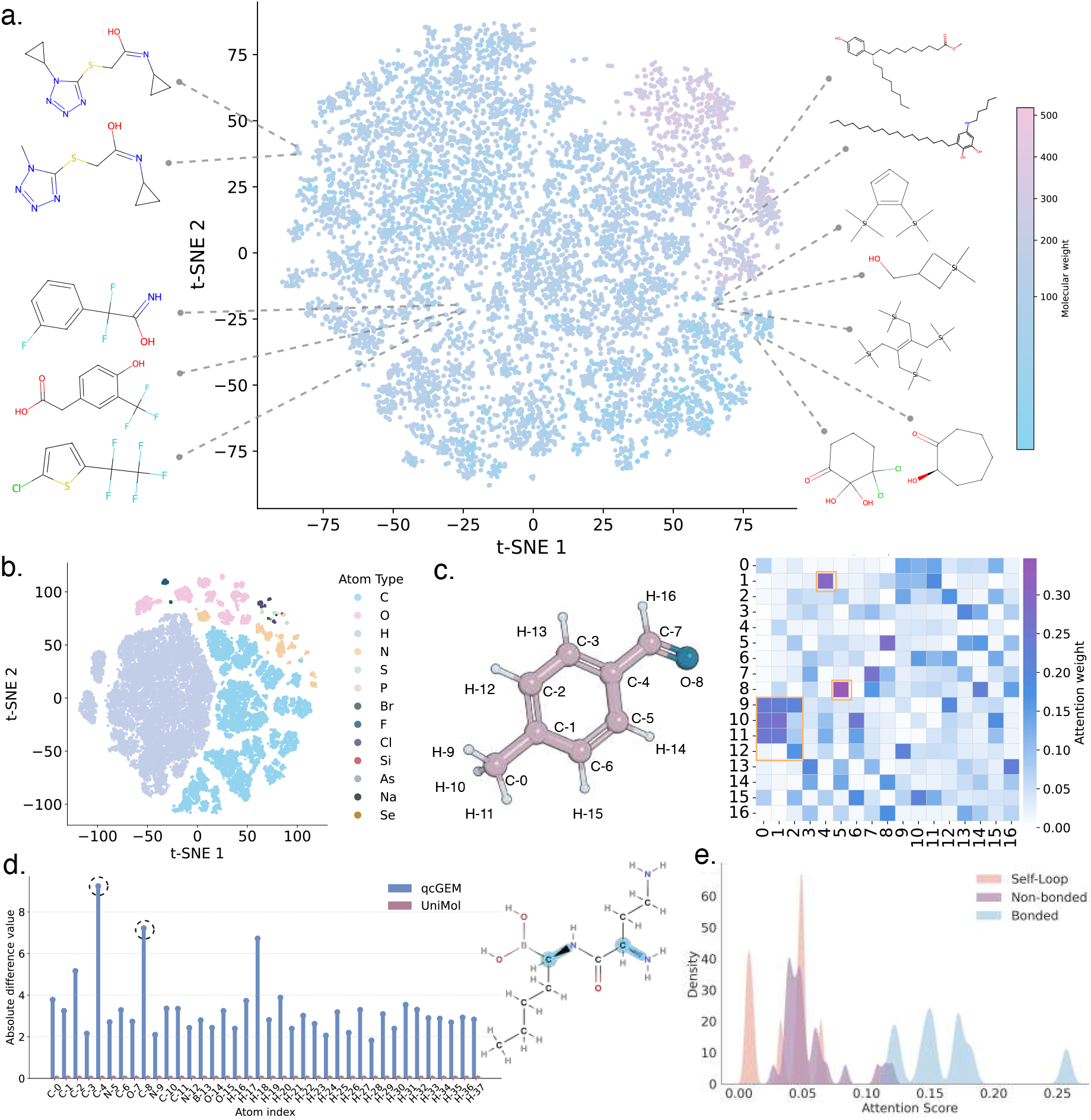
Analysis of the embedding of qcGEM. **a)** t-SNE visualization of qcGEM global-level representations, colored by molecular weight. Representative molecules from distinct regions highlight meaningful geometric clustering in the learned embedding space. **b)** t-SNE visualization of atom-level embeddings learned by qcGEM, colored by atom type, showing clear clustering by chemical element. **c)** Showcase of edge-level embeddings on an exemplar molecule (left). In the attention map (right), non-local interactions captured by qcGEM are highlighted by red boxes. **d)** Comparison of atom-level embedding differences between two stereoisomeric structures. In the right panel, atoms highlighted by blue shadow represent chiral centers. In the left panel, difference between stereoisomers is shown for qcGEM (blue) and UniMol (red) embeddings in the atom-wise manner, with chiral centers (*i.e*. C4 and C8) indicated by dashed circles. **e)** Distribution of attention scores for self-loops as well as bonded and non-bonded atom pairs.

Indeed, qcGEM has learned to encode essential interatomic communications into the graph-based embedding, from which all quantum descriptors could be recovered with low error levels (Supplementary Figure 4). Here, we present a case study on the edge-level information of qcGEM representation for 1-naphthaldehyde (Figure 6c), a commonly used lead compound in drug discovery and fluorescence probe design. Clearly, qcGEM captures the characteristic non-local interatomic interactions in this conjugate molecular system, for instance, the evident communication between C5 and O8 in the attention map. On the other hand, systematic analysis on the distribution of all cross-atom attention scores further reflects a physically meaningful hierarchy (Figure 6e): bonded atoms receive the highest attention scores, consistent with their role in defining molecular connectivity; non-bonded pairs also receive sufficiently high attention, highlighting the model’s sensitivity to through-space non-covalent interactions such as dispersion and *π*-*π* stacking; self-loops, in contrast, receive the least attention, reflecting a trade-off between preserving atomic identity and message updating.

Moreover, qcGEM representation preserves complete information of the molecular geometry, enabling accurate and successful three-dimensional structure reconstruction (Supplementary Figure 5). Bortezomib is an FDA-approved anticancer drug based on boronic acid, whose therapeutic effect critically depends on the configuration of its two chiral centers (Figure 6, right). In comparison with UniMol^19^ (a baseline trained with large-scale molecular structures), qcGEM is evidently advantageous in accurately discriminating the two stereoisomers of this compound (Figure 6d, left), which demonstrates its sensitivity to stereochemical features that are essential for molecular recognition.

Together with the physics-inspired model design, findings in this section demonstrate that qcGEM not only achieves strong predictive performance but also provides interpretable molecular representations.

## Discussion

In this study, we propose qcGEM, a quantum-chemistry-aware graph-based embedding of molecules, based on the localized quantum descriptors and the most stable molecular configurations, to address the shortcomings of nowadays molecular representations. Its physics-inspired architecture and rationally designed information flow enable efficient training, comprehensive encoding and strong interpretability. Evaluation results demonstrate the robustness and generalizability of qcGEM across 30 molecular property tasks, 30 activity cliff tasks, 5 protein-ligand interaction tasks and 6 opioid-specific tasks. The consistent superiority over 29 tested baseline methods underscores its potential for filtering promising drug candidates. qcGEM further exhibits consistent physical relevance and interpretability across multiple levels. The complete geometry encoding enables accurate discrimination of chiral enantiomers, reflecting sensitivity to subtle yet critical molecular variations. Furthermore, the qcGEM-Hybrid variant supports fast and reusable inference with robust accuracy, enabling scalable deployment in large-scale screening.

According to the Born-Oppenheimer approximation, motions of nuclei and electrons are roughly separable due to their markedly different time scales, and nuclei are conceived to move slowly within the electronic potential energy surface. Hence, in a graph-based molecular representation, nuclei are naturally modeled as nodes, with edges corresponding to the pairwise interactions between nuclei, which, in principle, should be quantified by electronic wave functions. However, in mainstream molecular representations, edges are frequently embedded from empirical chemical bonds (*e.g*., generated by RDKit^60^ or Open Babel^61^), a scheme that inherently overlooks the diffusive electronic orbitals spanning over more than two atomic centers, such as the *π* orbitals in conjugate molecular systems. The neglect of such non-local quantum effects will inevitably introduce systemic errors, which can hardly be rectified by simply amplifying the scale of data and labels as in many popular pre-training frameworks. To remedy this deficit, we train qcGEM by incorporating several localized quantum descriptors (*e.g*., natural bond orbitals, electron localization function, etc.) as well as the three-dimensional molecular structure. By this means, this physics-inspired framework effectively treats nuclei as well as inner-shell and lone-pair electrons as nodes, while considers the other valence-shell electronic orbitals as edges, agreeing with chemical intuition. The evident advantage of our representation over traditional ones in the large number of downstream tasks further reinforces the necessity of considering non-local (*i.e*. beyond empirical bonds) atomic interactions in molecular representation learning.

Despite the advantages, there remain a few limitations in the current qcGEM framework. First, incorporating additional level information like coarse-grained functional groups or molecular fragments may further enrich the representation with a new dimension of granularity, improving its chemical interpretability as well as practical applicability to a broader range of downstream tasks. Furthermore, the study of qcGEM-Hybrid suggests that while quantum chemical information is essential, low precision computation often suffices to achieve excellent performance. This is partly because overly precise signals tend to attenuate as they propagate through deep network layers, limiting their constraining ability. Moreover, downstream tasks often depend more on the distributional characteristics of representations than on their exact values. Therefore, future work will explore contrastive learning strategies to generate low-precision quantum chemical features for the further improvement of computational efficiency. Finally, although this study emphasizes the use of a relatively small pre-training dataset, further performance gains may be achieved by scaling up data volume and model capacity.

## Methods

### Molecular graph construction

qcGEM abstracts each molecule as a complete graph with multi-scales: ℳ = (𝒢, ℛ, 𝒱, ℰ ), where *G* stands for the gloabl state (*i.e*. molecular properties), 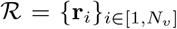 refers to the coordinates of atomic nuclei in the initial geometry (*i.e*. molecular structure), 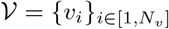 denotes the set of nodes (*i.e*. atomic centers), 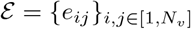 describes the set of edges (*i.e*. interatomic interactions), and *N*_*v*_ is the number of atoms.

The global state is initialized with a learnable token, while the nodes and edges are initialized using localized quantum chemical features. Quantum chemical features could be obtained from the qcMol database or computed from scratch using GFN2-xTB^58^ for geometry optimization and ORCA^62^ at the level of B3LYP-D3/def2-SV(P) for quantum chemistry calculation, followed by wave function analysis with NBO^63^ and Multiwfn^42^. The complete feature details of nodes and edges are provided in Supplementary Tables 1 and 2, respectively. We denote the initial features of global molecular state 𝒢, coordinates *R*, node *V* and edge *E* as *g, r*_*i*_, *v*_*i*_ and *e*_*ij*_, respectively. Notably, in the final representation graph *M*_rep_, *R* is incorporated into *V* in the rotranslation-invariant manner after encoding.

### Model architecture

The qcGEM model adopts an autoencoder framework to extract the representation of a compound (Algorithm 1). After feature pre-processing (Algorithm 2), the encoder employs 16 repeated encoding blocks to encode molecular graph *ℳ* as the representation *ℳ*_rep_ (Algorithm 3). This representation is then passed to the decoder (Algorithm 4), where it is used to reconstruct the molecular geometry, quantum chemical information and global state (Algorithm 5).

At the beginning of the encoding process, the geometry module extracts geometric features from the Cartesian co-ordinates ℛ, which are then used as positional embeddings *PE*_node_ and *PE*_edge_ for nodes and edges, respectively (Algorithm 6):

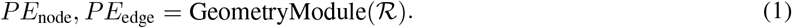

In every encoding block, the node features *v*_*i*_ are firstly updated to obtain the node embeddings *h*_*v*_*i* via a self-attention mechanism applied over all node pairs (including self-loops) based on both node features *v*_*i*_ and pairwise edge information *e*_*ij*_ (Algorithm 7). In this process, the geometric features *PE*_node_ and *PE*_edge_ are also incorporated through attention bias and feature concatenation:

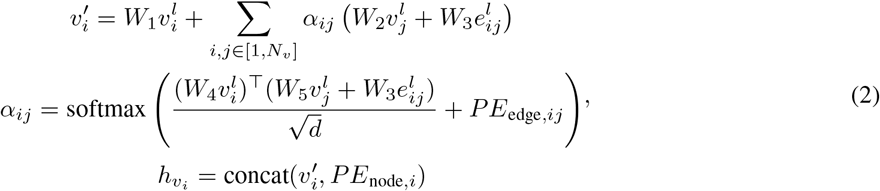

where the superscript *l* denotes the index of encoding block, *W* represents the trainable matrices for linear projection, *d* is the feature dimension of node *v, α*_*ij*_ stands for the attention score between *v*_*i*_ and *v*_*j*_, while softmax(*·*) and concat(*·*) refer to the softmax function and concatenation operator, respectively.

Subsequently, the GLI unit is applied to facilitate information exchange between global and local representations (Algorithm 8). In this module, node and edge representations are read out to form local representations, concatenated with the global token, and further processed to enable information fusion. The output is then broadcast to node and edge to update their representations:

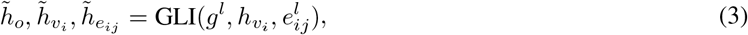

where 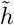 denotes the hidden states of output *o*, node *v* and edge *e*, respectively.

After sufficient information exchange, the node *v* and edge *e* are updated based on the contextual information learned through the GLI module (Algorithm 9). The global token is refined with aggregated local information to produce a meaningful global embedding (Algorithm 10):

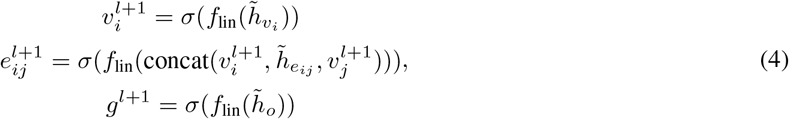

where *f*_lin_ denotes linear layers of the module, and *σ* represents the non-linear activation function.

To construct a hierarchical molecular representation and promote stable gradient propagation, we aggregate the output of encoding blocks from a pre-defined number (*L*) of layers, each with different semantic information. Finally, the input molecular graph ℳ is progressively transformed into a compact, quantum-chemistry-aware representation ℳ _rep_ comprising global, node and edge levels,

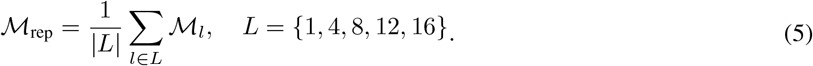

In the decoding stage, the folding optimizer, QC recovery unit and global predictor independently decode the graph molecular representation to recover the molecular geometry, quantum chemical features and global molecular state, respectively:

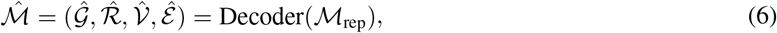

where Decoder(*·*) denotes the decoder function that consists of the folding optimizer, QC recovery unit and global predictor, and symbols with a hat 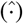 denote reconstructed results.

#### Geometry module

To mitigate geometry information loss observed in previous methods^64,65^, we referred to and modified the ComENet^66^ framework to enhance extraction efficiency while preserving compatibility with molecules processed by qcGEM (Algorithm 6). We split the entire geometry information into multiple local, complete geometries, utilizing (*d, θ, ϕ, τ* ) in the spherical coordinate system to achieve local completeness. By incorporating complete local geometries as positional embeddings into each qcGEM encoder block, the representation preserves a complete memory of the geometry. Furthermore, this operator could mitigate the over-smoothing problem in GNNs^67,68^. The detailed formulation is provided in Supplementary Information 1.2.

#### GLI Unit

The GLI unit is designed for information exchange between *g, v*_*i*_ and *e*_*ij*_. It is a bidirectional unit that facilitates both the bottom-up flow of local information to global representations and the top-down refinement from global to local features (Algorithm 8). In the local-to-global information flow, node and edge embeddings are aggregated via mean pooling and then used to update the global token. In the global-to-local phase, the global embedding is propagated back to local nodes, providing contextual guidance for refining the local representation. The concatenated information is further fused through the self-attention mechanism. This strategy provides a multi-level representation, facilitating adaptation to diverse tasks, while simultaneously enhancing the readout training of node and edge features to reduce global feature entanglement^33^. The detailed formulation is provided in Supplementary Information 1.3.

#### Folding optimizer

The folding optimizer is designed with a dual-flow architecture that incorporates both 1D and 2D geometry information (Supplementary Figure 1a; Algorithm 12). An initial geometry is generated from 1D features under the guidance of 2D structural information. This geometry is first evaluated through pairwise distance constraints in the 2D channel and then fed back into the 1D channel. The model then uses 1D features to predict the difference between initial and ground-truth geometries. Through iterative refinement, the folding optimizer drives the predicted structure toward a more stable and energetically favorable conformation. The detailed formulation is provided in Supplementary Information 1.4.

### Loss design

The total loss (Algorithm 18) is composed of three components: global prediction loss (Algorithm 19), node and edge feature recovery losses (Algorithms 20 and 21), and geometry loss (Algorithm 22). Non-geometry-related tasks are supervised under the *L*_1_ and cross-entropy losses. In contrast, for geometry-specific tasks, we introduce a tailored geometric constraining loss to enhance learning stability and convergence,

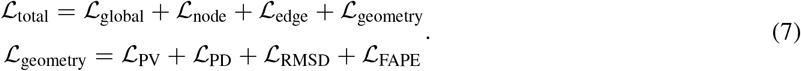

The geometric constraint loss contains PV loss, PD loss, RMSD loss and FAPE loss for model learning. The PV loss (pairwise vector difference), PD loss (pairwise distance difference) and RMSD loss (root mean square deviation) are used to constrain the overall molecular geometry. We specifically propose a small-molecule-oriented FAPE^45^ (*i.e*. frame aligned point error) loss to impose more effective structural constraints (Algorithm 23).

#### FAPE loss

To better constrain local geometric details, we define rigid triangles formed by heavy atoms as local frames (Supplementary Figure 1b). A local coordinate system is constructed for each frame to facilitate the calculation of geometric deviations. By evaluating geometric errors across all frames within a molecule, the model enforces constraining from multiple local perspectives. The FAPE loss serves as a strong geometric constraint on local details and is used when the convergence is challenging. The detailed formulation is provided in Supplementary Information 1.6.

### Pre-training strategy

The qcGEM model utilizes geometry denoising, node-edge feature recovery and correction, and global condensation for pre-training. For the denoising task, random noise in the range of -1 Å to 1 Å is added to the native coordinates, and the model is trained to refine the perturbed geometry toward the ground truth. For node-edge feature recovery and correction, input features are either randomly masked with a learnable token or mutated by randomly sampled features within the same training batch. The model learns to identify and differentiate between true and erroneous features and subsequently recover the correct one. We experimented with various masking and replacement ratios, and ultimately trained the model with a 30% masking ratio, a 30% replacement ratio and a 10% reversion ratio (Algorithm 2). For the global condensation task, the model condenses the global state from nodes and edges to update the token-initialized global features, which are subsequently used to predict target descriptors.

The overall pre-training process involves multiple stages (Supplementary Figure 2), with details listed in Supplementary Table 3.

### Evaluation metrics

For compound properties, we use RMSE to evaluate regression tasks and adopt the area under the ROC curve (ROC-AUC) for classification tasks. To further assess the robustness of different methods, we propose the TRSS score as a novel metric:

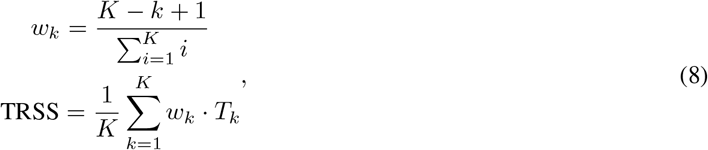

where *K* denotes the maximum in the top-ranking evaluation and is set to 5 by default, and *T*_*k*_ refers to the occurring count of a method within the top *k* ranks. TRSS quantifies the robustness of a method by measuring how frequently it ranks within the top *K* across multiple tasks. A linearly decaying weight *w*_*k*_ is assigned to the rank position *k* to favor higher rankings. Unlike metrics that penalize occasional poor performance, TRSS rewards consistent top-tier performance, thus mitigating the influence of outliers. This design makes it particularly suitable for real-world evaluation scenarios, where stability and reliability across diverse datasets are often more critical than occasional peak performance on a limited number of subsets.

### More computational details

We performed quantum chemical calculation with B3LYP-D3/def2-SV(P)//GFN2-xTB by ORCA 5.0.3^62,69–75^ and GFN2-xTB 6.5.1^58^. The post-analysis of wave functions was done by Multiwfn 3.8^42^ and NBO 7.0^63^. Molecules were initially processed by RDKit 2022.09.5^60^ and Open Babel 3.1.1^61^. The qcGEM model was implemented with PyTorch^76^ version 2.0.0 and PyTorch Geometric^77^ version 2.5.2 with CUDA version 12.0 and Python 3.8.19. Adam optimizer with learning rate ranging between 3*e*^*−*4^ and 3*e*^*−*5^ was used to optimize the model (Supplementary Table 3). The model comprises approximately 37 million parameters, of which about 36 million belong to the encoder. Learning rate of the MLP probe network was set to 1*e*^*−*4^ with a weight decay of 0.01.

## Data availability

### Pre-training dataset

We used the subset of PubChem quantum chemical annotation from the qcMol^44^ database as our pre-training materials.

### Evaluation dataset

qcGEM was evaluated on the MoleculeNet^47^ dataset and the ADMET subset from TDC^48^ dataset for compound properties tasks, on the MoleculeACE^51^ dataset for activity cliff estimation tasks, and on the BindingDB^53^, Human^54^, C. elegans^54^, KIBA^55^, Daivs^56^ and Opioids^57^ datasets for multi-level protein-ligand interaction tasks.

All relevant data supporting the key findings of this study are provided in the article and its Supplementary Information, and are also available for download from the Zenodo repository at https://doi.org/xxx.

## Code availability

The source code of qcGEM can be accessed from the GitHub repository at https://github.com/GHUSER-haoyu/qcGEM, with model parameters obtainable from the Zenodo repository at https://doi.org/10.5281/zenodo.17364257.

Pre-calculated qcGEM embeddings could be freely downloaded from the website of qcMol database at https://structpred.life.tsinghua.edu.cn/qcmol/. The online server of qcGEM provides embedding generation service for submitted molecules at https://structpred.life.tsinghua.edu.cn/server_qcGEM.html.

## Author contribution

H. G. proposed the theory and concept. H. W. and H. G. designed the methodology. H. W. accomplished coding, model training, performance evaluation, and data analysis. H. G. supervised the whole process. H. W. and H. G. wrote the manuscript. Both authors agreed with the final manuscript.

## Acknowledgements

This work has been supported by the Ministry of Science and Technology of China (#2023YFF1204400 to H. G.), by the National Natural Science Foundation of China (#32171243 to H. G.), and by the Beijing Frontier Research Center for Biological Structure.

## Supplementary Information

### 1 Supplementary Methods

#### 1.1 Model architecture

**Supplementary Figure 1.**
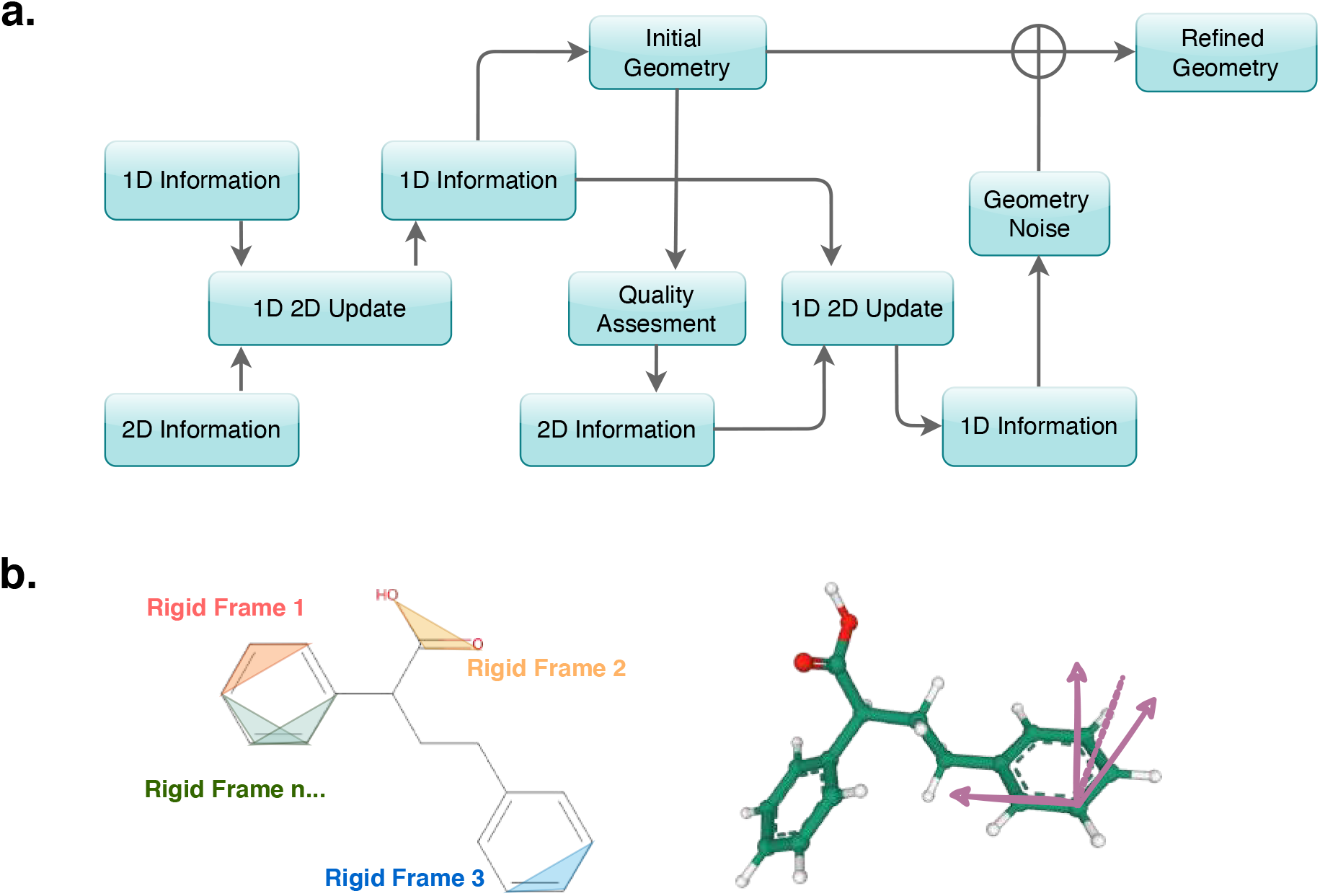
Detail of qcGEM Model Design. **a)** The folding optimizer produces an initial and refined geometry, supporting iterative geometry refinement. **b)** The FAPE loss specifically tailored for small molecules enhances geometric supervision by constructing chemically reasonable local coordinate systems from multiple perspectives.

##### 1.2 Geometry module

Let 𝒜𝒯 = {*at*_*i*_} denote the atom types and let ℛ = {**r**_*i*_} denote the set of atomic coordinates in a molecule, where **r**_*i*_ *∈* ℝ ^3^ represents the position of the *i*-th atom. The set of edges ℰ is defined as:

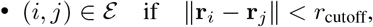

where *i* and *j* refer to indices of atomic nodes, *r*_cutoff_ denotes a pre-defined hyper-parameter that determines the spatial extent of local geometric neighborhoods.

To define the geometric feature for an edge, *e.g*., starting from node *i* and ending at node *j*, a local coordinate system is first established by finding the nearest and the second nearest neighboring atoms of node *i, n*_0_(*i*) and *n*_1_(*i*), based on Euclidean distances in the Cartesian space. Using these reference atoms, we derive four rotranslationally invariant variables:

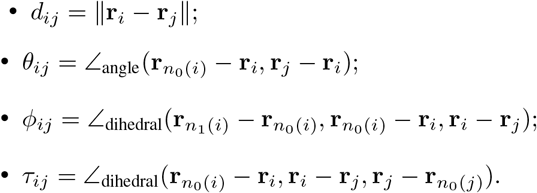

The set of variables (*d*_*ij*_, *θ*_*ij*_, *ϕ*_*ij*_, *τ*_*ij*_) are then used to compute relative geometric features in the local space,

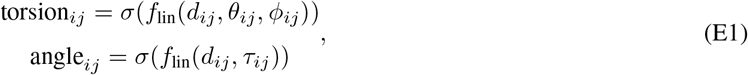

where *f*_lin_(*·*) denotes linear layers and *σ*(*·*) represents the non-linear activation function. The local geometric features are first transformed by non-linear operation,

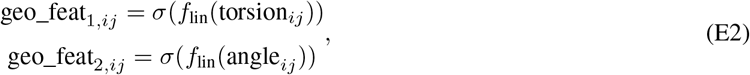

and then act as edge weights to integrate atom type information from neighborhood 𝒩 in the graph,

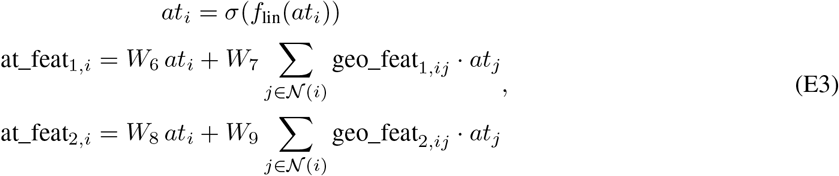

where *W* represents the trainable matrices for linear projection.

We subsequently perform a nonlinear transformation and aggregate these local information to derive spatial positional embeddings *PE*_node_ and *PE*_edge_ for nodes and edges, respectively,

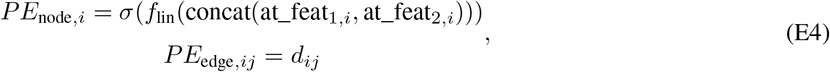

where concat(*·*) refers to the concatenation operator.

##### 1.3 GLI unit

For interactions between the local information and the global state, we employ GNN readouts combined with PyTorch’s broadcasting operations to facilitate message propagation:

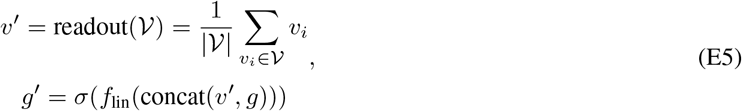

where *g* and *v*_*i*_ represent the input feature of global state and nodes, respectively.

A multi-head attention mechanism is further utilized to enable deep fusion of global and local information (Figure 1c):

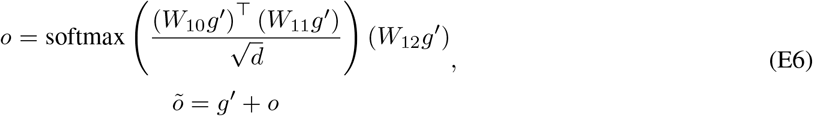

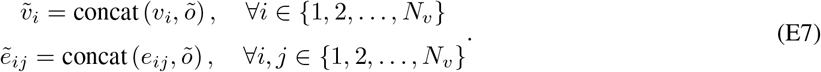

Here, *v*_*i*_ and *e*_*ij*_ represent the original inputs of nodes and edges, while 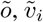 and 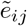 are the final outputs for the global state, nodes and edges, respectively. *W* represents the trainable matrices for linear projection, *d* is the dimension of key vector, and *N*_*v*_ denotes the number of atoms.

##### 1.4 Folding optimizer

First, we apply a nonlinear transformation to the global representation *g*_rep_, the node representation *n*_rep,*i*_, and the edge representation *e*_rep,*ij*_:

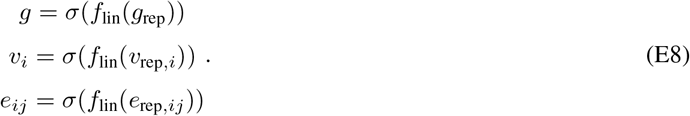

Here, *g, n*_*i*_ and *e*_*ij*_ are updated intermediate values of the global state, node and edge, respectively.

The node-edge exchange (NEE) module (see **the following section** for details) then aggregates node-edge information to produce updated representations. Using the updated node states, the decoder generates an initial structural prediction 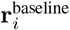 as a baseline:

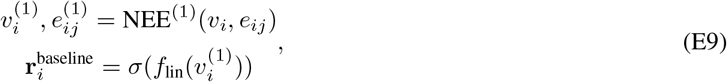

where the superscript represents the order of the NEE module.

This baseline is discretized via binning on the distance *d*_*ij*_ to estimate pairwise distance ranges, which are fed back as *b*_*ij*_ to edges for refinement:

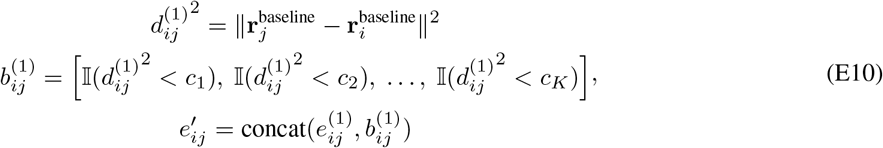

where (*c*_1_, *c*_2_, … , *c*_*K*_) is series of bin ranges, and I(*·*) is the indicator function, which outputs 1 if the condition is satisfied and 0 otherwise.

With the refined node and edge features, the model predicts the current structural noise *δ***r**_*i*_ to correct the baseline 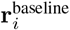, yielding the final structure 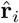:

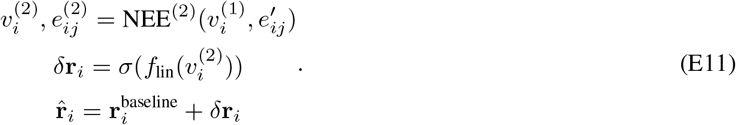

##### 1.5 NEE module

In the three-step process of updating node *v*_*i*_ and edge *e*_*ij*_ representations, the model first computes pairwise node differences by comparing the features of node pairs. Specifically, vector differences Δ_*ij*_, Euclidean distance squares 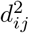, and cosine similarity cos *θ*_*ij*_ are calculated to construct node difference features *v*_diff_:

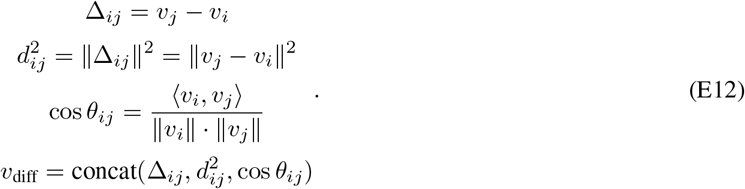

In the second step, these difference features are concatenated with the original node attributes *v*_*i*_ and *v*_*j*_, which are then used to update the corresponding edge features *e*_*ij*_:

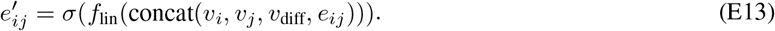

Finally, the updated edge features 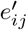 are aggregated as messages *m*_*i*_, which are integrated back into the node representations 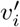, enabling bidirectional information exchange between nodes and edges:

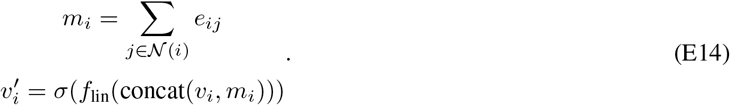

##### 1.6 Geometry loss design

To more effectively constrain molecular structures and facilitate rapid convergence, we introduce a set of loss functions targeting different aspects of geometric consistency. These include:

distance error loss

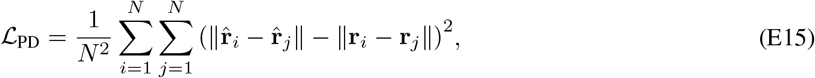

which penalizes deviations in inter-atom distances;

root mean square deviation (RMSD) loss

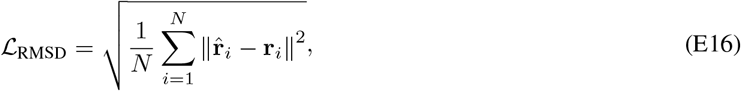

which evaluates global structural differences;

and vector difference loss

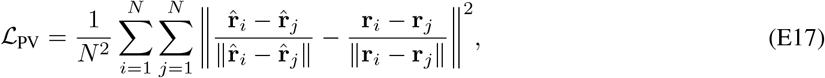

which jointly constrains both distance and directional deviations.

Here, the 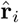 and **r**_*i*_ represent the ground-truth and predicted coordinates of atom *i*, respectively.

These losses serve to enforce the global structural integrity of molecular configurations. In addition, qcGEM incorporates the Frame Aligned Point Error (FAPE) loss

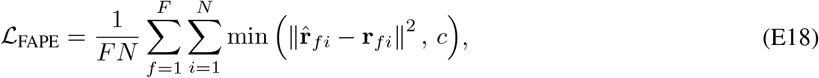

to provide localized structural supervision. Here, **r**_*fi*_ corresponds to the relative coordinate of atom *i* in the *f* -th local frame, and *c* is the cutoff, which is set as 10 Å by default. Our adapted version of FAPE is particularly effective in correcting local geometric distortions, thereby enhancing the fidelity of small-molecule reconstructions.

### 2 Supplementary Information for Model Pre-training

#### 2.1 Model features

**Supplementary Table 1.** Node features of qcGEM.

| Type | Description |
| --- | --- |
| Geometry | Cartesian atomic positions ($x, y, z$) defining the molecular geometry. |
| Atomic Number | Atomic number is the proton count that uniquely identifies an element. |
| Node Degree | Node degree is the count of bonds an atom forms with other atoms. |
| Formal Charge | Formal charge is the net atomic charge assigned from electron counting rules. |
| Linked Hydrogens | Linked hydrogens is the count of H atoms attached to an atom. |
| Radical Electron | Radical electron is the count of unpaired electrons on an atom. |
| Charge of NPA | Net atomic charge assigned by NPA. |
| Core of NPA | Number of electrons assigned to core (inner-shell) orbitals. |
| Valence of NPA | Number of electrons assigned to valence orbitals (involved in bonding). |
| Rydberg of NPA | NPA Rydberg population represents electrons assigned to diffuse, high-energy orbitals beyond the valence shell. |
| LI feature | LI quantifies the number of localized electrons associated with an atom, indicating electron localization. |
| ADCH charge | ADCH charge is a dipole-corrected variant of Hirshfeld atomic charges that better reproduces molecular dipole moments. |
| NAO feature | NAOs are orthogonalized atomic orbitals used as the localized basis in NBO analysis. |
| NBO-LP feature | Occupancy of a nonbonding localized orbital associated with a single atom (*i.e.* lone electron pair). |

**Abbreviations:**
NPA: Natural Population Analysis
LI: Localization Index
ADCH: Atomic Dipole Corrected Hirshfeld
NAO: Natural Atomic Orbital
NBO: Natural Bond Orbital
LP: Lone Pair

**Supplementary Table 2.**
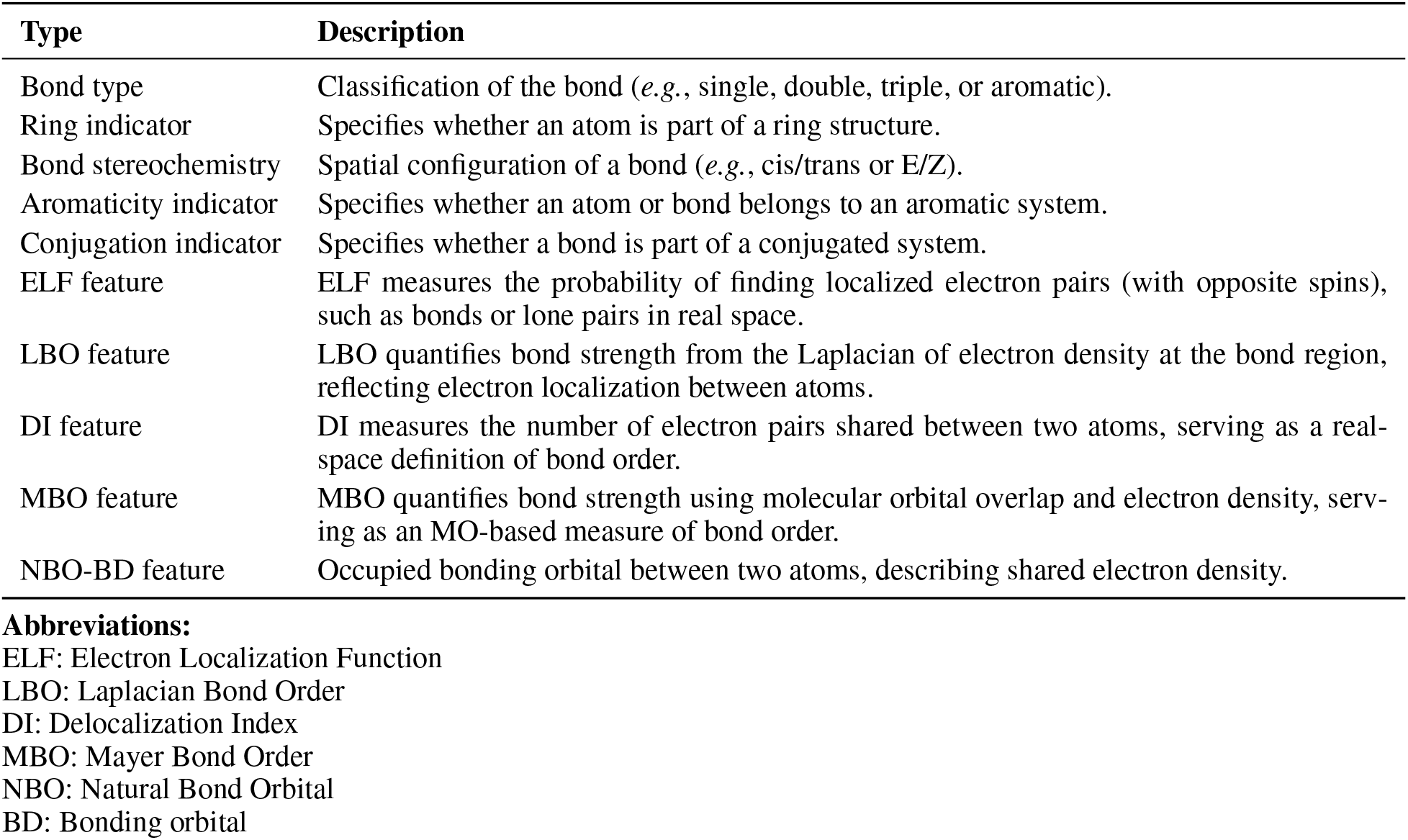
Edge features of qcGEM.

| Type | Description |
| --- | --- |
| Bond type | Classification of the bond (*e.g.*, single, double, triple, or aromatic). |
| Ring indicator | Specifies whether an atom is part of a ring structure. |
| Bond stereochemistry | Spatial configuration of a bond (*e.g.*, cis/trans or E/Z). |
| Aromaticity indicator | Specifies whether an atom or bond belongs to an aromatic system. |
| Conjugation indicator | Specifies whether a bond is part of a conjugated system. |
| ELF feature | ELF measures the probability of finding localized electron pairs (with opposite spins), such as bonds or lone pairs in real space. |
| LBO feature | LBO quantifies bond strength from the Laplacian of electron density at the bond region, reflecting electron localization between atoms. |
| DI feature | DI measures the number of electron pairs shared between two atoms, serving as a real-space definition of bond order. |
| MBO feature | MBO quantifies bond strength using molecular orbital overlap and electron density, serving as an MO-based measure of bond order. |
| NBO-BD feature | Occupied bonding orbital between two atoms, describing shared electron density. |

**Abbreviations:**
ELF: Electron Localization Function
LBO: Laplacian Bond Order
DI: Delocalization Index
MBO: Mayer Bond Order
NBO: Natural Bond Orbital
BD: Bonding orbital

#### 2.2 Pre-training process

**Supplementary Figure 2.**
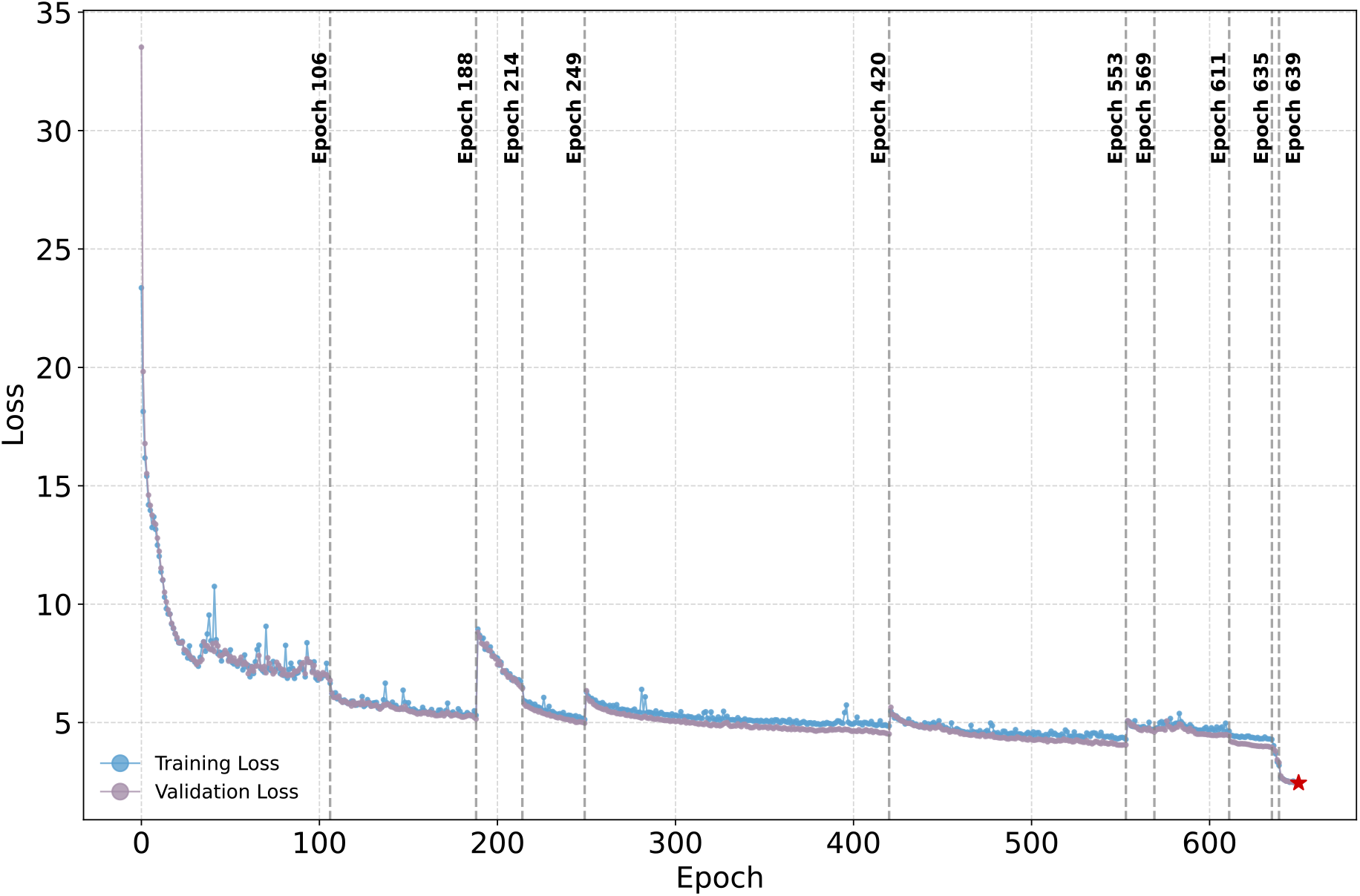
Pre-training loss curves across different stages. The model underwent 11 stages of manual fine-tuning, during which training loss consistently declined, resulting in enhanced downstream performance. Gray dashed lines mark the checkpoints connecting consecutive stages. The model selected for representation evaluation (Epoch 650) is labeled with a red star.

**Supplementary Table 3.**
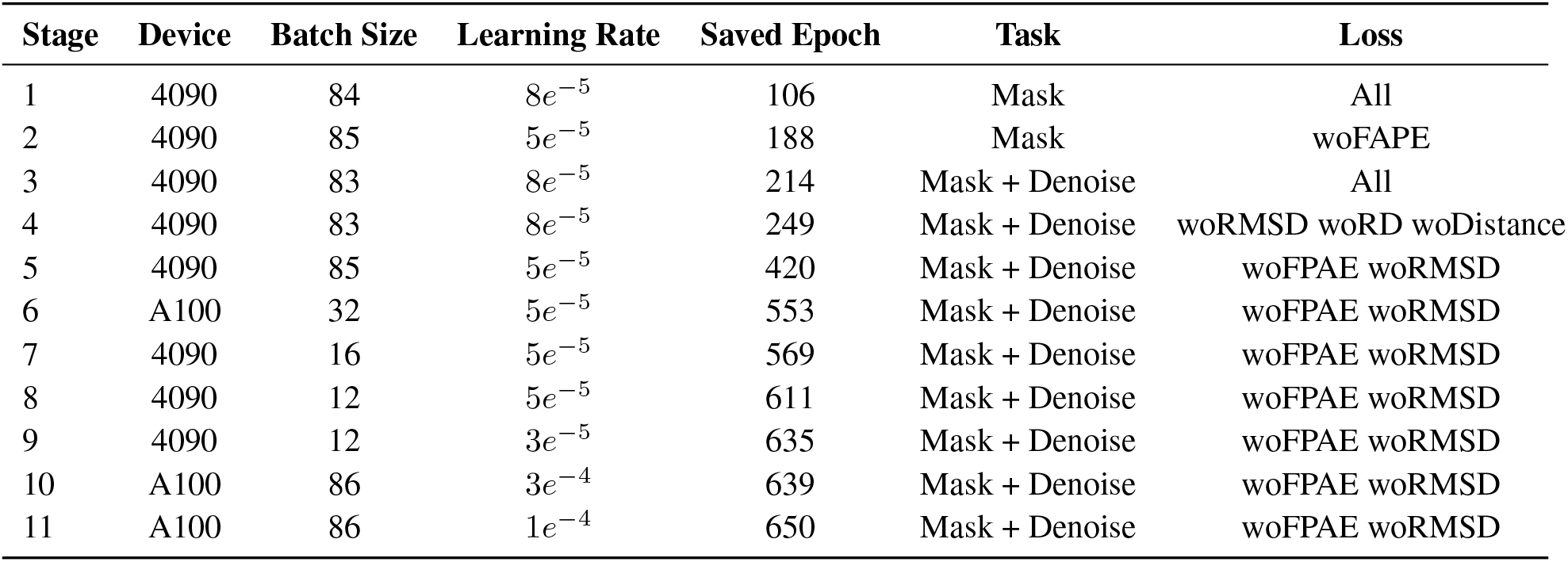
Pre-training process.

| Stage | Device | Batch Size | Learning Rate | Saved Epoch | Task | Loss |
| --- | --- | --- | --- | --- | --- | --- |
| 1 | 4090 | 84 | $8e^{-5}$ | 106 | Mask | All |
| 2 | 4090 | 85 | $5e^{-5}$ | 188 | Mask | woFAPE |
| 3 | 4090 | 83 | $8e^{-5}$ | 214 | Mask + Denoise | All |
| 4 | 4090 | 83 | $8e^{-5}$ | 249 | Mask + Denoise | woRMSD woRD woDistance |
| 5 | 4090 | 85 | $5e^{-5}$ | 420 | Mask + Denoise | woFPAE woRMSD |
| 6 | A100 | 32 | $5e^{-5}$ | 553 | Mask + Denoise | woFPAE woRMSD |
| 7 | 4090 | 16 | $5e^{-5}$ | 569 | Mask + Denoise | woFPAE woRMSD |
| 8 | 4090 | 12 | $5e^{-5}$ | 611 | Mask + Denoise | woFPAE woRMSD |
| 9 | 4090 | 12 | $3e^{-5}$ | 635 | Mask + Denoise | woFPAE woRMSD |
| 10 | A100 | 86 | $3e^{-4}$ | 639 | Mask + Denoise | woFPAE woRMSD |
| 11 | A100 | 86 | $1e^{-4}$ | 650 | Mask + Denoise | woFPAE woRMSD |

#### 2.3 Pre-training results

**Supplementary Figure 3.**
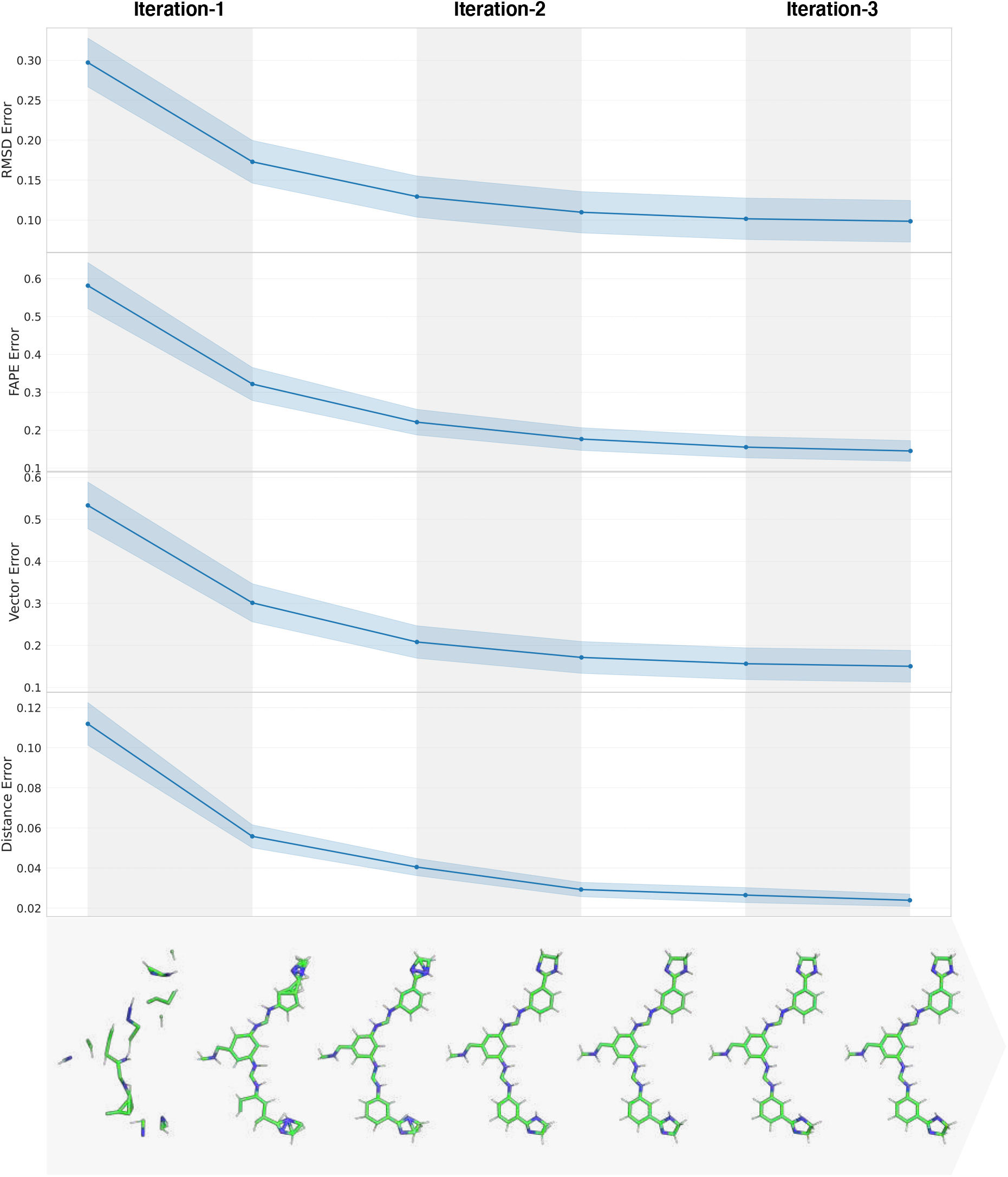
Effective structural reconstruction by the folding optimizer. During the pre-training of qcGEM, the folding optimizer learns to optimize the denoised molecular structure, through the comprehensive use of multiple geometry loss terms.

**Supplementary Figure 4.**
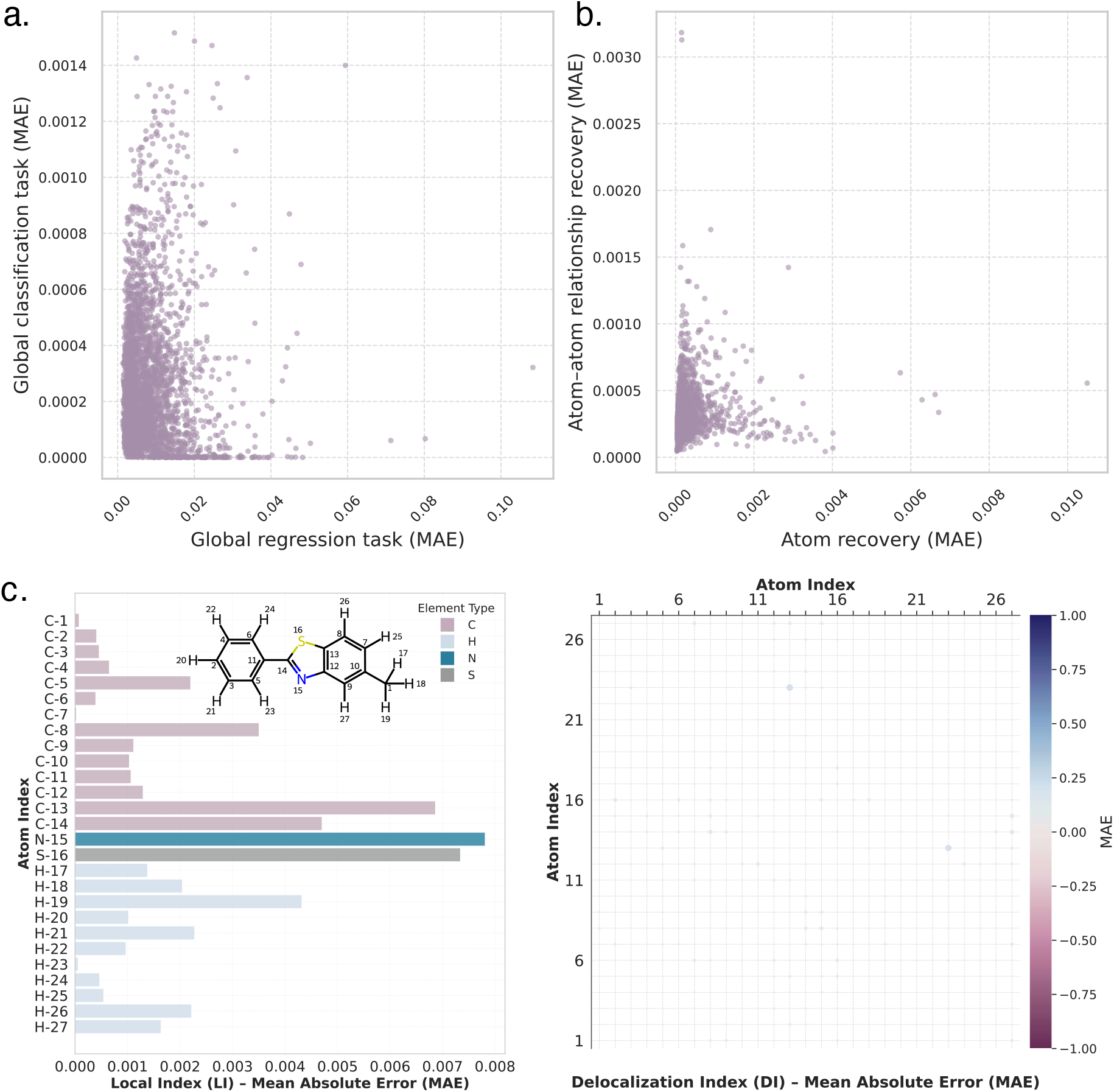
Recovery of quantum descriptors from the learned representation. **a)** Errors in the reconstruction of global properties. Each point refers to a compound, with the horizontal and vertical axes representing regression and classification tasks, respectively. **b)** Errors in the reconstruction of detailed quantum chemical information. Here, the horizontal and vertical axes represent node-level (*i.e*. atom) and edge-level (*i.e*. atom-atom relationship) quantum descriptors, respectively. **c)** Reconstruction of local features for an exemplar molecule. Errors of the localization index (LI, a node-level quantum feature) are shown in the left panel as a bar plot, whereas those of the delocalization index (DI, an edge-level quantum feature) are shown in the right panel as a heat map. The mean absolute error (MAE) is uniformly taken as the evaluation metric in this figure.

**Supplementary Figure 5.**
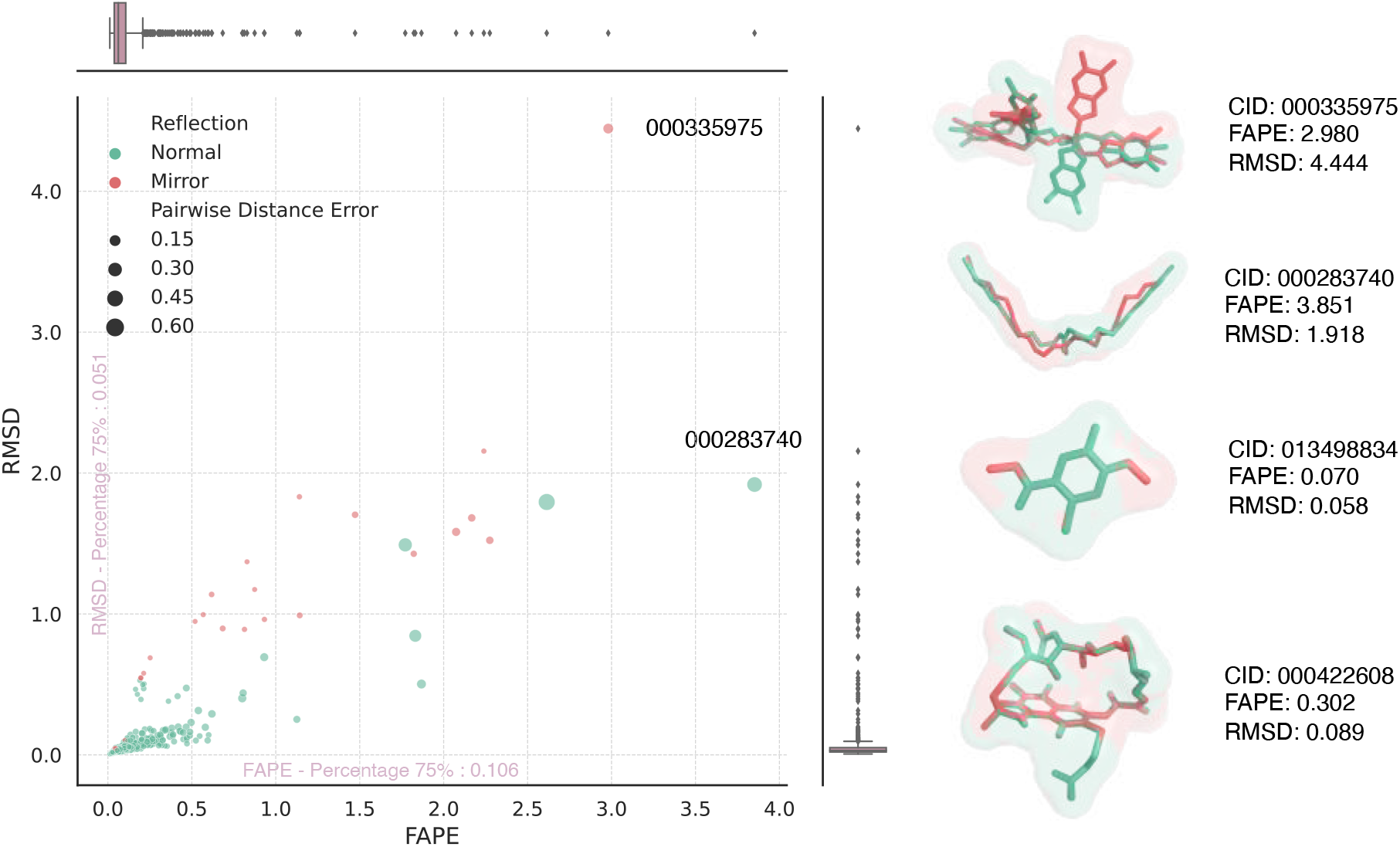
Reconstruction of molecular geometry from the learned representation. The scatter plot in the left panel presents the FAPE, RMSD, and PD (*i.e*. pairwise distance error) loss terms for molecules generated from the structure decoder, where the horizontal and vertical axes stand for FAPE and RMSD values, respectively, and each dot refers to one generated molecule, with dot size proportional to PD value. The PD term alone could not provide effective constraining, since molecules with large structure deviations (in RMSD and FAPE) frequently have small errors. Moreover, RMSD is ineffective in identifying mirror structures (*i.e*. stereoisomers). Figures in the right panel showcase exemplar generated molecules, with FAPE and RMSD values indicated aside. Clearly, the joint use of FAPE and RMSD ensures proper evaluation of structure generation. In general, the structure decoder of qcGEM allows successful generation of molecular structures for most tested molecules, as evidenced by the low FAPE and RMSD values.

### 3 Supplementary Results for Evaluation

#### 3.1 Evaluated methods

**Supplementary Table 4.** Representation methods tested in this study.

| Method | Type | Data demand | Dimension |
| --- | --- | --- | --- |
| qcGEM | Neural Embedding | 300 thousand | 128/128/128 |
| ECFP<sup>1</sup> | Molecular Fingerprint | / | 384 |
| FCFP<sup>1</sup> | Molecular Fingerprint | / | 384 |
| AvalonFP<sup>2</sup> | Molecular Fingerprint | / | 384 |
| TTGEN<sup>3</sup> | Molecular Fingerprint | / | 384 |
| APGEN<sup>4</sup> | Molecular Fingerprint | / | 384 |
| RDKFP<sup>5</sup> | Molecular Fingerprint | / | 384 |
| MACCS<sup>6</sup> | Molecular Fingerprint | / | 384 |
| LayeredFP<sup>5</sup> | Molecular Fingerprint | / | 384 |
| PatternFP<sup>5</sup> | Molecular Fingerprint | / | 384 |
| Molformer<sup>7</sup> | Neural Embedding | 11.1 billion | 768 |
| GROVER-Base<sup>8</sup> | Neural Embedding | 11 million | 3,400 |
| GROVER-Large<sup>8</sup> | Neural Embedding | 11 million | 5,000 |
| MoleBERT<sup>9</sup> | Neural Embedding | 2 million | 300 |
| UniMol-G<sup>10</sup> | Neural Embedding | 12 million + 19 million | 512 |
| UniMol-GA<sup>10</sup> | Neural Embedding | 12 million + 19 million | 1,024 |
| GCC<sup>11</sup> | Neural Embedding | 100 thousand | 128 |
| GPT-GNN<sup>12</sup> | Neural Embedding | 100 thousand | 128 |
| GraphCL<sup>13</sup> | Neural Embedding | 100 thousand | 128 |
| InfoGraph<sup>14</sup> | Neural Embedding | 100 thousand | 128 |
| JOAO<sup>15</sup> | Neural Embedding | 100 thousand | 128 |
| JOAOv2<sup>16</sup> | Neural Embedding | 100 thousand | 128 |
| PosPred<sup>17</sup> | Neural Embedding | 100 thousand | 128 |
| 3D-EMGP<sup>17</sup> | Neural Embedding | 100 thousand | 128 |
| 3D-InfoMax<sup>18</sup> | Neural Embedding | 100 thousand | 128 |
| AttrMask<sup>19</sup> | Neural Embedding | 100 thousand | 128 |
| EdgePred<sup>20</sup> | Neural Embedding | 100 thousand | 128 |
| GraphMVP<sup>21</sup> | Neural Embedding | 250 thousand | 300 |
| MolCLR-GCN<sup>22</sup> | Neural Embedding | 10 million | 512 |
| MolCLR-GIN<sup>22</sup> | Neural Embedding | 10 million | 512 |

Basic characteristics of different representation methods are shown in the table. Fingerprint-based representations are generated from pre-defined rules, whereas other approaches are learned from data.

#### 3.2 Evaluation datasets

**Supplementary Table 5.**
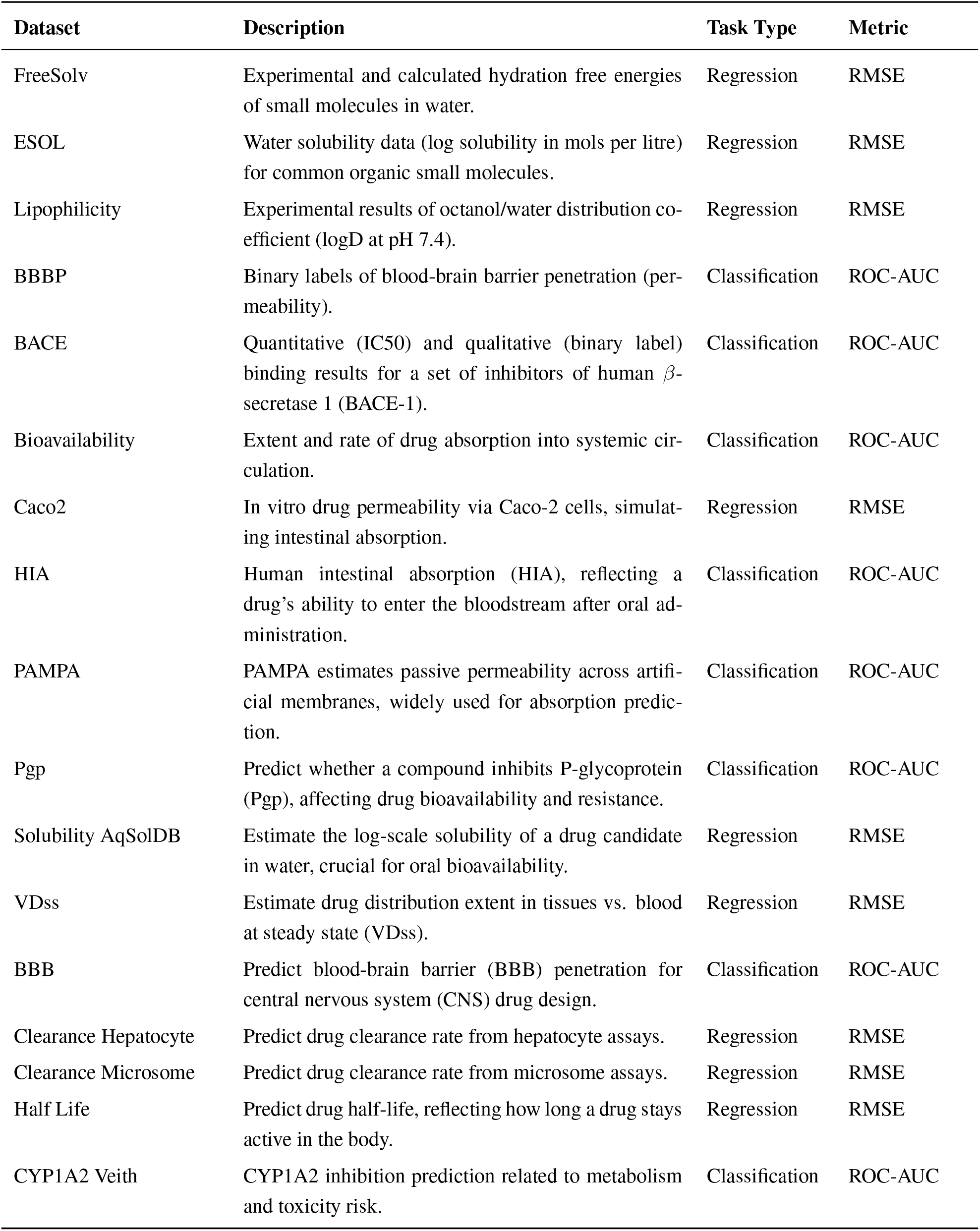
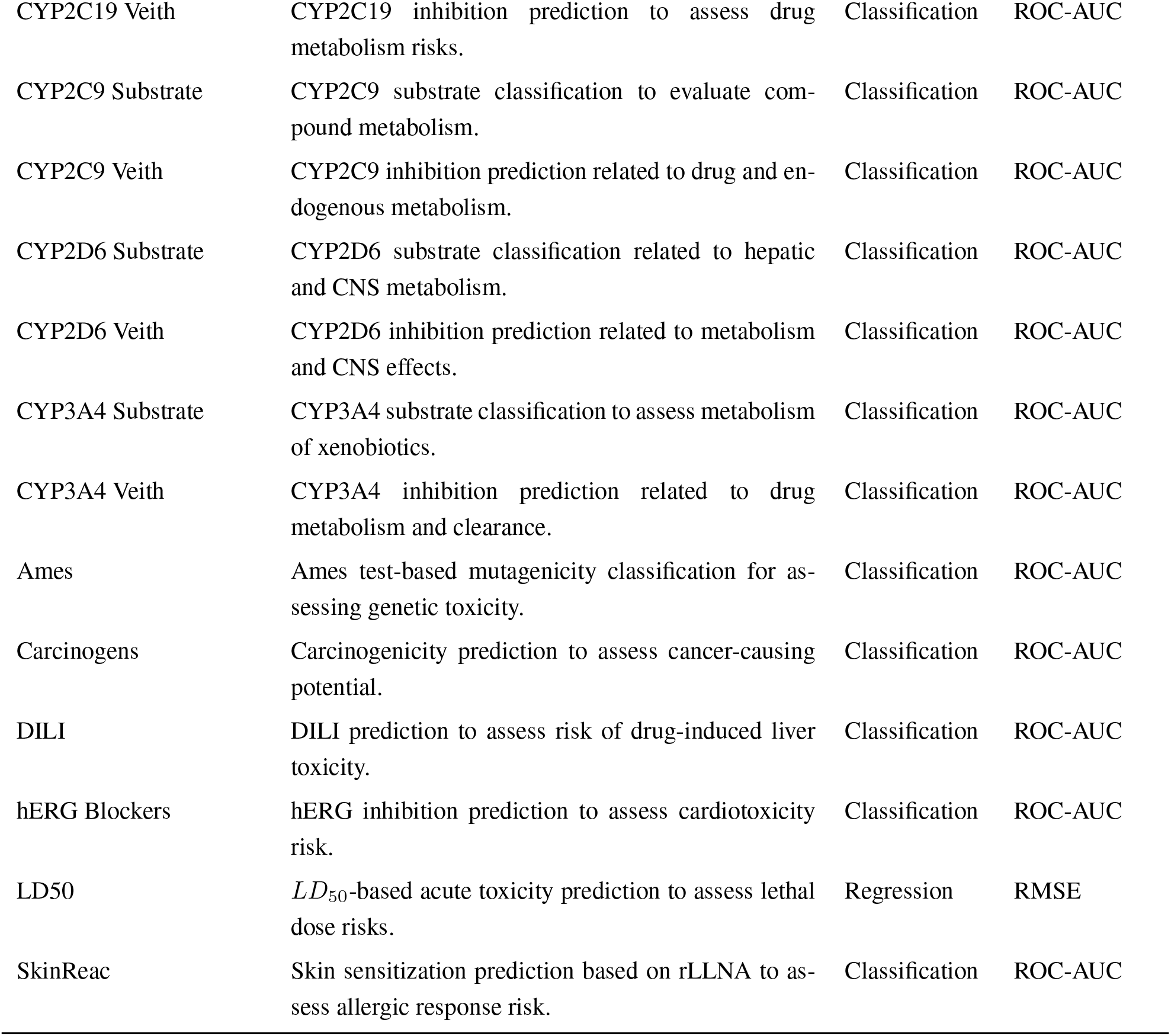
ADMET datasets.

| Dataset | Description | Task Type | Metric |
| --- | --- | --- | --- |
| FreeSolv | Experimental and calculated hydration free energies of small molecules in water. | Regression | RMSE |
| ESOL | Water solubility data (log solubility in mols per litre) for common organic small molecules. | Regression | RMSE |
| Lipophilicity | Experimental results of octanol/water distribution coefficient (logD at pH 7.4). | Regression | RMSE |
| BBBP | Binary labels of blood-brain barrier penetration (permeability). | Classification | ROC-AUC |
| BACE | Quantitative (IC50) and qualitative (binary label) binding results for a set of inhibitors of human β-secretase 1 (BACE-1). | Classification | ROC-AUC |
| Bioavailability | Extent and rate of drug absorption into systemic circulation. | Classification | ROC-AUC |
| Caco2 | In vitro drug permeability via Caco-2 cells, simulating intestinal absorption. | Regression | RMSE |
| HIA | Human intestinal absorption (HIA), reflecting a drug's ability to enter the bloodstream after oral administration. | Classification | ROC-AUC |
| PAMPA | PAMPA estimates passive permeability across artificial membranes, widely used for absorption prediction. | Classification | ROC-AUC |
| Pgp | Predict whether a compound inhibits P-glycoprotein (Pgp), affecting drug bioavailability and resistance. | Classification | ROC-AUC |
| Solubility AqSolDB | Estimate the log-scale solubility of a drug candidate in water, crucial for oral bioavailability. | Regression | RMSE |
| VDss | Estimate drug distribution extent in tissues vs. blood at steady state (VDss). | Regression | RMSE |
| BBB | Predict blood-brain barrier (BBB) penetration for central nervous system (CNS) drug design. | Classification | ROC-AUC |
| Clearance Hepatocyte | Predict drug clearance rate from hepatocyte assays. | Regression | RMSE |
| Clearance Microsome | Predict drug clearance rate from microsome assays. | Regression | RMSE |
| Half Life | Predict drug half-life, reflecting how long a drug stays active in the body. | Regression | RMSE |
| CYP1A2 Veith | CYP1A2 inhibition prediction related to metabolism and toxicity risk. | Classification | ROC-AUC |
| CYP2C19 Veith | CYP2C19 inhibition prediction to assess drug metabolism risks. | Classification | ROC-AUC |
| CYP2C9 Substrate | CYP2C9 substrate classification to evaluate compound metabolism. | Classification | ROC-AUC |
| CYP2C9 Veith | CYP2C9 inhibition prediction related to drug and endogenous metabolism. | Classification | ROC-AUC |
| CYP2D6 Substrate | CYP2D6 substrate classification related to hepatic and CNS metabolism. | Classification | ROC-AUC |
| CYP2D6 Veith | CYP2D6 inhibition prediction related to metabolism and CNS effects. | Classification | ROC-AUC |
| CYP3A4 Substrate | CYP3A4 substrate classification to assess metabolism of xenobiotics. | Classification | ROC-AUC |
| CYP3A4 Veith | CYP3A4 inhibition prediction related to drug metabolism and clearance. | Classification | ROC-AUC |
| Ames | Ames test-based mutagenicity classification for assessing genetic toxicity. | Classification | ROC-AUC |
| Carcinogens | Carcinogenicity prediction to assess cancer-causing potential. | Classification | ROC-AUC |
| DILI | DILI prediction to assess risk of drug-induced liver toxicity. | Classification | ROC-AUC |
| hERG Blockers | hERG inhibition prediction to assess cardiotoxicity risk. | Classification | ROC-AUC |
| LD50 | $LD_{50}$-based acute toxicity prediction to assess lethal dose risks. | Regression | RMSE |
| SkinReac | Skin sensitization prediction based on rLLNA to assess allergic response risk. | Classification | ROC-AUC |

**Supplementary Table 6.**
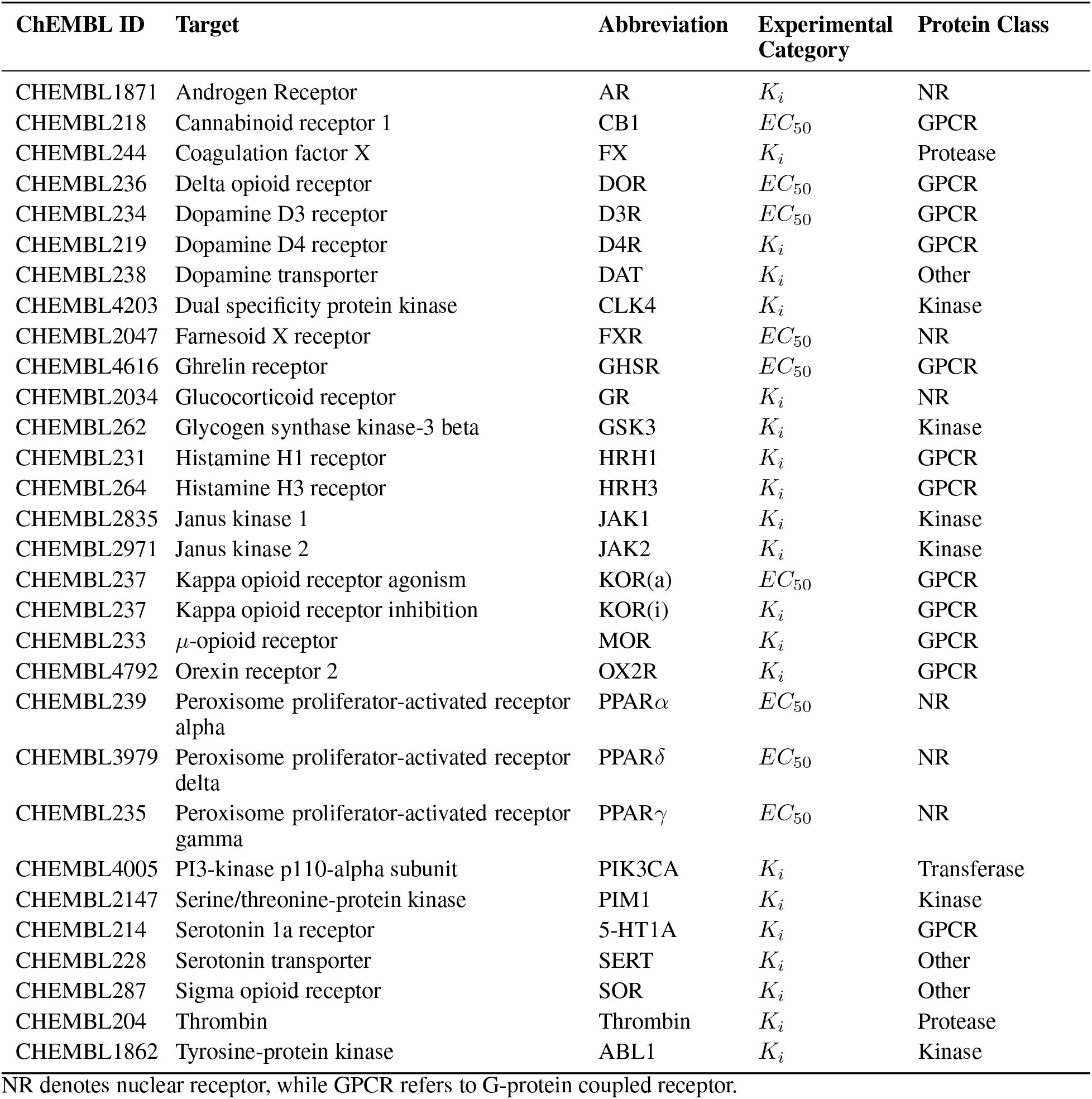
MoleculeACE datasets.

**Supplementary Table 7.** Protein-ligand interaction datasets.

| **Dataset** | **Description** | **Task Type** | **Metric** |
| --- | --- | --- | --- |
| Human | Derived from the DrugBank database, containing experimentally validated compound–protein interactions involving human proteins. | Classification | ROC-AUC |
| C.elegans | Contain interactions between compounds and *C. elegans* proteins, sourced from curated public databases. | Classification | ROC-AUC |
| KIBA | A model-based integration of bioactivities into a unified drug-target interaction matrix. | Regression | RMSE |
| DAVIS | The interaction of 72 kinase inhibitors with 442 kinases covering >80% of the human catalytic protein kinome. | Regression | RMSE |
| BindingDB | BindingDB is a public database of experimentally measured binding affinities between drug-target proteins and small molecules. | Classification | ROC-AUC |

**Supplementary Table 8.** Opioid-type datasets.

| **Dataset** | **Protein Name** | **ChEMBL ID** | **Task Type** | **Metric** |
| --- | --- | --- | --- | --- |
| CYP2D6 | Cytochrome P450 2D6 | ChEMBL 289 | Regression | RMSE |
| CYP3A4 | Cytochrome P450 3A4 | ChEMBL 340 | Regression | RMSE |
| DOR | δ-opioid receptor | ChEMBL 236 | Regression | RMSE |
| KOR | κ-opioid receptor | ChEMBL 237 | Regression | RMSE |
| MDR1 | Multidrug resistance protein 1 (P-glycoprotein 1) | ChEMBL 4302 | Regression | RMSE |
| MOR | μ-opioid receptor | ChEMBL 340 | Regression | RMSE |

#### 3.3 Evaluation results

##### 3.3.1 Molecular property prediction

**Supplementary Figure 6.**
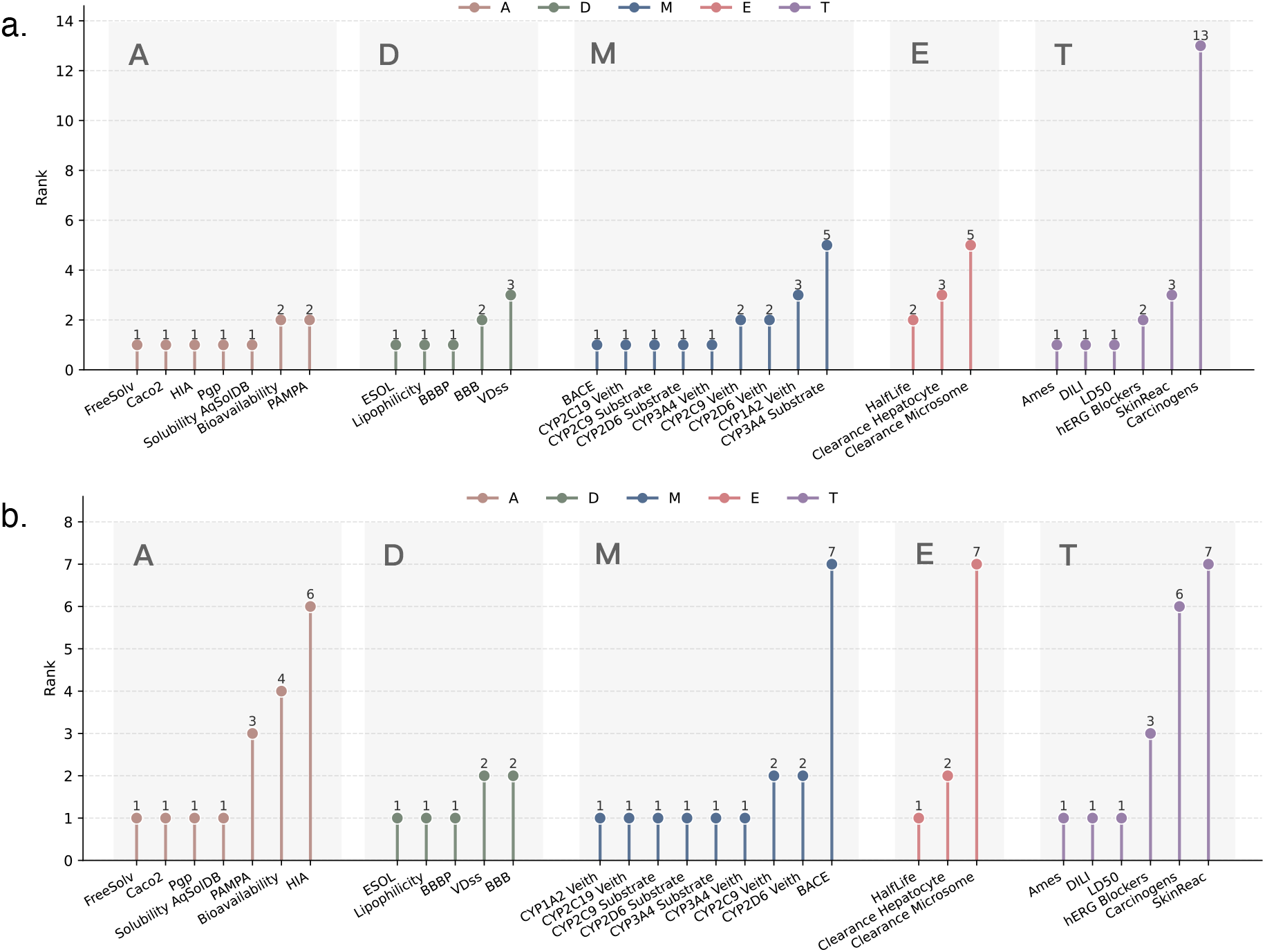
Detailed ranking information of qcGEM in individual ADMET tasks. **a)** In the scaffold split setting, qcGEM rankings are shown from left to right for absorption (A), distribution (D), metabolism (M), excretion (E), and toxicity (T). **b)** Results are shown for the random split setting following the same convention.

**Supplementary Figure 7.**
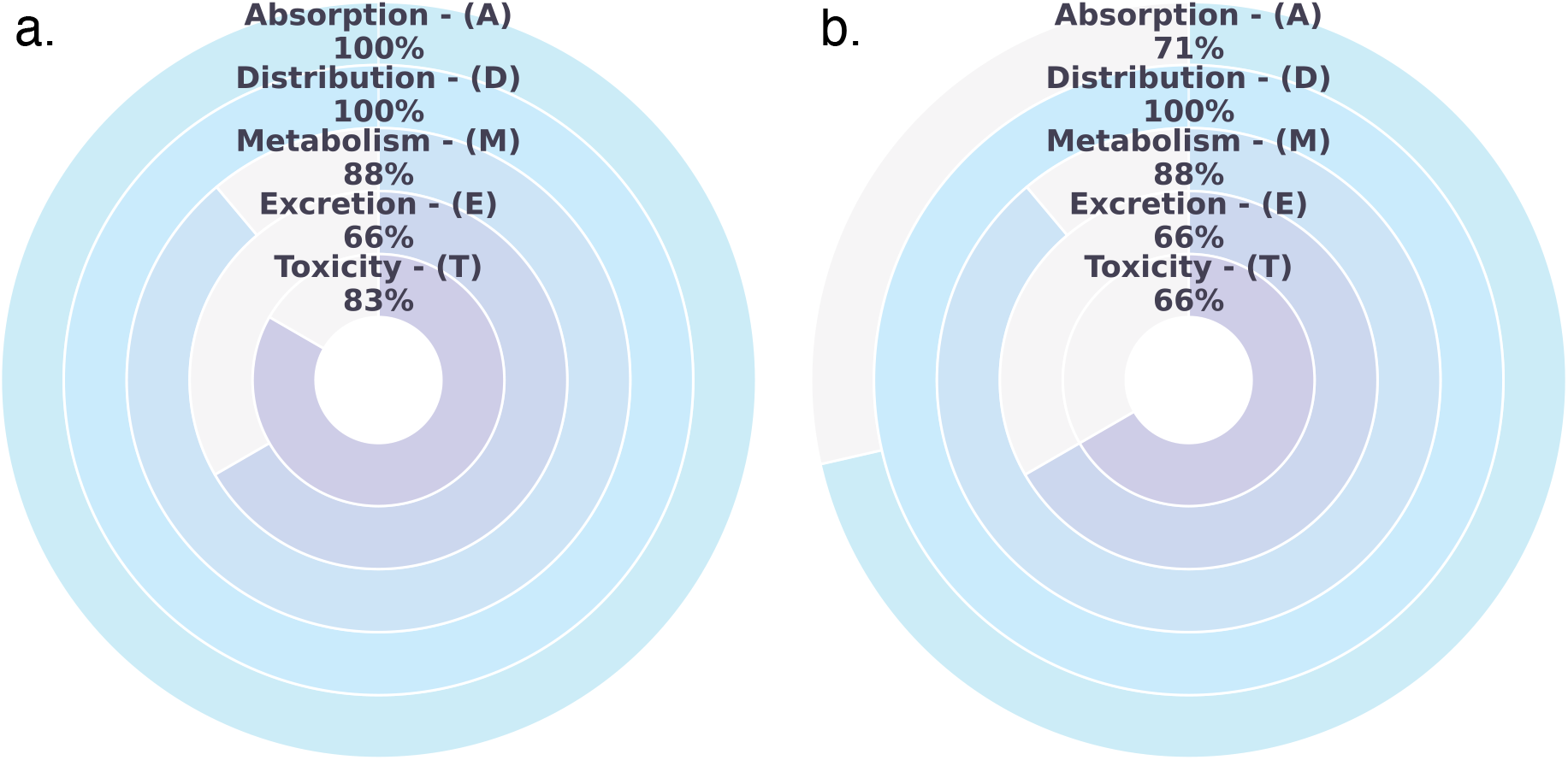
Proportion of tasks in which qcGEM achieves the top 3 ranking across the 5 ADMET task categories. **a)** Performance of qcGEM across ADMET task categories for all molecules in the scaffold split setting. Here, ADMET tasks are classified into absorption (A), distribution (D), metabolism (M), excretion (E), and toxicity (T). **b)** Performance of qcGEM across ADMET task categories for all molecules in the random split setting.

**Supplementary Figure 8.**
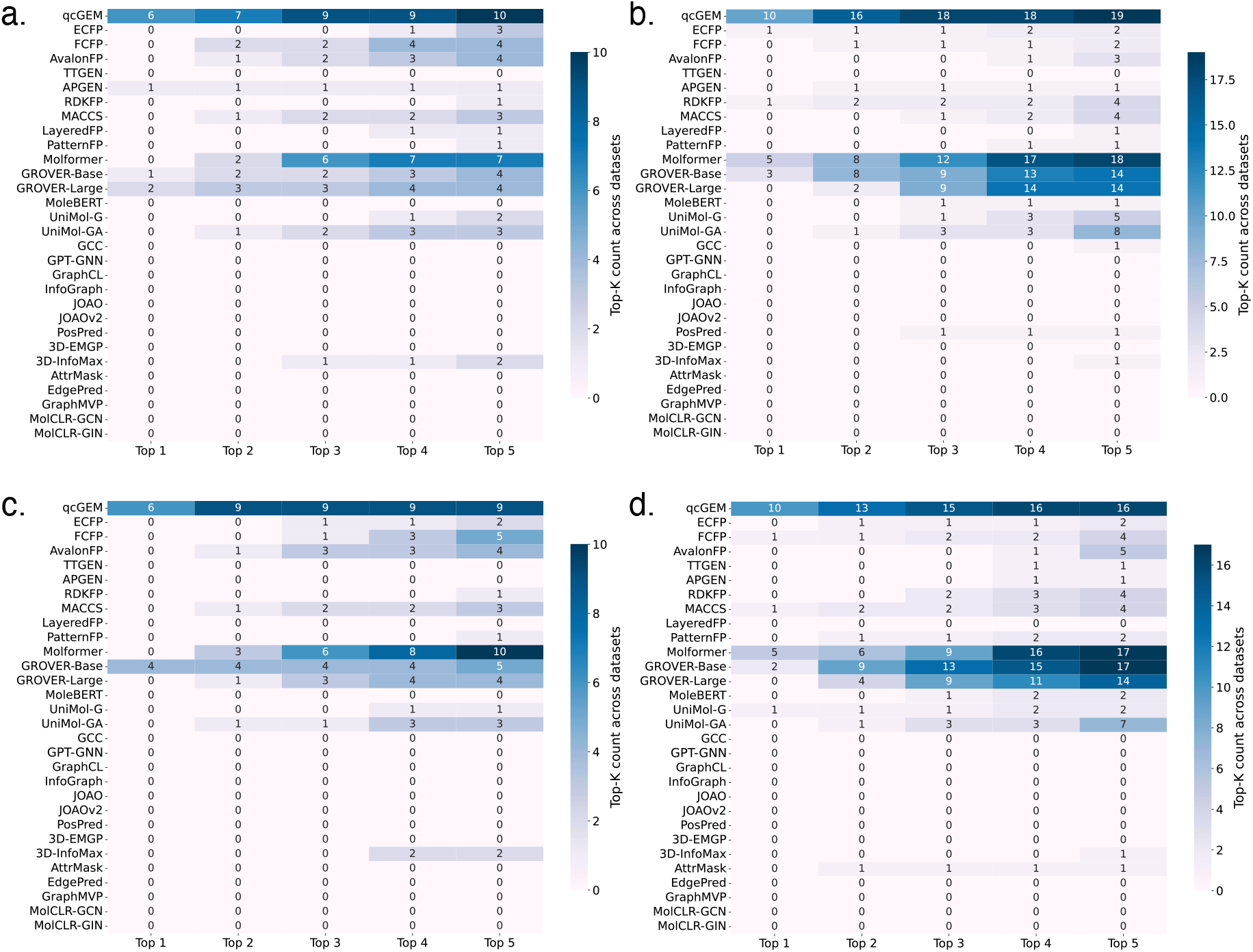
Statistics of the top 1-5 rankings for all methods in ADMET Tasks. **a)** Regression tasks in the scaffold split setting. **b)** Classification tasks in the scaffold split setting. **c)** Regression tasks in the random split setting. **d)** Classification tasks in the random split setting.

**Supplementary Table 9.** Molecular property regression task (scaffold split) – Part I.

| Method | FreeSolv | ESOL | Lipophilicity | Caco2 | Solubility<br>AqSolDB |
| --- | --- | --- | --- | --- | --- |
| qcGEM | 0.815 ± 0.232 | 0.564 ± 0.0468 | 0.637 ± 0.037 | 0.388 ± 0.0387 | 0.831 ± 0.0447 |
| ECFP | 2.67 ± 0.583 | 1.35 ± 0.129 | 0.89 ± 0.0294 | 0.594 ± 0.0625 | 1.29 ± 0.0278 |
| FCFP | 1.93 ± 0.45 | 1.37 ± 0.0973 | 0.873 ± 0.0199 | 0.488 ± 0.019 | 1.25 ± 0.0335 |
| AvalonFP | 1.53 ± 0.176 | 1.04 ± 0.068 | 0.898 ± 0.0544 | 0.557 ± 0.03 | 1.14 ± 0.0384 |
| TTGEN | 3.15 ± 0.505 | 1.28 ± 0.1 | 0.959 ± 0.0396 | 0.583 ± 0.0331 | 1.31 ± 0.0327 |
| APGEN | 2.35 ± 0.206 | 1.08 ± 0.0819 | 0.951 ± 0.0402 | 0.619 ± 0.0868 | 1.21 ± 0.0402 |
| RDKFP | 1.92 ± 0.388 | 1.23 ± 0.0283 | 0.883 ± 0.0483 | 0.575 ± 0.1 | 1.36 ± 0.0267 |
| MACCS | 1.16 ± 0.306 | 1.07 ± 0.108 | 0.878 ± 0.0543 | 0.46 ± 0.0344 | 1.19 ± 0.0334 |
| LayeredFP | 2.02 ± 0.357 | 0.941 ± 0.115 | 1.06 ± 0.0391 | 0.641 ± 0.134 | 1.22 ± 0.0339 |
| PatternFP | 1.95 ± 0.351 | 0.851 ± 0.111 | 0.955 ± 0.0432 | 0.63 ± 0.0773 | 1.15 ± 0.0192 |
| Molformer | 1.66 ± 0.219 | 0.841 ± 0.0838 | 0.7 ± 0.0387 | 0.445 ± 0.0321 | 0.901 ± 0.0415 |
| GROVER-Base | 3.73 ± 0.483 | 2.16 ± 0.084 | 1.18 ± 0.0576 | 0.637 ± 0.0541 | 0.967 ± 0.0515 |
| GROVER-Large | 3.72 ± 0.479 | 2.16 ± 0.0826 | 1.18 ± 0.056 | 0.849 ± 0.686 | 0.893 ± 0.0393 |
| MoleBERT | 2.35 ± 0.528 | 1.13 ± 0.149 | 1.01 ± 0.0451 | 0.56 ± 0.0564 | 1.36 ± 0.0367 |
| UniMol-G | 1.65 ± 0.258 | 0.846 ± 0.0497 | 0.981 ± 0.0471 | 0.567 ± 0.0578 | 1.03 ± 0.0207 |
| UniMol-GA | 1.4 ± 0.205 | 0.749 ± 0.0547 | 0.925 ± 0.0445 | 0.615 ± 0.0707 | 0.955 ± 0.0248 |
| GCC | 2.1 ± 0.388 | 1.06 ± 0.117 | 1.05 ± 0.0437 | 0.593 ± 0.0456 | 1.24 ± 0.0217 |
| GPT-GNN | 2.18 ± 0.256 | 0.977 ± 0.112 | 1.01 ± 0.0549 | 0.809 ± 0.0928 | 1.22 ± 0.0325 |
| GraphCL | 2.66 ± 0.804 | 5 ± 0 | 2.06 ± 1.75 | 5 ± 0 | 5 ± 0 |
| InfoGraph | 3.08 ± 0.509 | 1.36 ± 0.0979 | 1.13 ± 0.0515 | 0.683 ± 0.0668 | 1.47 ± 0.035 |
| JOAO | 2.47 ± 0.361 | 1.08 ± 0.103 | 1.09 ± 0.0465 | 0.85 ± 0.102 | 1.34 ± 0.0613 |
| JOAOv2 | 2.14 ± 0.359 | 0.995 ± 0.112 | 1.05 ± 0.0556 | 0.67 ± 0.0739 | 1.22 ± 0.0608 |
| PosPred | 2.53 ± 0.431 | 1.12 ± 0.178 | 1.1 ± 0.0485 | 2.67 ± 2.23 | 1.41 ± 0.117 |
| 3D-EMGP | 2.86 ± 1.14 | 2.79 ± 0.854 | 2.38 ± 2.28 | 5 ± 0 | 5 ± 0 |
| 3D-InfoMax | 2.15 ± 0.337 | 0.945 ± 0.0322 | 0.867 ± 0.0422 | 0.512 ± 0.0496 | 1.13 ± 0.0523 |
| AttrMask | 1.94 ± 0.435 | 0.98 ± 0.104 | 1.03 ± 0.0457 | 5 ± 0 | 5 ± 0 |
| EdgePred | 3.14 ± 0.493 | 5 ± 0 | 1.21 ± 0.0821 | 5 ± 0 | 1.81 ± 0.0704 |
| GraphMVP | 3.71 ± 0.472 | 2.15 ± 0.0833 | 1.18 ± 0.0592 | 0.76 ± 0.099 | 2.25 ± 0.068 |
| MolCLR-GCN | 3.74 ± 0.527 | 2.15 ± 0.082 | 1.18 ± 0.0578 | 0.761 ± 0.102 | 2.23 ± 0.065 |
| MolCLR-GIN | 3.84 ± 0.481 | 2.14 ± 0.086 | 1.18 ± 0.0582 | 0.776 ± 0.0924 | 2.23 ± 0.0672 |

Prediction is evaluated by RMSE. For each method, the mean and standard deviation are reported for the evaluation metrics (Values greater than 5 were capped at 5 for readability purposes) .

**Supplementary Table 10.** Molecular property regression task (scaffold split) – Part II.

| Method | VDss | Clearance<br>Hepatocyte | Clearance<br>Microsome | Half Life | LD50 |
| --- | --- | --- | --- | --- | --- |
| qcGEM | 0.0566 ± 0.0946 | 0.271 ± 0.0267 | 0.242 ± 0.0227 | 0.0238 ± 0.0184 | 0.603 ± 0.0211 |
| ECFP | 0.0566 ± 0.0946 | 0.281 ± 0.0387 | 0.249 ± 0.0237 | 0.0239 ± 0.018 | 0.712 ± 0.029 |
| FCFP | 0.0559 ± 0.0929 | 0.293 ± 0.036 | 0.233 ± 0.0199 | 0.024 ± 0.0178 | 0.718 ± 0.0254 |
| AvalonFP | 0.0567 ± 0.0952 | 0.302 ± 0.0354 | 0.259 ± 0.0253 | 0.0239 ± 0.018 | 0.632 ± 0.0195 |
| TTGEN | 0.059 ± 0.0977 | 0.307 ± 0.0465 | 0.25 ± 0.0228 | 0.0242 ± 0.0183 | 0.762 ± 0.0198 |
| APGEN | 0.0576 ± 0.0954 | 0.304 ± 0.0208 | 0.258 ± 0.0195 | 0.0237 ± 0.0184 | 0.696 ± 0.0272 |
| RDKFP | 0.058 ± 0.097 | 0.299 ± 0.0231 | 0.249 ± 0.0147 | 0.0241 ± 0.0182 | 0.669 ± 0.0239 |
| MACCS | 0.0576 ± 0.096 | 0.295 ± 0.0327 | 0.246 ± 0.0181 | 0.0242 ± 0.0181 | 0.699 ± 0.0304 |
| LayeredFP | 0.0588 ± 0.0975 | 0.292 ± 0.0387 | 0.259 ± 0.0217 | 0.0239 ± 0.0181 | 0.682 ± 0.026 |
| PatternFP | 0.0588 ± 0.0976 | 0.298 ± 0.0183 | 0.254 ± 0.019 | 0.0239 ± 0.0186 | 0.707 ± 0.0245 |
| Molformer | 0.057 ± 0.0962 | 0.277 ± 0.0227 | 0.236 ± 0.0253 | 0.024 ± 0.0179 | 0.638 ± 0.0141 |
| GROVER-Base | 0.0558 ± 0.0936 | 0.269 ± 0.0306 | 0.241 ± 0.0243 | 0.0242 ± 0.0179 | 0.722 ± 0.0511 |
| GROVER-Large | 0.0572 ± 0.0966 | 0.269 ± 0.0287 | 0.233 ± 0.0245 | 0.0251 ± 0.0175 | 0.65 ± 0.0456 |
| MoleBERT | 0.0594 ± 0.098 | 0.307 ± 0.0287 | 0.272 ± 0.027 | 0.0245 ± 0.0182 | 0.763 ± 0.0327 |
| UniMol-G | 0.0585 ± 0.0974 | 0.298 ± 0.0408 | 0.254 ± 0.0269 | 0.0245 ± 0.0186 | 0.739 ± 0.0264 |
| UniMol-GA | 0.0581 ± 0.0968 | 0.294 ± 0.0365 | 0.25 ± 0.0249 | 0.0242 ± 0.0188 | 0.711 ± 0.0331 |
| GCC | 0.0592 ± 0.0973 | 0.312 ± 0.0375 | 0.271 ± 0.032 | 0.0248 ± 0.0189 | 0.778 ± 0.0286 |
| GPT-GNN | 0.0623 ± 0.0976 | 0.309 ± 0.0345 | 0.261 ± 0.0265 | 0.0255 ± 0.0187 | 0.737 ± 0.0381 |
| GraphCL | 5 ± 0 | 5 ± 0 | 5 ± 0 | 5 ± 0 | 5 ± 0 |
| InfoGraph | 0.0595 ± 0.098 | 0.319 ± 0.0281 | 0.261 ± 0.0305 | 0.0244 ± 0.0184 | 0.909 ± 0.0251 |
| JOAO | 0.0664 ± 0.102 | 0.311 ± 0.0306 | 0.266 ± 0.0304 | 0.111 ± 0.14 | 0.83 ± 0.0243 |
| JOAOv2 | 0.0609 ± 0.0979 | 0.311 ± 0.0328 | 0.267 ± 0.0285 | 0.0245 ± 0.0189 | 0.793 ± 0.0244 |
| PosPred | 0.41 ± 0.595 | 0.313 ± 0.0322 | 0.274 ± 0.0317 | 0.275 ± 0.384 | 0.868 ± 0.0509 |
| 3D-EMGP | 5 ± 0 | 5 ± 0 | 5 ± 0 | 5 ± 0 | 5 ± 0 |
| 3D-InfoMax | 0.0586 ± 0.0977 | 0.307 ± 0.0194 | 0.253 ± 0.0342 | 0.0242 ± 0.0183 | 0.753 ± 0.0241 |
| AttrMask | 0.0593 ± 0.0976 | 0.317 ± 0.0453 | 0.267 ± 0.0256 | 5 ± 0 | 0.809 ± 0.0403 |
| EdgePred | 5 ± 0 | 5 ± 0 | 5 ± 0 | 5 ± 0 | 0.929 ± 0.0326 |
| GraphMVP | 0.0596 ± 0.098 | 0.32 ± 0.0397 | 0.273 ± 0.0273 | 0.0246 ± 0.0186 | 0.952 ± 0.0293 |
| MolCLR-GCN | 0.0597 ± 0.0982 | 0.318 ± 0.0374 | 0.273 ± 0.0273 | 0.0246 ± 0.0186 | 0.955 ± 0.0298 |
| MolCLR-GIN | 0.0597 ± 0.0982 | 0.32 ± 0.0426 | 0.273 ± 0.0272 | 0.0246 ± 0.0184 | 0.953 ± 0.0297 |

Prediction is evaluated by RMSE. For each method, the mean and standard deviation are reported for the evaluation metrics (Values greater than 5 were capped at 5 for readability purposes) .

**Supplementary Table 11.**
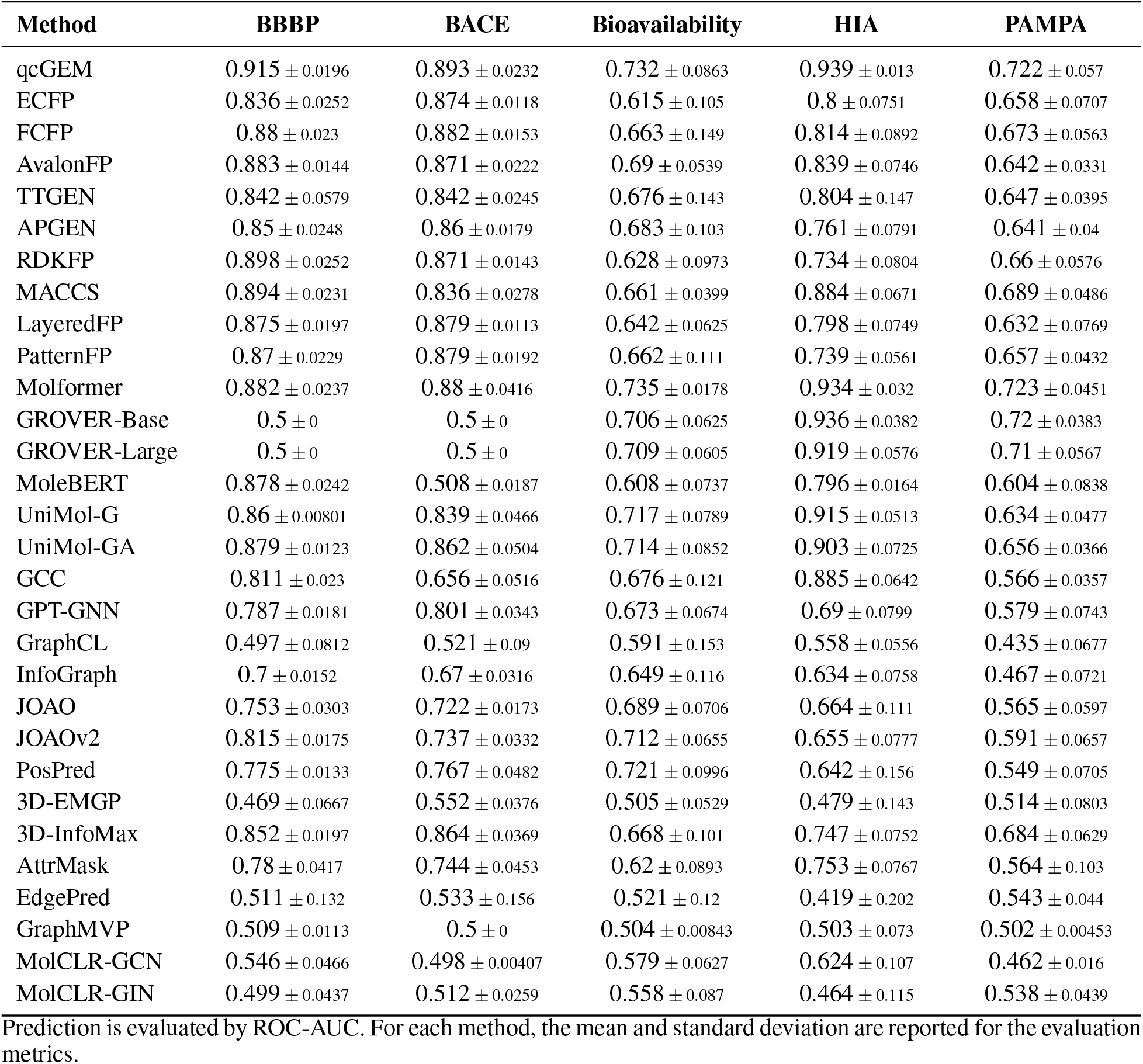
Molecular property classification task (scaffold split) – Part I.

**Supplementary Table 12.** Molecular property classification task (scaffold split) – Part II.

| Method | Pgp | BBB | CYP1A2 Veith | CYP2C19 Veith | CYP2C9 Substrate |
| --- | --- | --- | --- | --- | --- |
| qcGEM | 0.941 ± 0.0175 | 0.889 ± 0.0242 | 0.925 ± 0.00671 | 0.896 ± 0.00546 | 0.722 ± 0.0694 |
| ECFP | 0.909 ± 0.0314 | 0.851 ± 0.0461 | 0.851 ± 0.00833 | 0.662 ± 0.148 | 0.617 ± 0.0888 |
| FCFP | 0.906 ± 0.0359 | 0.866 ± 0.0155 | 0.871 ± 0.00589 | 0.673 ± 0.158 | 0.596 ± 0.116 |
| AvalonFP | 0.911 ± 0.0244 | 0.878 ± 0.0243 | 0.88 ± 0.00349 | 0.842 ± 0.00493 | 0.581 ± 0.128 |
| TTGEN | 0.91 ± 0.0325 | 0.84 ± 0.0315 | 0.86 ± 0.00474 | 0.649 ± 0.136 | 0.63 ± 0.0429 |
| APGEN | 0.908 ± 0.0227 | 0.858 ± 0.03 | 0.885 ± 0.0036 | 0.828 ± 0.0151 | 0.636 ± 0.0436 |
| RDKFP | 0.905 ± 0.0295 | 0.876 ± 0.0353 | 0.893 ± 0.00342 | 0.84 ± 0.00839 | 0.655 ± 0.0508 |
| MACCS | 0.89 ± 0.0144 | 0.873 ± 0.0385 | 0.872 ± 0.0118 | 0.834 ± 0.00926 | 0.695 ± 0.0388 |
| LayeredFP | 0.906 ± 0.0101 | 0.856 ± 0.037 | 0.86 ± 0.006 | 0.791 ± 0.00695 | 0.605 ± 0.0671 |
| PatternFP | 0.914 ± 0.0073 | 0.858 ± 0.0323 | 0.887 ± 0.00744 | 0.817 ± 0.00568 | 0.688 ± 0.054 |
| Molformer | 0.924 ± 0.0124 | 0.9 ± 0.0172 | 0.918 ± 0.00606 | 0.891 ± 0.00673 | 0.712 ± 0.0464 |
| GROVER-Base | 0.93 ± 0.0164 | 0.881 ± 0.0121 | 0.926 ± 0.00314 | 0.893 ± 0.00544 | 0.706 ± 0.0903 |
| GROVER-Large | 0.928 ± 0.021 | 0.888 ± 0.0242 | 0.925 ± 0.00384 | 0.891 ± 0.00381 | 0.707 ± 0.0857 |
| MoleBERT | 0.564 ± 0.0828 | 0.835 ± 0.023 | 0.79 ± 0.162 | 0.528 ± 0.062 | 0.615 ± 0.0756 |
| UniMol-G | 0.89 ± 0.0239 | 0.858 ± 0.0305 | 0.888 ± 0.00849 | 0.836 ± 0.00907 | 0.589 ± 0.0255 |
| UniMol-GA | 0.906 ± 0.019 | 0.871 ± 0.0258 | 0.901 ± 0.0061 | 0.853 ± 0.0096 | 0.605 ± 0.037 |
| GCC | 0.864 ± 0.0327 | 0.783 ± 0.0527 | 0.841 ± 0.00595 | 0.791 ± 0.0144 | 0.618 ± 0.077 |
| GPT-GNN | 0.878 ± 0.0253 | 0.808 ± 0.0322 | 0.861 ± 0.00765 | 0.775 ± 0.0133 | 0.681 ± 0.0672 |
| GraphCL | 0.747 ± 0.0876 | 0.508 ± 0.0754 | 0.808 ± 0.0127 | 0.709 ± 0.0307 | 0.502 ± 0.135 |
| InfoGraph | 0.859 ± 0.0385 | 0.749 ± 0.0364 | 0.801 ± 0.00902 | 0.682 ± 0.0127 | 0.56 ± 0.0569 |
| JOAO | 0.844 ± 0.0319 | 0.725 ± 0.0518 | 0.837 ± 0.00755 | 0.73 ± 0.0123 | 0.563 ± 0.0558 |
| JOAOv2 | 0.861 ± 0.0277 | 0.808 ± 0.0431 | 0.864 ± 0.01 | 0.789 ± 0.00921 | 0.59 ± 0.0559 |
| PosPred | 0.853 ± 0.0303 | 0.704 ± 0.0719 | 0.864 ± 0.00925 | 0.758 ± 0.01 | 0.599 ± 0.101 |
| 3D-EMGP | 0.728 ± 0.0594 | 0.501 ± 0.0584 | 0.742 ± 0.0294 | 0.637 ± 0.0112 | 0.542 ± 0.0739 |
| 3D-InfoMax | 0.918 ± 0.0153 | 0.865 ± 0.03 | 0.882 ± 0.00366 | 0.832 ± 0.00662 | 0.561 ± 0.0462 |
| AttrMask | 0.881 ± 0.0439 | 0.772 ± 0.0467 | 0.829 ± 0.00983 | 0.73 ± 0.0106 | 0.592 ± 0.117 |
| EdgePred | 0.75 ± 0.127 | 0.513 ± 0.127 | 0.553 ± 0.0615 | 0.565 ± 0.0131 | 0.482 ± 0.0544 |
| GraphMVP | 0.5 ± 0 | 0.501 ± 0.0118 | 0.5 ± 0 | 0.5 ± 0 | 0.511 ± 0.0596 |
| MolCLR-GCN | 0.508 ± 0.0383 | 0.489 ± 0.0508 | 0.494 ± 0.00875 | 0.5 ± 0.000173 | 0.471 ± 0.0524 |
| MolCLR-GIN | 0.515 ± 0.0706 | 0.554 ± 0.048 | 0.496 ± 0.0118 | 0.501 ± 0.0018 | 0.485 ± 0.126 |

Prediction is evaluated by ROC-AUC. For each method, the mean and standard deviation are reported for the evaluation metrics.

**Supplementary Table 13.** Molecular property classification task (scaffold split) – Part III.

| Method | CYP2C9 Veith | CYP2D6 Substrate | CYP2D6 Veith | CYP3A4 Substrate | CYP3A4 Veith |
| --- | --- | --- | --- | --- | --- |
| qcGEM | 0.893 ± 0.00364 | 0.762 ± 0.051 | 0.872 ± 0.0214 | 0.663 ± 0.0778 | 0.888 ± 0.00845 |
| ECFP | 0.836 ± 0.00812 | 0.704 ± 0.0369 | 0.794 ± 0.0195 | 0.665 ± 0.0609 | 0.569 ± 0.155 |
| FCFP | 0.844 ± 0.0108 | 0.74 ± 0.0596 | 0.816 ± 0.0226 | 0.654 ± 0.067 | 0.635 ± 0.186 |
| AvalonFP | 0.844 ± 0.00608 | 0.713 ± 0.0417 | 0.802 ± 0.0228 | 0.625 ± 0.0574 | 0.839 ± 0.0116 |
| TTGEN | 0.835 ± 0.00613 | 0.691 ± 0.0262 | 0.779 ± 0.0227 | 0.627 ± 0.107 | 0.538 ± 0.0857 |
| APGEN | 0.828 ± 0.00677 | 0.652 ± 0.103 | 0.792 ± 0.019 | 0.668 ± 0.0646 | 0.824 ± 0.0082 |
| RDKFP | 0.854 ± 0.00682 | 0.715 ± 0.0925 | 0.805 ± 0.0357 | 0.673 ± 0.103 | 0.841 ± 0.0154 |
| MACCS | 0.831 ± 0.00587 | 0.71 ± 0.0808 | 0.806 ± 0.0292 | 0.654 ± 0.0695 | 0.828 ± 0.0137 |
| LayeredFP | 0.799 ± 0.0084 | 0.697 ± 0.0554 | 0.791 ± 0.0218 | 0.646 ± 0.0611 | 0.798 ± 0.00851 |
| PatternFP | 0.823 ± 0.00919 | 0.685 ± 0.0805 | 0.794 ± 0.0323 | 0.659 ± 0.0676 | 0.826 ± 0.0101 |
| Molformer | 0.887 ± 0.00845 | 0.76 ± 0.0439 | 0.864 ± 0.0182 | 0.652 ± 0.045 | 0.883 ± 0.0087 |
| GROVER-Base | 0.893 ± 0.00421 | 0.72 ± 0.0528 | 0.873 ± 0.0241 | 0.632 ± 0.0542 | 0.884 ± 0.00815 |
| GROVER-Large | 0.891 ± 0.00409 | 0.718 ± 0.0409 | 0.852 ± 0.054 | 0.666 ± 0.0567 | 0.881 ± 0.011 |
| MoleBERT | 0.814 ± 0.0128 | 0.712 ± 0.069 | 0.765 ± 0.0277 | 0.535 ± 0.0979 | 0.578 ± 0.127 |
| UniMol-G | 0.841 ± 0.00796 | 0.758 ± 0.0687 | 0.787 ± 0.0224 | 0.642 ± 0.0635 | 0.823 ± 0.00997 |
| UniMol-GA | 0.855 ± 0.00829 | 0.759 ± 0.0682 | 0.804 ± 0.0233 | 0.646 ± 0.068 | 0.839 ± 0.00897 |
| GCC | 0.794 ± 0.00862 | 0.748 ± 0.0509 | 0.723 ± 0.0184 | 0.611 ± 0.0595 | 0.754 ± 0.0756 |
| GPT-GNN | 0.79 ± 0.0168 | 0.641 ± 0.0658 | 0.751 ± 0.0132 | 0.612 ± 0.0703 | 0.776 ± 0.0113 |
| GraphCL | 0.722 ± 0.0136 | 0.716 ± 0.0584 | 0.616 ± 0.0241 | 0.559 ± 0.0266 | 0.732 ± 0.00989 |
| InfoGraph | 0.655 ± 0.011 | 0.694 ± 0.0448 | 0.682 ± 0.011 | 0.61 ± 0.0547 | 0.714 ± 0.0101 |
| JOAO | 0.746 ± 0.0233 | 0.671 ± 0.0675 | 0.607 ± 0.0184 | 0.628 ± 0.0849 | 0.751 ± 0.0117 |
| JOAOv2 | 0.798 ± 0.0151 | 0.726 ± 0.0774 | 0.732 ± 0.0157 | 0.652 ± 0.0736 | 0.796 ± 0.0077 |
| PosPred | 0.766 ± 0.0157 | 0.701 ± 0.0467 | 0.693 ± 0.0202 | 0.615 ± 0.0553 | 0.763 ± 0.00844 |
| 3D-EMGP | 0.59 ± 0.0419 | 0.547 ± 0.054 | 0.556 ± 0.0253 | 0.52 ± 0.0559 | 0.662 ± 0.0502 |
| 3D-InfoMax | 0.837 ± 0.011 | 0.72 ± 0.0878 | 0.802 ± 0.0269 | 0.607 ± 0.093 | 0.839 ± 0.00759 |
| AttrMask | 0.729 ± 0.0258 | 0.743 ± 0.0796 | 0.668 ± 0.0253 | 0.646 ± 0.0755 | 0.743 ± 0.0168 |
| EdgePred | 0.628 ± 0.00892 | 0.51 ± 0.0244 | 0.539 ± 0.0189 | 0.571 ± 0.0702 | 0.643 ± 0.0157 |
| GraphMVP | 0.5 ± 0 | 0.513 ± 0.0283 | 0.5 ± 0.00382 | 0.5 ± 0 | 0.5 ± 0 |
| MolCLR-GCN | 0.499 ± 0.00209 | 0.552 ± 0.0619 | 0.487 ± 0.0136 | 0.49 ± 0.0169 | 0.496 ± 0.00784 |
| MolCLR-GIN | 0.501 ± 0.00282 | 0.456 ± 0.0421 | 0.484 ± 0.0165 | 0.54 ± 0.0813 | 0.503 ± 0.00666 |

Prediction is evaluated by ROC-AUC. For each method, the mean and standard deviation are reported for the evaluation metrics.

**Supplementary Table 14.**
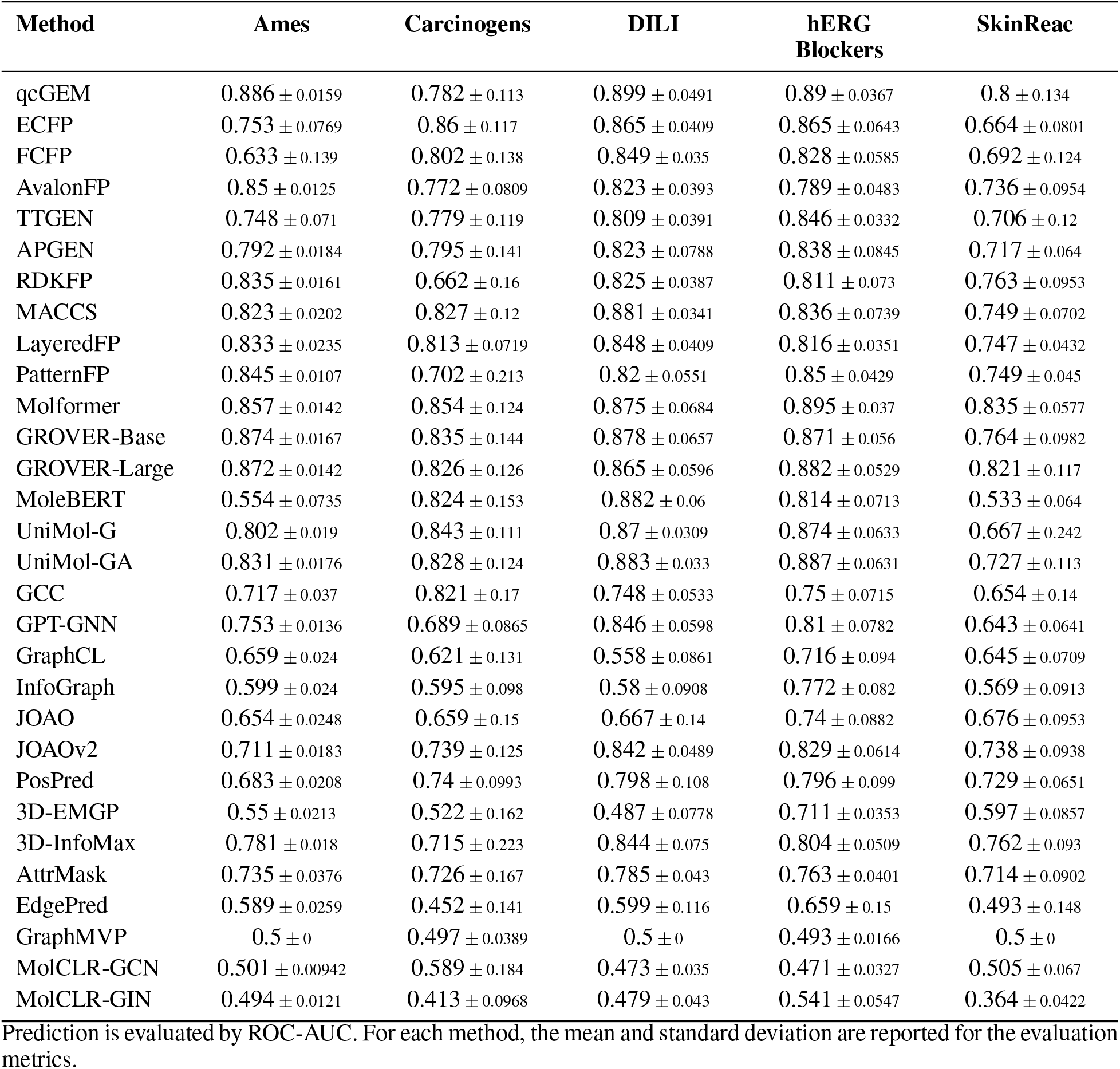
Molecular property classification task (scaffold split) – Part IV.

| Method | Ames | Carcinogens | DILI | hERG Blockers | SkinReac |
| --- | --- | --- | --- | --- | --- |
| qcGEM | 0.886 ± 0.0159 | 0.782 ± 0.113 | 0.899 ± 0.0491 | 0.89 ± 0.0367 | 0.8 ± 0.134 |
| ECFP | 0.753 ± 0.0769 | 0.86 ± 0.117 | 0.865 ± 0.0409 | 0.865 ± 0.0643 | 0.664 ± 0.0801 |
| FCFP | 0.633 ± 0.139 | 0.802 ± 0.138 | 0.849 ± 0.035 | 0.828 ± 0.0585 | 0.692 ± 0.124 |
| AvalonFP | 0.85 ± 0.0125 | 0.772 ± 0.0809 | 0.823 ± 0.0393 | 0.789 ± 0.0483 | 0.736 ± 0.0954 |
| TTGEN | 0.748 ± 0.071 | 0.779 ± 0.119 | 0.809 ± 0.0391 | 0.846 ± 0.0332 | 0.706 ± 0.12 |
| APGEN | 0.792 ± 0.0184 | 0.795 ± 0.141 | 0.823 ± 0.0788 | 0.838 ± 0.0845 | 0.717 ± 0.064 |
| RDKFP | 0.835 ± 0.0161 | 0.662 ± 0.16 | 0.825 ± 0.0387 | 0.811 ± 0.073 | 0.763 ± 0.0953 |
| MACCS | 0.823 ± 0.0202 | 0.827 ± 0.12 | 0.881 ± 0.0341 | 0.836 ± 0.0739 | 0.749 ± 0.0702 |
| LayeredFP | 0.833 ± 0.0235 | 0.813 ± 0.0719 | 0.848 ± 0.0409 | 0.816 ± 0.0351 | 0.747 ± 0.0432 |
| PatternFP | 0.845 ± 0.0107 | 0.702 ± 0.213 | 0.82 ± 0.0551 | 0.85 ± 0.0429 | 0.749 ± 0.045 |
| Molformer | 0.857 ± 0.0142 | 0.854 ± 0.124 | 0.875 ± 0.0684 | 0.895 ± 0.037 | 0.835 ± 0.0577 |
| GROVER-Base | 0.874 ± 0.0167 | 0.835 ± 0.144 | 0.878 ± 0.0657 | 0.871 ± 0.056 | 0.764 ± 0.0982 |
| GROVER-Large | 0.872 ± 0.0142 | 0.826 ± 0.126 | 0.865 ± 0.0596 | 0.882 ± 0.0529 | 0.821 ± 0.117 |
| MoleBERT | 0.554 ± 0.0735 | 0.824 ± 0.153 | 0.882 ± 0.06 | 0.814 ± 0.0713 | 0.533 ± 0.064 |
| UniMol-G | 0.802 ± 0.019 | 0.843 ± 0.111 | 0.87 ± 0.0309 | 0.874 ± 0.0633 | 0.667 ± 0.242 |
| UniMol-GA | 0.831 ± 0.0176 | 0.828 ± 0.124 | 0.883 ± 0.033 | 0.887 ± 0.0631 | 0.727 ± 0.113 |
| GCC | 0.717 ± 0.037 | 0.821 ± 0.17 | 0.748 ± 0.0533 | 0.75 ± 0.0715 | 0.654 ± 0.14 |
| GPT-GNN | 0.753 ± 0.0136 | 0.689 ± 0.0865 | 0.846 ± 0.0598 | 0.81 ± 0.0782 | 0.643 ± 0.0641 |
| GraphCL | 0.659 ± 0.024 | 0.621 ± 0.131 | 0.558 ± 0.0861 | 0.716 ± 0.094 | 0.645 ± 0.0709 |
| InfoGraph | 0.599 ± 0.024 | 0.595 ± 0.098 | 0.58 ± 0.0908 | 0.772 ± 0.082 | 0.569 ± 0.0913 |
| JOAO | 0.654 ± 0.0248 | 0.659 ± 0.15 | 0.667 ± 0.14 | 0.74 ± 0.0882 | 0.676 ± 0.0953 |
| JOAOv2 | 0.711 ± 0.0183 | 0.739 ± 0.125 | 0.842 ± 0.0489 | 0.829 ± 0.0614 | 0.738 ± 0.0938 |
| PosPred | 0.683 ± 0.0208 | 0.74 ± 0.0993 | 0.798 ± 0.108 | 0.796 ± 0.099 | 0.729 ± 0.0651 |
| 3D-EMGP | 0.55 ± 0.0213 | 0.522 ± 0.162 | 0.487 ± 0.0778 | 0.711 ± 0.0353 | 0.597 ± 0.0857 |
| 3D-InfoMax | 0.781 ± 0.018 | 0.715 ± 0.223 | 0.844 ± 0.075 | 0.804 ± 0.0509 | 0.762 ± 0.093 |
| AttrMask | 0.735 ± 0.0376 | 0.726 ± 0.167 | 0.785 ± 0.043 | 0.763 ± 0.0401 | 0.714 ± 0.0902 |
| EdgePred | 0.589 ± 0.0259 | 0.452 ± 0.141 | 0.599 ± 0.116 | 0.659 ± 0.15 | 0.493 ± 0.148 |
| GraphMVP | 0.5 ± 0 | 0.497 ± 0.0389 | 0.5 ± 0 | 0.493 ± 0.0166 | 0.5 ± 0 |
| MolCLR-GCN | 0.501 ± 0.00942 | 0.589 ± 0.184 | 0.473 ± 0.035 | 0.471 ± 0.0327 | 0.505 ± 0.067 |
| MolCLR-GIN | 0.494 ± 0.0121 | 0.413 ± 0.0968 | 0.479 ± 0.043 | 0.541 ± 0.0547 | 0.364 ± 0.0422 |

Prediction is evaluated by ROC-AUC. For each method, the mean and standard deviation are reported for the evaluation metrics.

**Supplementary Table 15.**
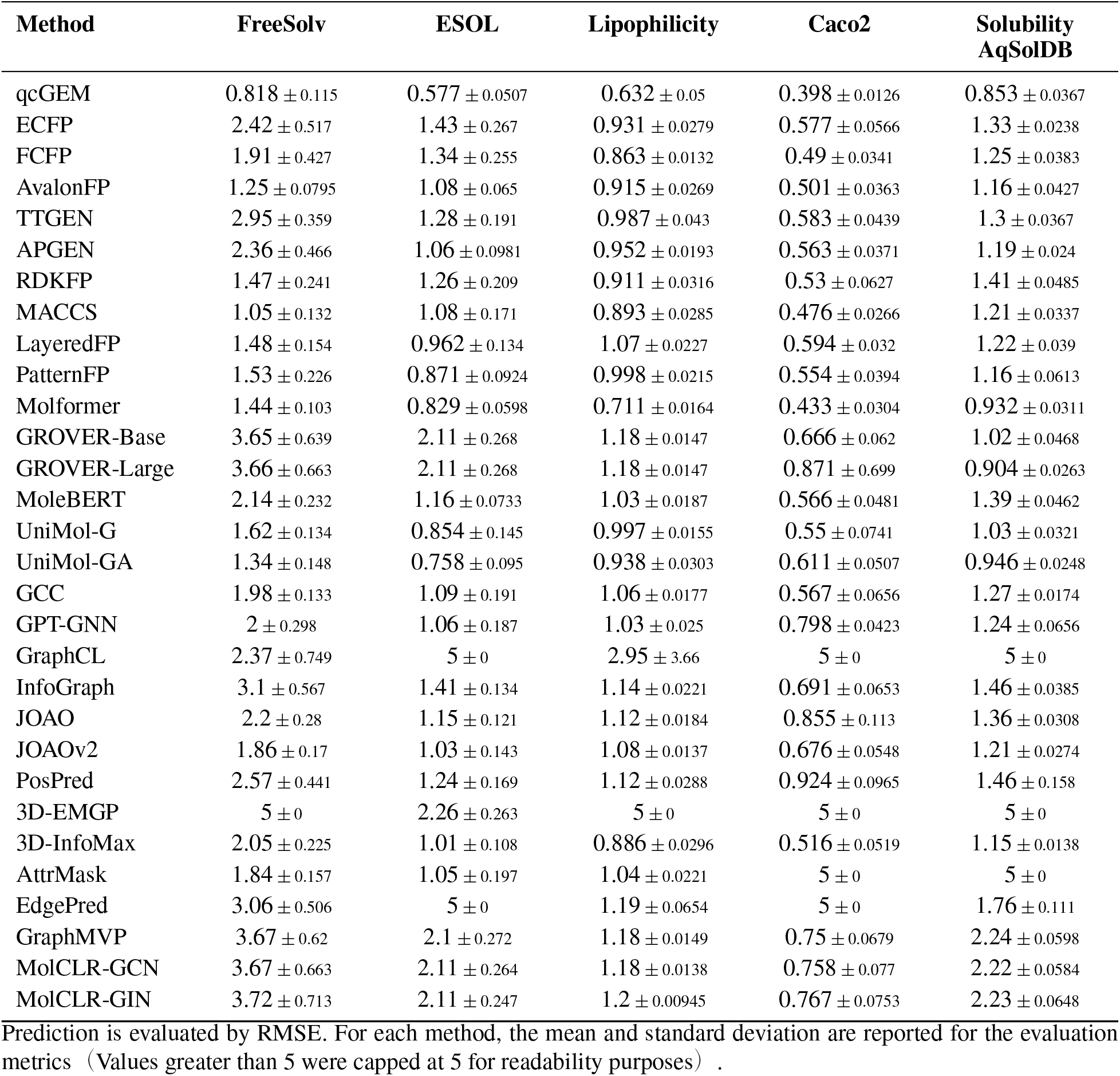
Molecular property regression task (random split) – Part I.

**Supplementary Table 16.** Molecular property regression task (random split) – Part II.

| Method | VDss | Clearance<br>Hepatocyte | Clearance<br>Microsome | Half Life | LD50 |
| --- | --- | --- | --- | --- | --- |
| qcGEM | 0.007 ± 0.00213 | 0.275 ± 0.0125 | 0.254 ± 0.0202 | 0.0649 ± 0.0738 | 0.57 ± 0.0121 |
| ECFP | 0.00728 ± 0.00221 | 0.295 ± 0.0134 | 0.258 ± 0.0293 | 0.0651 ± 0.0742 | 0.673 ± 0.0148 |
| FCFP | 0.00715 ± 0.00211 | 0.303 ± 0.0157 | 0.249 ± 0.0342 | 0.0653 ± 0.0742 | 0.67 ± 0.0172 |
| AvalonFP | 0.00707 ± 0.00227 | 0.304 ± 0.0144 | 0.258 ± 0.0169 | 0.0655 ± 0.0746 | 0.593 ± 0.0081 |
| TTGEN | 0.00756 ± 0.00191 | 0.308 ± 0.0125 | 0.259 ± 0.0125 | 0.0655 ± 0.0742 | 0.724 ± 0.0255 |
| APGEN | 0.00736 ± 0.00198 | 0.306 ± 0.015 | 0.264 ± 0.0173 | 0.0655 ± 0.0745 | 0.663 ± 0.0172 |
| RDKFP | 0.00732 ± 0.00204 | 0.307 ± 0.00383 | 0.257 ± 0.0146 | 0.0655 ± 0.0745 | 0.622 ± 0.0186 |
| MACCS | 0.00717 ± 0.00223 | 0.303 ± 0.0176 | 0.252 ± 0.017 | 0.0658 ± 0.0745 | 0.649 ± 0.0191 |
| LayeredFP | 0.00769 ± 0.00217 | 0.305 ± 0.0224 | 0.261 ± 0.0139 | 0.0658 ± 0.0744 | 0.637 ± 0.0229 |
| PatternFP | 0.0078 ± 0.00201 | 0.304 ± 0.00746 | 0.262 ± 0.0171 | 0.0654 ± 0.0745 | 0.653 ± 0.0125 |
| Molformer | 0.00714 ± 0.00211 | 0.277 ± 0.0177 | 0.237 ± 0.0154 | 0.0654 ± 0.0742 | 0.617 ± 0.00927 |
| GROVER-Base | 0.00681 ± 0.00197 | 0.264 ± 0.0241 | 0.237 ± 0.0193 | 0.0649 ± 0.0724 | 0.691 ± 0.0583 |
| GROVER-Large | 0.00753 ± 0.00178 | 0.28 ± 0.0274 | 0.238 ± 0.0195 | 0.0661 ± 0.074 | 0.594 ± 0.0205 |
| MoleBERT | 0.00751 ± 0.00239 | 0.312 ± 0.0109 | 0.273 ± 0.0152 | 0.0659 ± 0.0746 | 0.731 ± 0.0233 |
| UniMol-G | 0.00962 ± 0.00437 | 0.307 ± 0.017 | 0.255 ± 0.0137 | 0.0719 ± 0.0714 | 0.714 ± 0.0349 |
| UniMol-GA | 0.0109 ± 0.00399 | 0.307 ± 0.0157 | 0.255 ± 0.0138 | 0.0717 ± 0.0714 | 0.687 ± 0.0383 |
| GCC | 0.00767 ± 0.00221 | 0.315 ± 0.0102 | 0.273 ± 0.0154 | 0.066 ± 0.0746 | 0.728 ± 0.0174 |
| GPT-GNN | 0.00878 ± 0.00271 | 0.32 ± 0.00457 | 0.26 ± 0.0153 | 0.0666 ± 0.0737 | 0.705 ± 0.0202 |
| GraphCL | 5 ± 0 | 5 ± 0 | 5 ± 0 | 5 ± 0 | 0.894 ± 0.0611 |
| InfoGraph | 0.00768 ± 0.00233 | 0.328 ± 0.0104 | 0.265 ± 0.018 | 0.0667 ± 0.0744 | 0.865 ± 0.0356 |
| JOAO | 0.0194 ± 0.012 | 0.323 ± 0.00676 | 0.272 ± 0.0142 | 0.169 ± 0.201 | 0.774 ± 0.0404 |
| JOAOv2 | 0.00816 ± 0.00245 | 0.32 ± 0.00724 | 0.27 ± 0.0157 | 0.0688 ± 0.0756 | 0.755 ± 0.0317 |
| PosPred | 0.186 ± 0.239 | 0.32 ± 0.0089 | 0.352 ± 0.151 | 0.193 ± 0.141 | 0.825 ± 0.0693 |
| 3D-EMGP | 5 ± 0 | 5 ± 0 | 5 ± 0 | 5 ± 0 | 5 ± 0 |
| 3D-InfoMax | 0.00734 ± 0.00215 | 0.309 ± 0.0144 | 0.247 ± 0.0209 | 0.0654 ± 0.0747 | 0.718 ± 0.0124 |
| AttrMask | 0.00771 ± 0.00225 | 0.324 ± 0.013 | 0.268 ± 0.0148 | 5 ± 0 | 0.764 ± 0.0424 |
| EdgePred | 5 ± 0 | 5 ± 0 | 5 ± 0 | 5 ± 0 | 0.911 ± 0.0471 |
| GraphMVP | 0.00777 ± 0.00236 | 0.327 ± 0.0132 | 0.274 ± 0.0153 | 0.0659 ± 0.0746 | 0.915 ± 0.0321 |
| MolCLR-GCN | 0.0078 ± 0.00228 | 0.328 ± 0.015 | 0.275 ± 0.015 | 0.0659 ± 0.0745 | 0.92 ± 0.0305 |
| MolCLR-GIN | 0.00795 ± 0.00223 | 0.327 ± 0.0153 | 0.274 ± 0.0157 | 0.0659 ± 0.0745 | 0.918 ± 0.0306 |

Prediction is evaluated by RMSE. For each method, the mean and standard deviation are reported for the evaluation metrics (Values greater than 5 were capped at 5 for readability purposes) .

**Supplementary Table 17.** Molecular property classification task (random split) – Part I.

| Method | BBBP | BACE | Bioavailability | HIA | PAMPA |
| --- | --- | --- | --- | --- | --- |
| qcGEM | 0.907 ± 0.0239 | 0.848 ± 0.0614 | 0.707 ± 0.0638 | 0.936 ± 0.059 | 0.718 ± 0.0596 |
| ECFP | 0.848 ± 0.0246 | 0.864 ± 0.044 | 0.664 ± 0.0539 | 0.806 ± 0.079 | 0.695 ± 0.0317 |
| FCFP | 0.874 ± 0.031 | 0.871 ± 0.0327 | 0.659 ± 0.0879 | 0.817 ± 0.0386 | 0.715 ± 0.04 |
| AvalonFP | 0.87 ± 0.0265 | 0.845 ± 0.0274 | 0.698 ± 0.11 | 0.837 ± 0.0882 | 0.658 ± 0.0333 |
| TTGEN | 0.839 ± 0.0172 | 0.82 ± 0.0387 | 0.683 ± 0.0109 | 0.789 ± 0.0738 | 0.646 ± 0.0381 |
| APGEN | 0.858 ± 0.0211 | 0.855 ± 0.0201 | 0.675 ± 0.0136 | 0.854 ± 0.0922 | 0.673 ± 0.039 |
| RDKFP | 0.872 ± 0.0396 | 0.857 ± 0.0246 | 0.637 ± 0.0614 | 0.817 ± 0.138 | 0.673 ± 0.0876 |
| MACCS | 0.865 ± 0.0392 | 0.841 ± 0.0299 | 0.688 ± 0.0616 | 0.938 ± 0.0482 | 0.712 ± 0.0352 |
| LayeredFP | 0.858 ± 0.0284 | 0.83 ± 0.0317 | 0.684 ± 0.0494 | 0.862 ± 0.0344 | 0.652 ± 0.018 |
| PatternFP | 0.867 ± 0.0169 | 0.859 ± 0.0217 | 0.631 ± 0.0584 | 0.76 ± 0.186 | 0.638 ± 0.0479 |
| Molformer | 0.885 ± 0.0254 | 0.862 ± 0.0257 | 0.804 ± 0.0394 | 0.964 ± 0.0173 | 0.737 ± 0.0373 |
| GROVER-Base | 0.5 ± 0 | 0.5 ± 0 | 0.741 ± 0.0611 | 0.956 ± 0.0393 | 0.724 ± 0.046 |
| GROVER-Large | 0.5 ± 0 | 0.5 ± 0 | 0.778 ± 0.0518 | 0.924 ± 0.0763 | 0.717 ± 0.0354 |
| MoleBERT | 0.831 ± 0.0271 | 0.488 ± 0.0884 | 0.649 ± 0.102 | 0.898 ± 0.0843 | 0.636 ± 0.0198 |
| UniMol-G | 0.847 ± 0.0296 | 0.778 ± 0.0649 | 0.646 ± 0.165 | 0.952 ± 0.018 | 0.67 ± 0.0235 |
| UniMol-GA | 0.854 ± 0.0295 | 0.799 ± 0.0576 | 0.645 ± 0.137 | 0.955 ± 0.0383 | 0.673 ± 0.0396 |
| GCC | 0.792 ± 0.0318 | 0.709 ± 0.0482 | 0.684 ± 0.0317 | 0.845 ± 0.0221 | 0.616 ± 0.0398 |
| GPT-GNN | 0.799 ± 0.0323 | 0.749 ± 0.0366 | 0.635 ± 0.061 | 0.687 ± 0.167 | 0.595 ± 0.066 |
| GraphCL | 0.455 ± 0.0774 | 0.536 ± 0.0668 | 0.574 ± 0.108 | 0.51 ± 0.138 | 0.468 ± 0.0456 |
| InfoGraph | 0.747 ± 0.0486 | 0.626 ± 0.0249 | 0.606 ± 0.135 | 0.744 ± 0.212 | 0.527 ± 0.0173 |
| JOAO | 0.746 ± 0.0469 | 0.716 ± 0.036 | 0.647 ± 0.0548 | 0.657 ± 0.215 | 0.564 ± 0.0758 |
| JOAOv2 | 0.793 ± 0.0291 | 0.734 ± 0.0362 | 0.686 ± 0.0542 | 0.792 ± 0.154 | 0.561 ± 0.0488 |
| PosPred | 0.735 ± 0.0456 | 0.757 ± 0.0428 | 0.679 ± 0.0606 | 0.718 ± 0.11 | 0.546 ± 0.0497 |
| 3D-EMGP | 0.465 ± 0.0221 | 0.533 ± 0.0629 | 0.588 ± 0.0874 | 0.389 ± 0.181 | 0.493 ± 0.0357 |
| 3D-InfoMax | 0.844 ± 0.0179 | 0.838 ± 0.032 | 0.674 ± 0.115 | 0.862 ± 0.1 | 0.698 ± 0.048 |
| AttrMask | 0.734 ± 0.038 | 0.718 ± 0.0512 | 0.661 ± 0.0678 | 0.808 ± 0.148 | 0.582 ± 0.0835 |
| EdgePred | 0.526 ± 0.173 | 0.474 ± 0.134 | 0.556 ± 0.0973 | 0.497 ± 0.265 | 0.447 ± 0.0722 |
| GraphMVP | 0.497 ± 0.0114 | 0.5 ± 0 | 0.506 ± 0.0087 | 0.539 ± 0.0809 | 0.521 ± 0.0392 |
| MolCLR-GCN | 0.506 ± 0.0242 | 0.532 ± 0.0475 | 0.52 ± 0.0675 | 0.599 ± 0.17 | 0.473 ± 0.0323 |
| MolCLR-GIN | 0.518 ± 0.0496 | 0.518 ± 0.029 | 0.573 ± 0.118 | 0.517 ± 0.0513 | 0.523 ± 0.0318 |

Prediction is evaluated by ROC-AUC. For each method, the mean and standard deviation are reported for the evaluation metrics.

**Supplementary Table 18.** Molecular property classification task (random split) – Part II.

| Method | Pgp | BBB | CYP1A2 Veith | CYP2C19 Veith | CYP2C9 Substrate |
| --- | --- | --- | --- | --- | --- |
| qcGEM | 0.95 ± 0.00496 | 0.893 ± 0.0248 | 0.928 ± 0.0112 | 0.888 ± 0.00595 | 0.768 ± 0.0564 |
| ECFP | 0.915 ± 0.0314 | 0.865 ± 0.0219 | 0.857 ± 0.00784 | 0.658 ± 0.144 | 0.678 ± 0.093 |
| FCFP | 0.92 ± 0.0206 | 0.871 ± 0.0176 | 0.869 ± 0.0115 | 0.726 ± 0.127 | 0.615 ± 0.103 |
| AvalonFP | 0.904 ± 0.0255 | 0.892 ± 0.0203 | 0.885 ± 0.00945 | 0.844 ± 0.00896 | 0.599 ± 0.166 |
| TTGEN | 0.902 ± 0.0158 | 0.819 ± 0.0256 | 0.86 ± 0.00829 | 0.681 ± 0.104 | 0.693 ± 0.137 |
| APGEN | 0.919 ± 0.0195 | 0.862 ± 0.0174 | 0.887 ± 0.00822 | 0.819 ± 0.0135 | 0.651 ± 0.074 |
| RDKFP | 0.89 ± 0.0267 | 0.889 ± 0.0256 | 0.893 ± 0.00527 | 0.825 ± 0.0125 | 0.68 ± 0.126 |
| MACCS | 0.904 ± 0.0257 | 0.894 ± 0.0227 | 0.872 ± 0.0098 | 0.828 ± 0.00663 | 0.633 ± 0.124 |
| LayeredFP | 0.901 ± 0.0229 | 0.854 ± 0.0153 | 0.858 ± 0.00861 | 0.787 ± 0.009 | 0.585 ± 0.12 |
| PatternFP | 0.909 ± 0.0243 | 0.868 ± 0.0135 | 0.886 ± 0.0107 | 0.821 ± 0.00396 | 0.637 ± 0.109 |
| Molformer | 0.932 ± 0.019 | 0.892 ± 0.0157 | 0.924 ± 0.0117 | 0.883 ± 0.0046 | 0.69 ± 0.0636 |
| GROVER-Base | 0.942 ± 0.0247 | 0.891 ± 0.0238 | 0.927 ± 0.00967 | 0.886 ± 0.00743 | 0.713 ± 0.085 |
| GROVER-Large | 0.941 ± 0.0212 | 0.885 ± 0.0272 | 0.924 ± 0.00861 | 0.885 ± 0.00407 | 0.732 ± 0.0697 |
| MoleBERT | 0.553 ± 0.0968 | 0.86 ± 0.0241 | 0.784 ± 0.159 | 0.53 ± 0.0666 | 0.678 ± 0.0661 |
| UniMol-G | 0.912 ± 0.0192 | 0.839 ± 0.0232 | 0.89 ± 0.00231 | 0.828 ± 0.00661 | 0.575 ± 0.103 |
| UniMol-GA | 0.93 ± 0.011 | 0.86 ± 0.021 | 0.899 ± 0.5 | 0.838 ± 0.5 | 0.572 ± 0.106 |
| GCC | 0.879 ± 0.0209 | 0.779 ± 0.0391 | 0.848 ± 0.00713 | 0.778 ± 0.00964 | 0.677 ± 0.0605 |
| GPT-GNN | 0.895 ± 0.0229 | 0.786 ± 0.0272 | 0.861 ± 0.00752 | 0.786 ± 0.0124 | 0.65 ± 0.0693 |
| GraphCL | 0.713 ± 0.0895 | 0.475 ± 0.0403 | 0.82 ± 0.0133 | 0.701 ± 0.011 | 0.536 ± 0.0649 |
| InfoGraph | 0.878 ± 0.0293 | 0.716 ± 0.0239 | 0.808 ± 0.00999 | 0.671 ± 0.0125 | 0.617 ± 0.0331 |
| JOAO | 0.874 ± 0.0203 | 0.697 ± 0.0409 | 0.838 ± 0.00414 | 0.722 ± 0.00671 | 0.549 ± 0.101 |
| JOAOv2 | 0.884 ± 0.0222 | 0.768 ± 0.0447 | 0.868 ± 0.00489 | 0.786 ± 0.0103 | 0.648 ± 0.0405 |
| PosPred | 0.881 ± 0.0259 | 0.705 ± 0.0613 | 0.867 ± 0.00447 | 0.746 ± 0.0111 | 0.635 ± 0.0679 |
| 3D-EMGP | 0.778 ± 0.0366 | 0.547 ± 0.0471 | 0.702 ± 0.076 | 0.64 ± 0.0129 | 0.552 ± 0.0709 |
| 3D-InfoMax | 0.921 ± 0.025 | 0.852 ± 0.0248 | 0.886 ± 0.00657 | 0.826 ± 0.00723 | 0.585 ± 0.128 |
| AttrMask | 0.908 ± 0.0105 | 0.71 ± 0.0552 | 0.827 ± 0.016 | 0.711 ± 0.0287 | 0.682 ± 0.0829 |
| EdgePred | 0.584 ± 0.337 | 0.517 ± 0.156 | 0.582 ± 0.068 | 0.565 ± 0.0228 | 0.535 ± 0.0698 |
| GraphMVP | 0.5 ± 0 | 0.491 ± 0.0231 | 0.5 ± 0 | 0.5 ± 0 | 0.525 ± 0.0345 |
| MolCLR-GCN | 0.521 ± 0.0456 | 0.495 ± 0.045 | 0.51 ± 0.0136 | 0.502 ± 0.0084 | 0.552 ± 0.085 |
| MolCLR-GIN | 0.515 ± 0.0296 | 0.559 ± 0.087 | 0.503 ± 0.0092 | 0.503 ± 0.00531 | 0.478 ± 0.0257 |

Prediction is evaluated by ROC-AUC. For each method, the mean and standard deviation are reported for the evaluation metrics.

**Supplementary Table 19.** Molecular property classification task (random split) – Part III.

| Method | CYP2C9 Veith | CYP2D6 Substrate | CYP2D6 Veith | CYP3A4 Substrate | CYP3A4 Veith |
| --- | --- | --- | --- | --- | --- |
| qcGEM | 0.885 ± 0.00408 | 0.797 ± 0.048 | 0.861 ± 0.0123 | 0.679 ± 0.0809 | 0.886 ± 0.00578 |
| ECFP | 0.835 ± 0.00561 | 0.751 ± 0.0183 | 0.792 ± 0.011 | 0.644 ± 0.0893 | 0.675 ± 0.162 |
| FCFP | 0.837 ± 0.00816 | 0.738 ± 0.0225 | 0.807 ± 0.00929 | 0.584 ± 0.131 | 0.691 ± 0.177 |
| AvalonFP | 0.841 ± 0.00527 | 0.733 ± 0.1 | 0.804 ± 0.0223 | 0.642 ± 0.0729 | 0.842 ± 0.0105 |
| TTGEN | 0.829 ± 0.0066 | 0.74 ± 0.0574 | 0.775 ± 0.0211 | 0.602 ± 0.0981 | 0.6 ± 0.138 |
| APGEN | 0.824 ± 0.0138 | 0.739 ± 0.039 | 0.783 ± 0.0174 | 0.649 ± 0.0956 | 0.825 ± 0.0128 |
| RDKFP | 0.844 ± 0.00839 | 0.786 ± 0.0475 | 0.804 ± 0.00954 | 0.646 ± 0.0941 | 0.835 ± 0.0146 |
| MACCS | 0.829 ± 0.00575 | 0.772 ± 0.0438 | 0.798 ± 0.0166 | 0.618 ± 0.11 | 0.822 ± 0.00687 |
| LayeredFP | 0.805 ± 0.00918 | 0.727 ± 0.0607 | 0.773 ± 0.0159 | 0.597 ± 0.0605 | 0.789 ± 0.0108 |
| PatternFP | 0.824 ± 0.00401 | 0.747 ± 0.0637 | 0.789 ± 0.0142 | 0.659 ± 0.101 | 0.823 ± 0.00641 |
| Molformer | 0.88 ± 0.00734 | 0.78 ± 0.0475 | 0.853 ± 0.013 | 0.627 ± 0.113 | 0.884 ± 0.00856 |
| GROVER-Base | 0.885 ± 0.00548 | 0.778 ± 0.0642 | 0.866 ± 0.0136 | 0.613 ± 0.0763 | 0.885 ± 0.00623 |
| GROVER-Large | 0.881 ± 0.0073 | 0.767 ± 0.0628 | 0.86 ± 0.00846 | 0.647 ± 0.0727 | 0.879 ± 0.00893 |
| MoleBERT | 0.811 ± 0.00619 | 0.745 ± 0.0473 | 0.767 ± 0.0199 | 0.456 ± 0.0564 | 0.616 ± 0.131 |
| UniMol-G | 0.836 ± 0.00933 | 0.76 ± 0.0394 | 0.782 ± 0.015 | 0.643 ± 0.0976 | 0.825 ± 0.0148 |
| UniMol-GA | 0.847 ± 0.00662 | 0.768 ± 0.039 | 0.799 ± 0.0151 | 0.657 ± 0.0903 | 0.849 ± 0.0107 |
| GCC | 0.797 ± 0.00834 | 0.683 ± 0.0665 | 0.731 ± 0.021 | 0.593 ± 0.0533 | 0.75 ± 0.0843 |
| GPT-GNN | 0.79 ± 0.0147 | 0.593 ± 0.1 | 0.744 ± 0.0176 | 0.638 ± 0.101 | 0.779 ± 0.0091 |
| GraphCL | 0.711 ± 0.0181 | 0.626 ± 0.0851 | 0.613 ± 0.00807 | 0.614 ± 0.0827 | 0.718 ± 0.0169 |
| InfoGraph | 0.681 ± 0.0165 | 0.707 ± 0.0842 | 0.658 ± 0.0133 | 0.619 ± 0.0923 | 0.718 ± 0.0103 |
| JOAO | 0.742 ± 0.0135 | 0.646 ± 0.0699 | 0.6 ± 0.0296 | 0.605 ± 0.0808 | 0.744 ± 0.0144 |
| JOAOv2 | 0.791 ± 0.0137 | 0.704 ± 0.045 | 0.72 ± 0.018 | 0.637 ± 0.0905 | 0.789 ± 0.0127 |
| PosPred | 0.77 ± 0.0161 | 0.675 ± 0.0548 | 0.697 ± 0.0186 | 0.605 ± 0.116 | 0.763 ± 0.00725 |
| 3D-EMGP | 0.597 ± 0.0268 | 0.512 ± 0.0785 | 0.54 ± 0.0357 | 0.627 ± 0.0604 | 0.663 ± 0.0565 |
| 3D-InfoMax | 0.84 ± 0.00934 | 0.713 ± 0.0488 | 0.785 ± 0.0178 | 0.572 ± 0.101 | 0.837 ± 0.0084 |
| AttrMask | 0.711 ± 0.00968 | 0.789 ± 0.0433 | 0.699 ± 0.0593 | 0.588 ± 0.0861 | 0.73 ± 0.0153 |
| EdgePred | 0.628 ± 0.0263 | 0.482 ± 0.0464 | 0.549 ± 0.0253 | 0.538 ± 0.0905 | 0.627 ± 0.0145 |
| GraphMVP | 0.5 ± 0 | 0.495 ± 0.0111 | 0.496 ± 0.00983 | 0.5 ± 0 | 0.5 ± 0 |
| MolCLR-GCN | 0.502 ± 0.0101 | 0.458 ± 0.078 | 0.499 ± 0.00142 | 0.444 ± 0.0774 | 0.497 ± 0.00417 |
| MolCLR-GIN | 0.502 ± 0.00515 | 0.552 ± 0.0989 | 0.501 ± 0.0025 | 0.531 ± 0.0525 | 0.498 ± 0.0175 |

Prediction is evaluated by ROC-AUC. For each method, the mean and standard deviation are reported for the evaluation metrics.

**Supplementary Table 20.** Molecular property classification task (random split) – Part IV.

| Method | Ames | Carcinogens | DILI | hERG Blockers | SkinReac |
| --- | --- | --- | --- | --- | --- |
| qcGEM | 0.896 ± 0.0226 | 0.802 ± 0.129 | 0.937 ± 0.00873 | 0.863 ± 0.0952 | 0.741 ± 0.0805 |
| ECFP | 0.683 ± 0.168 | 0.708 ± 0.245 | 0.869 ± 0.0423 | 0.847 ± 0.0607 | 0.739 ± 0.0564 |
| FCFP | 0.566 ± 0.146 | 0.745 ± 0.0851 | 0.875 ± 0.0251 | 0.829 ± 0.0417 | 0.681 ± 0.063 |
| AvalonFP | 0.86 ± 0.0199 | 0.734 ± 0.133 | 0.896 ± 0.027 | 0.798 ± 0.0742 | 0.7 ± 0.112 |
| TTGEN | 0.766 ± 0.0581 | 0.731 ± 0.133 | 0.863 ± 0.0254 | 0.798 ± 0.0573 | 0.621 ± 0.13 |
| APGEN | 0.804 ± 0.0119 | 0.717 ± 0.183 | 0.87 ± 0.0224 | 0.839 ± 0.0372 | 0.718 ± 0.104 |
| RDKFP | 0.844 ± 0.0148 | 0.733 ± 0.0898 | 0.911 ± 0.0291 | 0.817 ± 0.0899 | 0.779 ± 0.0787 |
| MACCS | 0.83 ± 0.0206 | 0.763 ± 0.103 | 0.929 ± 0.00777 | 0.843 ± 0.0737 | 0.756 ± 0.0423 |
| LayeredFP | 0.837 ± 0.0164 | 0.752 ± 0.134 | 0.898 ± 0.0445 | 0.815 ± 0.0733 | 0.732 ± 0.121 |
| PatternFP | 0.854 ± 0.0216 | 0.736 ± 0.153 | 0.864 ± 0.02 | 0.837 ± 0.0124 | 0.744 ± 0.113 |
| Molformer | 0.871 ± 0.0212 | 0.742 ± 0.226 | 0.914 ± 0.0139 | 0.87 ± 0.081 | 0.804 ± 0.0723 |
| GROVER-Base | 0.883 ± 0.0177 | 0.827 ± 0.139 | 0.922 ± 0.0215 | 0.855 ± 0.11 | 0.785 ± 0.0508 |
| GROVER-Large | 0.889 ± 0.0138 | 0.806 ± 0.187 | 0.911 ± 0.0276 | 0.868 ± 0.105 | 0.753 ± 0.0213 |
| MoleBERT | 0.5 ± 0 | 0.809 ± 0.116 | 0.923 ± 0.0329 | 0.806 ± 0.0965 | 0.569 ± 0.0634 |
| UniMol-G | 0.816 ± 0.0185 | 0.835 ± 0.0916 | 0.876 ± 0.0551 | 0.823 ± 0.0475 | 0.687 ± 0.0892 |
| UniMol-GA | 0.843 ± 0.0216 | 0.831 ± 0.12 | 0.883 ± 0.0681 | 0.844 ± 0.0436 | 0.694 ± 0.0824 |
| GCC | 0.74 ± 0.0136 | 0.796 ± 0.12 | 0.836 ± 0.0563 | 0.769 ± 0.065 | 0.59 ± 0.0881 |
| GPT-GNN | 0.765 ± 0.0226 | 0.766 ± 0.13 | 0.825 ± 0.0912 | 0.822 ± 0.065 | 0.587 ± 0.0972 |
| GraphCL | 0.678 ± 0.0298 | 0.54 ± 0.203 | 0.68 ± 0.122 | 0.721 ± 0.0782 | 0.622 ± 0.16 |
| InfoGraph | 0.642 ± 0.0101 | 0.552 ± 0.139 | 0.56 ± 0.0629 | 0.795 ± 0.0205 | 0.655 ± 0.0643 |
| JOAO | 0.692 ± 0.0105 | 0.641 ± 0.197 | 0.749 ± 0.104 | 0.789 ± 0.108 | 0.666 ± 0.138 |
| JOAOv2 | 0.733 ± 0.0159 | 0.64 ± 0.16 | 0.915 ± 0.0305 | 0.83 ± 0.0793 | 0.649 ± 0.0982 |
| PosPred | 0.713 ± 0.00632 | 0.685 ± 0.181 | 0.826 ± 0.0342 | 0.815 ± 0.0775 | 0.665 ± 0.0845 |
| 3D-EMGP | 0.566 ± 0.0435 | 0.563 ± 0.123 | 0.417 ± 0.0637 | 0.713 ± 0.0896 | 0.508 ± 0.14 |
| 3D-InfoMax | 0.798 ± 0.0219 | 0.744 ± 0.133 | 0.918 ± 0.0503 | 0.802 ± 0.112 | 0.718 ± 0.0372 |
| AttrMask | 0.754 ± 0.0202 | 0.787 ± 0.18 | 0.842 ± 0.0224 | 0.742 ± 0.119 | 0.544 ± 0.166 |
| EdgePred | 0.586 ± 0.0293 | 0.607 ± 0.245 | 0.633 ± 0.105 | 0.671 ± 0.156 | 0.59 ± 0.0649 |
| GraphMVP | 0.5 ± 0 | 0.503 ± 0.0645 | 0.5 ± 0 | 0.495 ± 0.0109 | 0.527 ± 0.0614 |
| MolCLR-GCN | 0.5 ± 0 | 0.638 ± 0.0921 | 0.525 ± 0.0455 | 0.468 ± 0.0893 | 0.456 ± 0.0979 |
| MolCLR-GIN | 0.5 ± 0 | 0.432 ± 0.135 | 0.495 ± 0.0478 | 0.516 ± 0.0439 | 0.513 ± 0.137 |

Prediction is evaluated by ROC-AUC. For each method, the mean and standard deviation are reported for the evaluation metrics.

##### 3.3.2 Activity cliff prediction

**Supplementary Figure 9.**
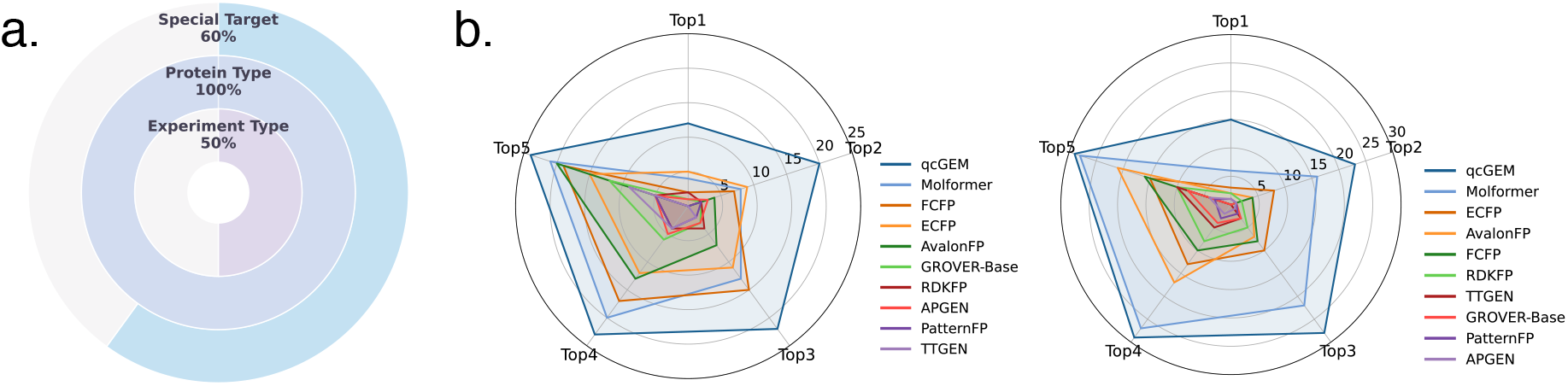
Ranking evaluation of qcGEM in activity cliff tasks. **a)** This figure supplements Figure 3d, with focus shifted to activity cliff molecules instead of all molecules. Specifically, the multi-layer pie chart illustrates proportions of qcGEM achieving the best ranking for activity cliff molecules in 30 tasks under three task classification criteria: individual proteins targets (outer), protein classes (middle), and experimental categories (inner). **b)** The radar chart on the left shows the frequency with which each method appears in the top 1-5 rankings for all molecules, while the chart on the right shows the results for activity cliff molecules.

**Supplementary Figure 10.**
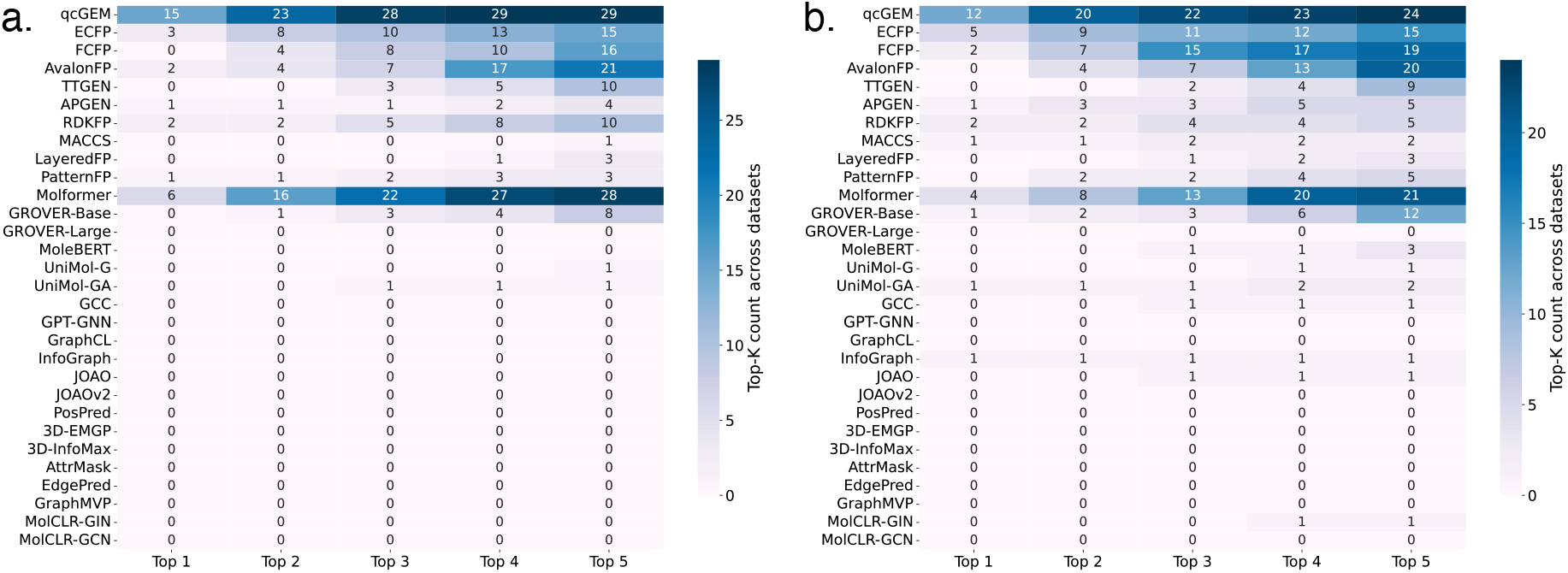
Statistics of the top 1-5 rankings for all methods in activity cliff tasks. **a)** Evaluation results of qcGEM and baseline methods on all molecules. **b)** Evaluation results of qcGEM and baseline methods on activity cliff molecules.

**Supplementary Figure 11.**
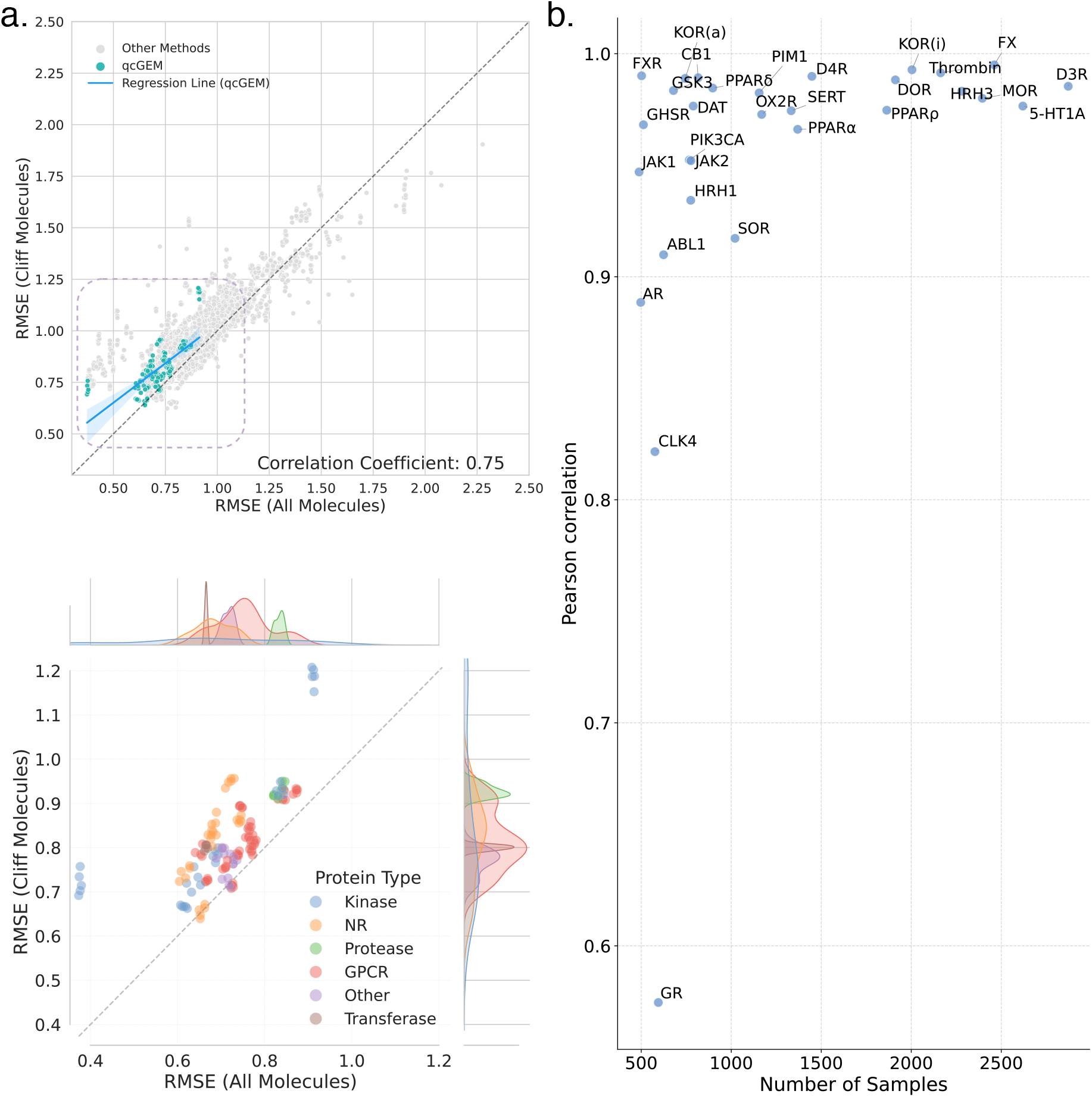
Additional results of qcGEM in activity cliff tasks. **a)** The upper scatter plot compares the prediction error (in RMSE) on all molecules (horizontal axis) vs. activity cliff molecules (vertical axis). Each point refers to a protein target. qcGEM results are highlighted in teal, while the other methods are shown in gray, with the dashed diagonal indicating equal performance between the two subsets for ideal methods with perfect generalizability. A fitted regression line for qcGEM (blue) demonstrates the positive correlation (Pearson correlation coefficient *r* = 0.75) between the two groups of molecules. The lower scatter plot (corresponding to the dash-line enclosed region in the upper plot) provides a detailed view of qcGEM results. Here, protein targets are colored by their protein classes: kinase, NR, protease, GPCR, transferase, and other. Kernel density estimates are shown along the margins to highlight the distribution of RMSE values. **b)** Correlation between inference values of models optimized on all and activity cliff molecules. Each point represents one protein target, with the horizontal and vertical axes showing the number of available samples and the Pearson correlation coefficient, respectively.

**Supplementary Figure 12.**
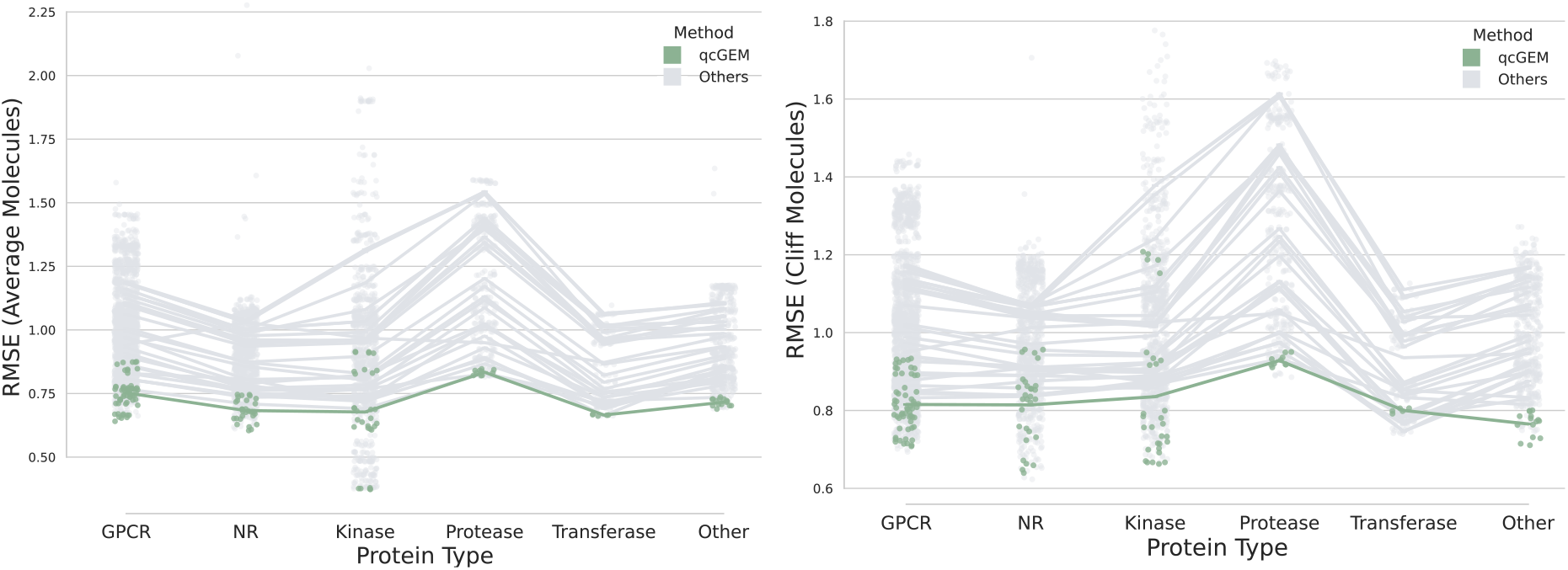
Evaluation of qcGEM vs. other methods among the 6 protein types in the activity cliff prediction. The prediction error (in RMSE) is compared for qcGEM (colored in green) and other methods (colored in gray) for all molecules and activity cliff molecules in the left and right panels, respectively. For each protein type, results on individual protein targets are presented as dots, whereas the mean results are connected by solid lines across protein types.

**Supplementary Figure 13.**
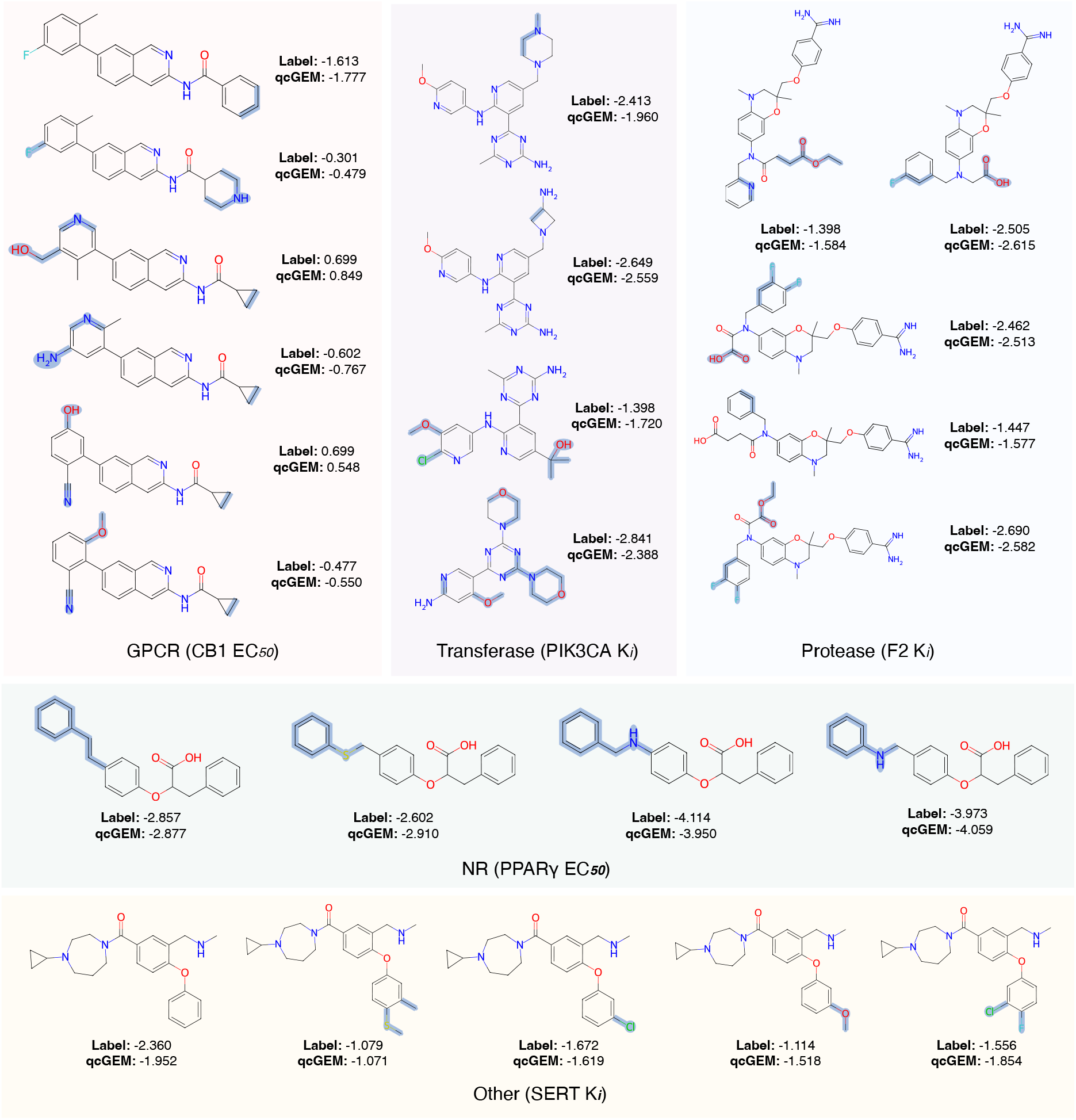
Examples of the identification of activity cliff molecules for different kinds of protein targets. As a supplement to the kinase target in Figure 3e, this figure presents activity cliff molecules for five protein targets falling in the other protein classes (GPCR, protease, NR, transferase, and other). Among ligands targeted to the same protein, difference in chemical structure is highlighted in blue shadow for each ligand, while the ground-truth binding affinity and prediction based on qcGEM are listed aside.

**Supplementary Table 21.**
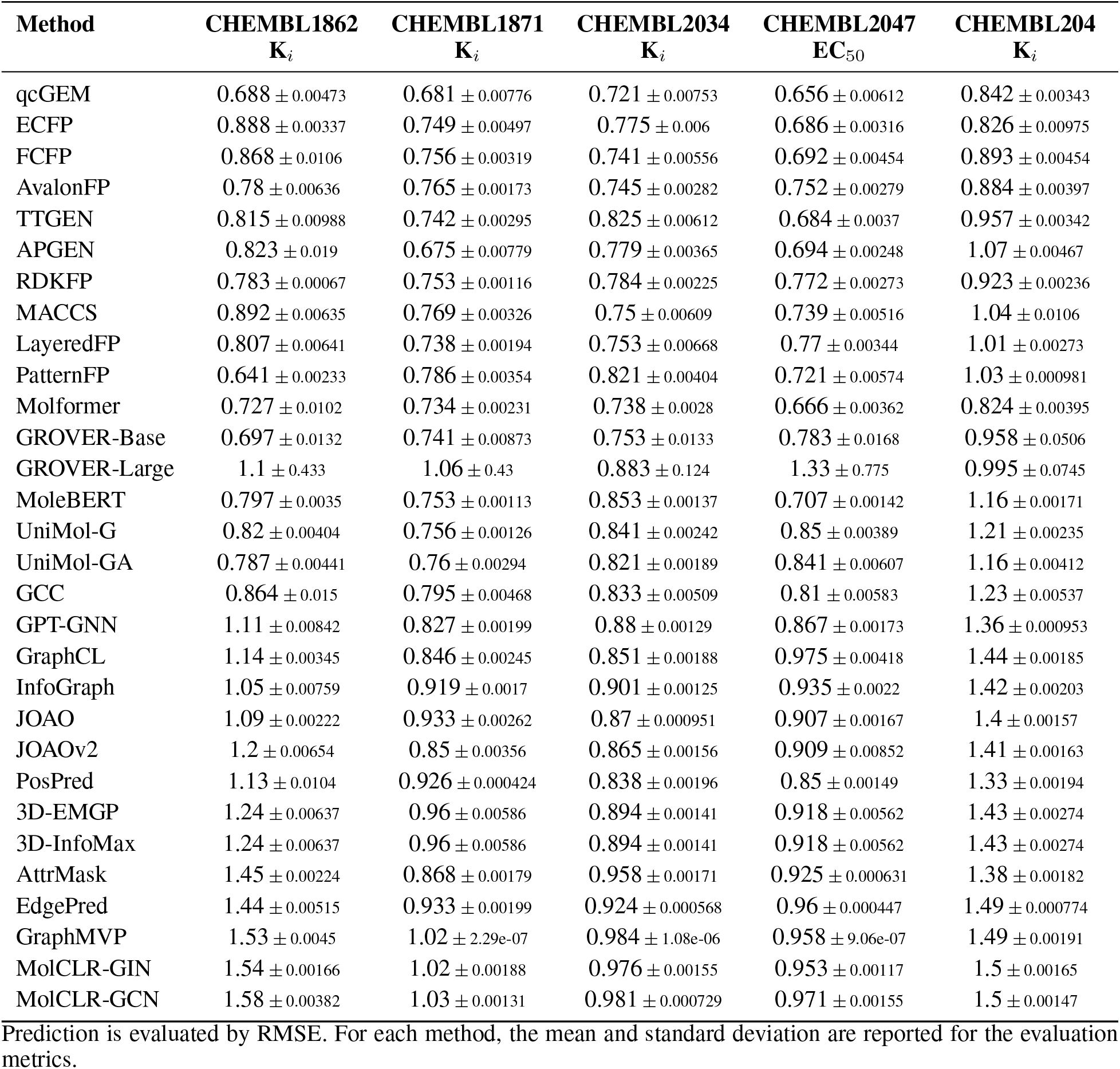
Activity cliff prediction (all molecules) – Part I.

**Supplementary Table 22.** Activity cliff prediction (all molecules) – Part II.

| Method | CHEMBL2147<br>$K_i$ | CHEMBL214<br>$K_i$ | CHEMBL218<br>$EC_{50}$ | CHEMBL219<br>$K_i$ | CHEMBL228<br>$K_i$ |
| --- | --- | --- | --- | --- | --- |
| qcGEM | 0.615 ± 0.00602 | 0.745 ± 0.00297 | 0.766 ± 0.00773 | 0.774 ± 0.00587 | 0.728 ± 0.00518 |
| ECFP | 0.744 ± 0.01 | 0.733 ± 0.00376 | 0.874 ± 0.00304 | 0.872 ± 0.00115 | 0.812 ± 0.00265 |
| FCFP | 0.676 ± 0.00556 | 0.755 ± 0.00211 | 0.836 ± 0.00279 | 0.862 ± 0.00328 | 0.859 ± 0.00727 |
| AvalonFP | 0.711 ± 0.00149 | 0.714 ± 0.00198 | 0.779 ± 0.00435 | 0.854 ± 0.00345 | 0.841 ± 0.0033 |
| TTGEN | 0.717 ± 0.00587 | 0.825 ± 0.00441 | 0.803 ± 0.00515 | 0.848 ± 0.00514 | 0.896 ± 0.00113 |
| APGEN | 0.837 ± 0.0065 | 0.843 ± 0.00519 | 0.877 ± 0.00471 | 0.89 ± 0.00255 | 0.915 ± 0.0059 |
| RDKFP | 0.683 ± 0.0039 | 0.805 ± 0.00284 | 0.732 ± 0.00491 | 0.832 ± 0.0053 | 0.875 ± 0.00298 |
| MACCS | 0.913 ± 0.00596 | 0.889 ± 0.00321 | 0.834 ± 0.00548 | 0.958 ± 0.00233 | 0.921 ± 0.00625 |
| LayeredFP | 0.813 ± 0.00444 | 0.875 ± 0.00377 | 0.827 ± 0.0006 | 0.915 ± 0.000919 | 0.855 ± 0.00244 |
| PatternFP | 0.772 ± 0.00529 | 0.873 ± 0.00262 | 0.78 ± 0.00183 | 0.873 ± 0.000734 | 0.905 ± 0.00264 |
| Molformer | 0.685 ± 0.0033 | 0.735 ± 0.0039 | 0.782 ± 0.0024 | 0.803 ± 0.00478 | 0.771 ± 0.00428 |
| GROVER-Base | 0.707 ± 0.016 | 0.774 ± 0.018 | 0.851 ± 0.0348 | 0.88 ± 0.0239 | 0.859 ± 0.0276 |
| GROVER-Large | 0.88 ± 0.169 | 0.864 ± 0.077 | 1.06 ± 0.304 | 0.967 ± 0.0478 | 0.932 ± 0.0523 |
| MoleBERT | 0.921 ± 0.00395 | 0.945 ± 0.00184 | 0.914 ± 0.00197 | 0.941 ± 0.0015 | 0.975 ± 0.00168 |
| UniMol-G | 0.968 ± 0.00276 | 0.951 ± 0.00195 | 0.927 ± 0.000354 | 1.03 ± 0.000899 | 1.02 ± 0.00433 |
| UniMol-GA | 0.911 ± 0.00616 | 0.927 ± 0.00213 | 0.923 ± 0.00342 | 1.02 ± 0.001 | 0.98 ± 0.00278 |
| GCC | 1.05 ± 0.00766 | 0.984 ± 0.00275 | 0.989 ± 0.0053 | 0.983 ± 0.00121 | 1.03 ± 0.00537 |
| GPT-GNN | 1.17 ± 0.00126 | 1.05 ± 0.000554 | 0.989 ± 0.000663 | 1.02 ± 0.00185 | 1.12 ± 0.00206 |
| GraphCL | 1.18 ± 0.00243 | 1.04 ± 0.00102 | 0.971 ± 0.00241 | 1.01 ± 0.000673 | 1.11 ± 0.00211 |
| InfoGraph | 1.38 ± 0.00547 | 1.09 ± 0.00193 | 0.992 ± 0.000772 | 1.07 ± 0.00107 | 1.17 ± 0.00333 |
| JOAO | 1.23 ± 0.00553 | 1.05 ± 0.00039 | 0.971 ± 0.000552 | 1.03 ± 0.000423 | 1.15 ± 0.00166 |
| JOAOv2 | 1.14 ± 0.00514 | 1.05 ± 0.000847 | 0.979 ± 0.000666 | 1.01 ± 0.000746 | 1.11 ± 0.00386 |
| PosPred | 1.21 ± 0.00463 | 1.03 ± 0.000576 | 0.958 ± 0.000836 | 0.989 ± 0.00134 | 1.14 ± 0.00145 |
| 3D-EMGP | 1.36 ± 0.0116 | 1.07 ± 0.00134 | 1 ± 0.00507 | 1.06 ± 0.00377 | 1.13 ± 0.00499 |
| 3D-InfoMax | 1.36 ± 0.0116 | 1.07 ± 0.00134 | 1 ± 0.00507 | 1.06 ± 0.00377 | 1.13 ± 0.00499 |
| AttrMask | 1.24 ± 0.00276 | 1.06 ± 0.000992 | 1.01 ± 0.00085 | 1.06 ± 0.000442 | 1.17 ± 0.000269 |
| EdgePred | 1.69 ± 0.00181 | 1.07 ± 0.000468 | 0.987 ± 0.000212 | 1.06 ± 0.000372 | 1.15 ± 0.00177 |
| GraphMVP | 1.9 ± 0.00473 | 1.1 ± 8.51e-07 | 1.03 ± 3.65e-06 | 1.07 ± 2.88e-06 | 1.17 ± 3.07e-07 |
| MolCLR-GIN | 1.9 ± 0.00425 | 1.1 ± 0.000224 | 1.02 ± 0.00233 | 1.08 ± 0.00158 | 1.17 ± 0.000672 |
| MolCLR-GCN | 1.9 ± 0.000866 | 1.1 ± 0.000164 | 1.03 ± 0.00222 | 1.08 ± 0.000208 | 1.17 ± 0.00068 |

Prediction is evaluated by RMSE. For each method, the mean and standard deviation are reported for the evaluation metrics.

**Supplementary Table 23.** Activity cliff prediction (all molecules) – Part III.

| Method | CHEMBL231<br>$K_i$ | CHEMBL233<br>$K_i$ | CHEMBL234<br>$K_i$ | CHEMBL235<br>$EC_{50}$ | CHEMBL236<br>$K_i$ |
| --- | --- | --- | --- | --- | --- |
| qcGEM | 0.654 ± 0.00873 | 0.872 ± 0.0039 | 0.726 ± 0.00194 | 0.681 ± 0.00479 | 0.766 ± 0.00297 |
| ECFP | 0.932 ± 0.0091 | 0.894 ± 0.00274 | 0.715 ± 0.00498 | 0.761 ± 0.00293 | 0.758 ± 0.0033 |
| FCFP | 0.82 ± 0.00521 | 0.935 ± 0.00593 | 0.746 ± 0.00288 | 0.765 ± 0.00378 | 0.813 ± 0.00325 |
| AvalonFP | 0.915 ± 0.00846 | 0.881 ± 0.00247 | 0.727 ± 0.00328 | 0.754 ± 0.00254 | 0.802 ± 0.00111 |
| TTGEN | 0.812 ± 0.00506 | 0.91 ± 0.00357 | 0.74 ± 0.00243 | 0.796 ± 0.00311 | 0.841 ± 0.000559 |
| APGEN | 0.816 ± 0.00503 | 0.993 ± 0.00323 | 0.806 ± 0.00497 | 0.78 ± 0.00914 | 0.922 ± 0.00399 |
| RDKFP | 0.877 ± 0.0037 | 0.977 ± 0.00186 | 0.782 ± 0.00195 | 0.723 ± 0.00213 | 0.916 ± 0.00305 |
| MACCS | 0.956 ± 0.00426 | 1.07 ± 0.00242 | 0.858 ± 0.00755 | 0.871 ± 0.00597 | 1 ± 0.0063 |
| LayeredFP | 0.85 ± 0.00513 | 1.03 ± 0.000597 | 0.833 ± 0.00268 | 0.805 ± 0.00489 | 0.959 ± 0.00367 |
| PatternFP | 0.837 ± 0.00825 | 1.02 ± 0.00132 | 0.867 ± 0.000343 | 0.8 ± 0.00247 | 0.933 ± 0.00477 |
| Molformer | 0.772 ± 0.00571 | 0.882 ± 0.00155 | 0.712 ± 0.00208 | 0.654 ± 0.00383 | 0.812 ± 0.00299 |
| GROVER-Base | 0.819 ± 0.0394 | 0.914 ± 0.0244 | 0.767 ± 0.0119 | 0.821 ± 0.0314 | 0.846 ± 0.021 |
| GROVER-Large | 1.11 ± 0.388 | 0.921 ± 0.0258 | 0.776 ± 0.00738 | 0.841 ± 0.0524 | 0.905 ± 0.0813 |
| MoleBERT | 0.986 ± 0.00307 | 1.1 ± 0.000863 | 0.897 ± 0.000958 | 0.9 ± 0.00179 | 1 ± 0.00102 |
| UniMol-G | 0.949 ± 0.00337 | 1.09 ± 0.0016 | 0.92 ± 0.000983 | 0.934 ± 0.00225 | 1.05 ± 0.00301 |
| UniMol-GA | 0.893 ± 0.00384 | 1.06 ± 0.00178 | 0.9 ± 0.00131 | 0.909 ± 0.00587 | 1.03 ± 0.00243 |
| GCC | 0.993 ± 0.00518 | 1.11 ± 0.00245 | 0.922 ± 0.000947 | 0.931 ± 0.00249 | 1.06 ± 0.00593 |
| GPT-GNN | 1.07 ± 0.00197 | 1.21 ± 0.000852 | 0.974 ± 0.0011 | 1.03 ± 0.00113 | 1.26 ± 0.00165 |
| GraphCL | 1.07 ± 0.00262 | 1.26 ± 0.00118 | 0.967 ± 0.0008 | 1.01 ± 0.00308 | 1.29 ± 0.00186 |
| InfoGraph | 1.14 ± 0.000782 | 1.22 ± 0.00126 | 1.03 ± 0.00131 | 1.05 ± 0.00116 | 1.3 ± 0.00329 |
| JOAO | 1.14 ± 0.00132 | 1.25 ± 0.000532 | 0.979 ± 0.000409 | 1.02 ± 0.000637 | 1.27 ± 0.00138 |
| JOAOv2 | 1.15 ± 0.0016 | 1.23 ± 0.000983 | 0.938 ± 0.000478 | 1.01 ± 0.000723 | 1.26 ± 0.00261 |
| PosPred | 1.09 ± 0.00372 | 1.17 ± 0.00143 | 0.941 ± 0.000843 | 0.995 ± 0.00207 | 1.17 ± 0.00412 |
| 3D-EMGP | 1.23 ± 0.00413 | 1.24 ± 0.00327 | 1 ± 0.00109 | 1.05 ± 0.00113 | 1.31 ± 0.00222 |
| 3D-InfoMax | 1.23 ± 0.00413 | 1.24 ± 0.00327 | 1 ± 0.00109 | 1.05 ± 0.00113 | 1.31 ± 0.00222 |
| AttrMask | 1.24 ± 0.00805 | 1.26 ± 0.00176 | 1.04 ± 0.00105 | 1.06 ± 0.00135 | 1.3 ± 0.00174 |
| EdgePred | 1.22 ± 0.0019 | 1.27 ± 0.000401 | 1.07 ± 0.000195 | 1.05 ± 5.96e-05 | 1.32 ± 0.00041 |
| GraphMVP | 1.33 ± 5.72e-08 | 1.28 ± 1.79e-06 | 1.1 ± 2.88e-06 | 1.07 ± 6.04e-06 | 1.33 ± 1.94e-06 |
| MolCLR-GIN | 1.33 ± 0.00182 | 1.28 ± 0.000152 | 1.1 ± 0.000176 | 1.06 ± 0.000736 | 1.33 ± 0.00117 |
| MolCLR-GCN | 1.34 ± 0.00121 | 1.28 ± 9.05e-05 | 1.1 ± 0.000296 | 1.06 ± 0.000239 | 1.33 ± 0.000609 |

Prediction is evaluated by RMSE. For each method, the mean and standard deviation are reported for the evaluation metrics.

**Supplementary Table 24.** Activity cliff prediction (all molecules) – Part IV.

| Method | CHEMBL237<br>EC<sub>50</sub> | CHEMBL237<br>K<sub>i</sub> | CHEMBL238<br>K<sub>i</sub> | CHEMBL239<br>EC<sub>50</sub> | CHEMBL244<br>K<sub>i</sub> |
| --- | --- | --- | --- | --- | --- |
| qcGEM | 0.841 ± 0.0066 | 0.739 ± 0.00247 | 0.702 ± 0.00748 | 0.742 ± 0.00383 | 0.825 ± 0.00596 |
| ECFP | 0.823 ± 0.00636 | 0.791 ± 0.00513 | 0.805 ± 0.004 | 0.85 ± 0.00278 | 0.827 ± 0.00506 |
| FCFP | 0.88 ± 0.00263 | 0.788 ± 0.00361 | 0.853 ± 0.00391 | 0.789 ± 0.00258 | 0.877 ± 0.00398 |
| AvalonFP | 0.9 ± 0.00466 | 0.818 ± 0.00299 | 0.765 ± 0.00493 | 0.764 ± 0.00531 | 0.849 ± 0.00603 |
| TTGEN | 0.974 ± 0.00782 | 0.799 ± 0.00187 | 0.762 ± 0.00431 | 0.864 ± 0.00321 | 0.994 ± 0.00494 |
| APGEN | 0.965 ± 0.00461 | 0.884 ± 0.00706 | 0.811 ± 0.00915 | 0.909 ± 0.00208 | 1.1 ± 0.00635 |
| RDKFP | 1 ± 0.00221 | 0.954 ± 0.00234 | 0.789 ± 0.00972 | 0.828 ± 0.00361 | 0.859 ± 0.00606 |
| MACCS | 1.02 ± 0.00674 | 0.974 ± 0.00637 | 0.817 ± 0.00247 | 0.886 ± 0.00811 | 1 ± 0.0125 |
| LayeredFP | 1.09 ± 0.00271 | 0.906 ± 0.000763 | 0.783 ± 0.00617 | 0.859 ± 0.0026 | 1.02 ± 0.00483 |
| PatternFP | 1.11 ± 0.00264 | 0.958 ± 0.000242 | 0.814 ± 0.00189 | 0.852 ± 0.00318 | 1.01 ± 0.00864 |
| Molformer | 0.838 ± 0.00751 | 0.798 ± 0.00374 | 0.739 ± 0.00502 | 0.818 ± 0.00109 | 0.838 ± 0.00139 |
| GROVER-Base | 0.905 ± 0.0362 | 0.828 ± 0.021 | 0.813 ± 0.0202 | 0.918 ± 0.0293 | 0.901 ± 0.0118 |
| GROVER-Large | 1.2 ± 0.227 | 0.905 ± 0.0856 | 1.15 ± 0.4 | 0.988 ± 0.0879 | 0.905 ± 0.0262 |
| MoleBERT | 1.16 ± 0.00357 | 1.02 ± 0.0019 | 0.909 ± 0.00216 | 0.892 ± 0.000968 | 1.1 ± 0.00157 |
| UniMol-G | 1.24 ± 0.00169 | 1.05 ± 0.00391 | 0.965 ± 0.00291 | 0.93 ± 0.000748 | 1.13 ± 0.00197 |
| UniMol-GA | 1.19 ± 0.00347 | 1 ± 0.00517 | 0.927 ± 0.00204 | 0.911 ± 0.00139 | 1.07 ± 0.00355 |
| GCC | 1.18 ± 0.00751 | 1.02 ± 0.00829 | 0.955 ± 0.00525 | 0.942 ± 0.00199 | 1.18 ± 0.00362 |
| GPT-GNN | 1.39 ± 0.00312 | 1.18 ± 0.000853 | 1.08 ± 0.00168 | 1.03 ± 0.000444 | 1.33 ± 0.00198 |
| GraphCL | 1.33 ± 0.00131 | 1.24 ± 0.00106 | 1.08 ± 0.0032 | 1.02 ± 0.000658 | 1.41 ± 0.000787 |
| InfoGraph | 1.32 ± 0.0018 | 1.2 ± 0.000648 | 1.17 ± 0.00334 | 1.03 ± 0.00167 | 1.4 ± 0.00219 |
| JOAO | 1.37 ± 0.000605 | 1.2 ± 0.000857 | 1.1 ± 0.00188 | 1.01 ± 0.000268 | 1.41 ± 0.00105 |
| JOAOv2 | 1.35 ± 0.003 | 1.27 ± 0.000806 | 1.04 ± 0.00202 | 1.01 ± 0.000243 | 1.32 ± 0.00543 |
| PosPred | 1.31 ± 0.00378 | 1.14 ± 0.000963 | 1.07 ± 0.00328 | 0.999 ± 0.000696 | 1.32 ± 0.00398 |
| 3D-EMGP | 1.35 ± 0.0028 | 1.26 ± 0.00227 | 1.08 ± 0.0047 | 1.07 ± 0.000832 | 1.45 ± 0.002 |
| 3D-InfoMax | 1.35 ± 0.0028 | 1.26 ± 0.00227 | 1.08 ± 0.0047 | 1.07 ± 0.000832 | 1.45 ± 0.002 |
| AttrMask | 1.38 ± 0.0041 | 1.22 ± 0.000405 | 1.16 ± 0.000727 | 1.05 ± 0.00127 | 1.41 ± 0.00152 |
| EdgePred | 1.44 ± 0.000722 | 1.31 ± 0.00102 | 1.11 ± 0.00313 | 1.06 ± 0.000141 | 1.58 ± 0.000153 |
| GraphMVP | 1.45 ± 0.000153 | 1.33 ± 3.15e-07 | 1.16 ± 7.24e-07 | 1.08 ± 1.23e-06 | 1.58 ± 1.23e-07 |
| MolCLR-GIN | 1.45 ± 0.000512 | 1.33 ± 0.000268 | 1.16 ± 0.00147 | 1.08 ± 0.00204 | 1.59 ± 0.000403 |
| MolCLR-GCN | 1.45 ± 0.0012 | 1.33 ± 0.000197 | 1.17 ± 0.00144 | 1.09 ± 0.000883 | 1.59 ± 0.000157 |

Prediction is evaluated by RMSE. For each method, the mean and standard deviation are reported for the evaluation metrics.

**Supplementary Table 25.** Activity cliff prediction (all molecules) – Part V.

| Method | CHEMBL262<br>$K_i$ | CHEMBL264<br>$K_i$ | CHEMBL2835<br>$K_i$ | CHEMBL287<br>$K_i$ | CHEMBL2971<br>$K_i$ |
| --- | --- | --- | --- | --- | --- |
| qcGEM | 0.836 ± 0.00693 | 0.667 ± 0.0025 | 0.376 ± 0.00235 | 0.718 ± 0.00986 | 0.638 ± 0.0125 |
| ECFP | 0.86 ± 0.0058 | 0.733 ± 0.0035 | 0.407 ± 0.00335 | 0.733 ± 0.00702 | 0.719 ± 0.00549 |
| FCFP | 0.787 ± 0.00381 | 0.728 ± 0.00211 | 0.378 ± 0.00205 | 0.753 ± 0.00412 | 0.759 ± 0.00629 |
| AvalonFP | 0.805 ± 0.000678 | 0.725 ± 0.00351 | 0.431 ± 0.00251 | 0.825 ± 0.00671 | 0.636 ± 0.00238 |
| TTGEN | 0.893 ± 0.00333 | 0.773 ± 0.00198 | 0.43 ± 0.00353 | 0.768 ± 0.00296 | 0.675 ± 0.00673 |
| APGEN | 0.832 ± 0.00362 | 0.763 ± 0.00623 | 0.43 ± 0.00221 | 0.794 ± 0.00508 | 0.741 ± 0.00513 |
| RDKFP | 0.819 ± 0.0051 | 0.786 ± 0.00387 | 0.382 ± 0.00121 | 0.892 ± 0.0028 | 0.687 ± 0.00463 |
| MACCS | 0.922 ± 0.00279 | 0.866 ± 0.00777 | 0.456 ± 0.00334 | 0.843 ± 0.00462 | 0.73 ± 0.00219 |
| LayeredFP | 0.842 ± 0.00343 | 0.837 ± 0.00465 | 0.441 ± 0.00281 | 0.799 ± 0.00366 | 0.779 ± 0.00128 |
| PatternFP | 0.861 ± 0.00475 | 0.808 ± 0.00388 | 0.382 ± 0.00172 | 0.787 ± 0.0022 | 0.692 ± 0.00366 |
| Molformer | 0.756 ± 0.009 | 0.673 ± 0.00383 | 0.403 ± 0.0023 | 0.695 ± 0.00328 | 0.651 ± 0.00604 |
| GROVER-Base | 0.854 ± 0.0283 | 0.711 ± 0.0116 | 0.387 ± 0.00144 | 0.713 ± 0.0172 | 0.665 ± 0.0103 |
| GROVER-Large | 1.19 ± 0.43 | 0.737 ± 0.014 | 0.463 ± 0.115 | 0.837 ± 0.0991 | 0.817 ± 0.156 |
| MoleBERT | 0.903 ± 0.00362 | 0.871 ± 0.00096 | 0.45 ± 0.00298 | 0.776 ± 0.000439 | 0.832 ± 0.00296 |
| UniMol-G | 0.951 ± 0.000644 | 0.89 ± 0.00192 | 0.487 ± 0.00253 | 0.815 ± 0.00201 | 0.783 ± 0.00227 |
| UniMol-GA | 0.932 ± 0.00159 | 0.86 ± 0.00231 | 0.484 ± 0.00225 | 0.79 ± 0.00184 | 0.759 ± 0.00366 |
| GCC | 0.989 ± 0.00838 | 0.931 ± 0.00246 | 0.518 ± 0.00186 | 0.816 ± 0.0052 | 0.869 ± 0.00979 |
| GPT-GNN | 1.02 ± 0.00217 | 1.02 ± 0.00117 | 0.52 ± 0.00103 | 0.853 ± 0.000658 | 0.881 ± 0.00253 |
| GraphCL | 1.02 ± 0.00271 | 1.01 ± 0.000838 | 0.499 ± 0.000896 | 0.824 ± 0.000912 | 0.87 ± 0.00227 |
| InfoGraph | 1.02 ± 0.00411 | 1.04 ± 0.000788 | 0.563 ± 0.00154 | 0.911 ± 0.00215 | 0.914 ± 0.00166 |
| JOAO | 1.02 ± 0.000561 | 0.989 ± 0.000459 | 0.482 ± 0.00116 | 0.91 ± 0.00027 | 0.885 ± 0.00232 |
| JOAOv2 | 1.03 ± 0.00143 | 0.992 ± 0.00106 | 0.484 ± 0.00128 | 0.813 ± 0.0018 | 0.855 ± 0.00237 |
| PosPred | 1.01 ± 0.00145 | 1.01 ± 0.00025 | 0.496 ± 0.000835 | 0.846 ± 0.00149 | 0.813 ± 0.00194 |
| 3D-EMGP | 1.03 ± 0.00561 | 0.999 ± 0.00252 | 0.548 ± 0.00375 | 0.893 ± 0.00487 | 0.978 ± 0.00808 |
| 3D-InfoMax | 1.03 ± 0.00561 | 0.999 ± 0.00252 | 0.548 ± 0.00375 | 0.893 ± 0.00487 | 0.978 ± 0.00808 |
| AttrMask | 1.12 ± 0.000809 | 1.05 ± 0.00134 | 0.688 ± 0.111 | 0.92 ± 0.00343 | 0.901 ± 0.00442 |
| EdgePred | 1.05 ± 0.000559 | 1.05 ± 0.000425 | 0.706 ± 0.000365 | 0.918 ± 0.000527 | 1.18 ± 0.00125 |
| GraphMVP | 1.12 ± 0.000476 | 1.06 ± 0.000203 | 0.863 ± 0.000388 | 0.969 ± 0.000428 | 1.37 ± 0.000778 |
| MolCLR-GIN | 1.12 ± 0.00225 | 1.06 ± 0.000389 | 0.863 ± 6.61e-05 | 0.969 ± 0.000563 | 1.35 ± 0.000349 |
| MolCLR-GCN | 1.12 ± 0.000666 | 1.06 ± 0.000682 | 0.863 ± 5.07e-05 | 0.98 ± 0.000395 | 1.38 ± 0.00167 |

Prediction is evaluated by RMSE. For each method, the mean and standard deviation are reported for the evaluation metrics.

**Supplementary Table 26.** Activity cliff prediction (all molecules) – Part VI.

| Method | CHEMBL3979<br>EC<sub>50</sub> | CHEMBL4005<br>K<sub>i</sub> | CHEMBL4203<br>K<sub>i</sub> | CHEMBL4616<br>EC<sub>50</sub> | CHEMBL4792<br>K<sub>i</sub> |
| --- | --- | --- | --- | --- | --- |
| qcGEM | 0.618 ± 0.011 | 0.665 ± 0.00283 | 0.912 ± 0.00258 | 0.71 ± 0.00385 | 0.77 ± 0.00268 |
| ECFP | 0.62 ± 0.0055 | 0.701 ± 0.00476 | 0.988 ± 0.00868 | 0.673 ± 0.0076 | 0.881 ± 0.00266 |
| FCFP | 0.666 ± 0.00832 | 0.69 ± 0.00408 | 0.978 ± 0.0068 | 0.725 ± 0.0057 | 0.81 ± 0.0048 |
| AvalonFP | 0.671 ± 0.00985 | 0.714 ± 0.00346 | 0.954 ± 0.00592 | 0.744 ± 0.00136 | 0.83 ± 0.00355 |
| TTGEN | 0.66 ± 0.0176 | 0.7 ± 0.00317 | 1.06 ± 0.00855 | 0.792 ± 0.00303 | 0.848 ± 0.00274 |
| APGEN | 0.744 ± 0.00988 | 0.749 ± 0.00101 | 0.958 ± 0.00624 | 0.763 ± 0.00403 | 0.837 ± 0.0041 |
| RDKFP | 0.659 ± 0.0122 | 0.664 ± 0.00135 | 1.03 ± 0.00405 | 0.757 ± 0.00141 | 0.897 ± 0.00182 |
| MACCS | 0.719 ± 0.0163 | 0.734 ± 0.00186 | 1.02 ± 0.00296 | 0.762 ± 0.00134 | 0.977 ± 0.00464 |
| LayeredFP | 0.696 ± 0.0026 | 0.756 ± 0.00286 | 1.01 ± 0.00254 | 0.786 ± 0.00124 | 0.994 ± 0.00247 |
| PatternFP | 0.717 ± 0.00456 | 0.702 ± 0.00144 | 0.985 ± 0.00154 | 0.746 ± 0.00168 | 0.92 ± 0.00425 |
| Molformer | 0.646 ± 0.00559 | 0.722 ± 0.00183 | 0.919 ± 0.00372 | 0.686 ± 0.00587 | 0.714 ± 0.00445 |
| GROVER-Base | 0.825 ± 0.0226 | 0.736 ± 0.0214 | 0.972 ± 0.00543 | 0.769 ± 0.00459 | 0.888 ± 0.028 |
| GROVER-Large | 1.07 ± 0.302 | 0.872 ± 0.168 | 1.36 ± 0.54 | 0.872 ± 0.118 | 0.981 ± 0.0864 |
| MoleBERT | 0.724 ± 0.00719 | 0.768 ± 0.000703 | 0.976 ± 0.0021 | 0.861 ± 0.000484 | 0.982 ± 0.00302 |
| UniMol-G | 0.927 ± 0.00599 | 0.822 ± 0.00067 | 0.957 ± 0.00202 | 0.828 ± 0.00216 | 1 ± 0.00224 |
| UniMol-GA | 0.899 ± 0.00634 | 0.794 ± 0.00241 | 0.945 ± 0.00286 | 0.817 ± 0.00121 | 0.968 ± 0.00369 |
| GCC | 0.928 ± 0.0108 | 0.865 ± 0.00328 | 1.09 ± 0.00203 | 0.877 ± 0.00375 | 1.04 ± 0.00294 |
| GPT-GNN | 1.07 ± 0.00279 | 0.964 ± 0.00293 | 1.07 ± 0.00236 | 0.912 ± 0.000443 | 1.08 ± 0.000912 |
| GraphCL | 1 ± 0.00274 | 1.01 ± 0.00128 | 1.06 ± 0.00242 | 0.926 ± 0.00335 | 1.13 ± 0.00116 |
| InfoGraph | 1.02 ± 0.00454 | 0.965 ± 0.00218 | 1.03 ± 0.000971 | 0.906 ± 0.000861 | 1.16 ± 0.00131 |
| JOAO | 1.03 ± 0.00255 | 0.948 ± 0.00216 | 1.04 ± 0.00309 | 0.905 ± 0.000256 | 1.13 ± 0.000431 |
| JOAOv2 | 1.04 ± 0.00175 | 0.958 ± 0.00305 | 1.05 ± 0.000537 | 0.893 ± 0.000406 | 1.13 ± 0.00148 |
| PosPred | 1 ± 0.0053 | 0.944 ± 0.0031 | 1.04 ± 0.000788 | 0.902 ± 0.0014 | 1.03 ± 0.00127 |
| 3D-EMGP | 1.12 ± 0.00506 | 1.02 ± 0.00508 | 1.05 ± 0.00481 | 0.968 ± 0.0067 | 1.16 ± 0.00353 |
| 3D-InfoMax | 1.12 ± 0.00506 | 1.02 ± 0.00508 | 1.05 ± 0.00481 | 0.968 ± 0.0067 | 1.16 ± 0.00353 |
| AttrMask | 1.06 ± 0.00259 | 1.01 ± 0.00176 | 1.07 ± 0.000379 | 0.923 ± 0.000926 | 1.15 ± 0.00085 |
| EdgePred | 1.07 ± 0.00101 | 0.99 ± 0.000305 | 1.05 ± 0.00253 | 0.919 ± 0.000126 | 1.19 ± 0.000209 |
| GraphMVP | 1.08 ± 0.00165 | 1.06 ± 0.000885 | 1.07 ± 2.4e-07 | 0.928 ± 0.000181 | 1.18 ± 0.000216 |
| MolCLR-GIN | 1.08 ± 0.000864 | 1.06 ± 0.00187 | 1.07 ± 0.00351 | 0.93 ± 0.0023 | 1.19 ± 0.00162 |
| MolCLR-GCN | 1.1 ± 0.00496 | 1.06 ± 0.000597 | 1.06 ± 0.000595 | 0.939 ± 0.00157 | 1.19 ± 0.000584 |

Prediction is evaluated by RMSE. For each method, the mean and standard deviation are reported for the evaluation metrics.

**Supplementary Table 27.** Activity cliff prediction (cliff molecules) – Part I.

| Method | CHEMBL1862<br>$K_i$ | CHEMBL1871<br>$K_i$ | CHEMBL2034<br>$K_i$ | CHEMBL2047<br>$EC_{50}$ | CHEMBL204<br>$K_i$ |
| --- | --- | --- | --- | --- | --- |
| qcGEM | 0.785 ± 0.0126 | 0.851 ± 0.0193 | 0.949 ± 0.00888 | 0.656 ± 0.0127 | 0.94 ± 0.00975 |
| ECFP | 0.834 ± 0.0235 | 0.882 ± 0.0106 | 0.871 ± 0.00232 | 0.722 ± 0.00497 | 0.973 ± 0.01 |
| FCFP | 0.805 ± 0.00792 | 0.889 ± 0.0103 | 0.855 ± 0.0117 | 0.741 ± 0.00778 | 1.02 ± 0.00755 |
| AvalonFP | 0.829 ± 0.0108 | 1.06 ± 0.018 | 0.865 ± 0.0111 | 0.798 ± 0.00471 | 1.02 ± 0.00368 |
| TTGEN | 0.878 ± 0.00282 | 0.928 ± 0.0134 | 1.02 ± 0.0113 | 0.745 ± 0.00626 | 1.11 ± 0.00378 |
| APGEN | 0.802 ± 0.0165 | 0.906 ± 0.015 | 0.969 ± 0.0116 | 0.688 ± 0.00987 | 1.21 ± 0.00751 |
| RDKFP | 0.831 ± 0.00374 | 0.953 ± 0.00933 | 0.912 ± 0.00364 | 0.754 ± 0.0198 | 1.05 ± 0.00675 |
| MACCS | 0.899 ± 0.0125 | 0.97 ± 0.00392 | 0.859 ± 0.00189 | 0.722 ± 0.013 | 1.2 ± 0.016 |
| LayeredFP | 0.876 ± 0.00749 | 0.972 ± 0.0121 | 0.934 ± 0.00637 | 0.8 ± 0.00818 | 1.19 ± 0.00459 |
| PatternFP | 0.831 ± 0.0022 | 0.985 ± 0.00408 | 0.97 ± 0.00739 | 0.703 ± 0.00738 | 1.21 ± 0.00608 |
| Molformer | 0.809 ± 0.00857 | 0.948 ± 0.0227 | 0.926 ± 0.00403 | 0.7 ± 0.014 | 0.974 ± 0.0081 |
| GROVER-Base | 0.833 ± 0.0132 | 0.932 ± 0.0355 | 0.9 ± 0.00965 | 0.646 ± 0.0138 | 1.05 ± 0.0386 |
| GROVER-Large | 0.975 ± 0.131 | 1.11 ± 0.175 | 0.968 ± 0.0698 | 1.11 ± 0.635 | 1.13 ± 0.0741 |
| MoleBERT | 0.901 ± 0.00358 | 0.989 ± 0.00418 | 0.89 ± 0.00417 | 0.671 ± 0.00625 | 1.31 ± 0.00618 |
| UniMol-G | 0.952 ± 0.0154 | 0.999 ± 0.00643 | 0.864 ± 0.0111 | 0.805 ± 0.0246 | 1.37 ± 0.00797 |
| UniMol-GA | 0.942 ± 0.0111 | 1.03 ± 0.00793 | 0.842 ± 0.0123 | 0.796 ± 0.0157 | 1.31 ± 0.00659 |
| GCC | 0.907 ± 0.0132 | 1.06 ± 0.0178 | 0.92 ± 0.0102 | 0.781 ± 0.00255 | 1.36 ± 0.00683 |
| GPT-GNN | 1.11 ± 0.0275 | 1.13 ± 0.0036 | 0.965 ± 0.00241 | 0.801 ± 0.00591 | 1.54 ± 0.0127 |
| GraphCL | 1.11 ± 0.0263 | 1.04 ± 0.00643 | 0.972 ± 0.00413 | 0.955 ± 0.00572 | 1.57 ± 0.00808 |
| InfoGraph | 1.35 ± 0.0305 | 1.17 ± 0.0103 | 0.903 ± 0.00191 | 0.886 ± 0.00631 | 1.57 ± 0.0234 |
| JOAO | 1.13 ± 0.0208 | 1.16 ± 0.0127 | 0.926 ± 0.0015 | 0.834 ± 0.0106 | 1.55 ± 0.0152 |
| JOAOv2 | 1.1 ± 0.0224 | 1.22 ± 0.00635 | 0.944 ± 0.00254 | 0.857 ± 0.0152 | 1.56 ± 0.0109 |
| PosPred | 1.03 ± 0.043 | 1.18 ± 0.00607 | 0.911 ± 0.00266 | 0.841 ± 0.0104 | 1.47 ± 0.0103 |
| 3D-EMGP | 1.21 ± 0.0209 | 1.13 ± 0.0163 | 1.02 ± 0.0108 | 0.833 ± 0.0101 | 1.55 ± 0.0141 |
| 3D-InfoMax | 1.21 ± 0.0209 | 1.13 ± 0.0163 | 1.02 ± 0.0108 | 0.833 ± 0.0101 | 1.55 ± 0.0141 |
| AttrMask | 1.31 ± 0.0141 | 1.19 ± 0.00494 | 0.935 ± 0.00152 | 0.864 ± 0.00674 | 1.57 ± 0.0203 |
| EdgePred | 1.37 ± 0.0162 | 1.18 ± 0.00637 | 0.95 ± 0.00192 | 0.881 ± 0.00523 | 1.69 ± 0.0101 |
| GraphMVP | 1.45 ± 0.0153 | 1.18 ± 0.000215 | 0.938 ± 0.00022 | 0.873 ± 0.000582 | 1.68 ± 0.0106 |
| MolCLR-GIN | 1.42 ± 0.00609 | 1.17 ± 0.0116 | 0.931 ± 0.00366 | 0.888 ± 0.00582 | 1.66 ± 0.00593 |
| MolCLR-GCN | 1.44 ± 0.00331 | 1.18 ± 0.0112 | 0.933 ± 0.0024 | 0.88 ± 0.0034 | 1.67 ± 0.00494 |

Prediction is evaluated by RMSE. For each method, the mean and standard deviation are reported for the evaluation metrics.

**Supplementary Table 28.** Activity cliff prediction (cliff molecules) – Part II.

| Method | CHEMBL2147<br>$K_i$ | CHEMBL214<br>$K_i$ | CHEMBL218<br>$EC_{50}$ | CHEMBL219<br>$K_i$ | CHEMBL228<br>$K_i$ |
| --- | --- | --- | --- | --- | --- |
| qcGEM | 0.667 ± 0.00263 | 0.893 ± 0.00218 | 0.823 ± 0.00459 | 0.811 ± 0.00572 | 0.771 ± 0.00907 |
| ECFP | 0.771 ± 0.0135 | 0.788 ± 0.00182 | 0.875 ± 0.00342 | 0.893 ± 0.00508 | 0.872 ± 0.00404 |
| FCFP | 0.693 ± 0.00913 | 0.842 ± 0.0045 | 0.789 ± 0.00281 | 0.898 ± 0.00691 | 0.888 ± 0.00543 |
| AvalonFP | 0.783 ± 0.00477 | 0.796 ± 0.00335 | 0.806 ± 0.0101 | 0.867 ± 0.00393 | 0.836 ± 0.00389 |
| TTGEN | 0.804 ± 0.00726 | 0.874 ± 0.00265 | 0.814 ± 0.00876 | 0.857 ± 0.00689 | 0.92 ± 0.00399 |
| APGEN | 0.786 ± 0.0047 | 0.934 ± 0.00681 | 0.858 ± 0.00888 | 0.884 ± 0.00514 | 0.926 ± 0.00417 |
| RDKFP | 0.732 ± 0.00611 | 0.897 ± 0.00248 | 0.757 ± 0.0133 | 0.902 ± 0.00437 | 0.898 ± 0.00495 |
| MACCS | 0.927 ± 0.00787 | 0.963 ± 0.0027 | 0.865 ± 0.00739 | 0.944 ± 0.00118 | 0.992 ± 0.006 |
| LayeredFP | 0.801 ± 0.00568 | 0.957 ± 0.00666 | 0.883 ± 0.00322 | 0.92 ± 0.00234 | 0.889 ± 0.00322 |
| PatternFP | 0.851 ± 0.00448 | 0.935 ± 0.00559 | 0.879 ± 0.00378 | 0.887 ± 0.00175 | 0.978 ± 0.00278 |
| Molformer | 0.791 ± 0.00522 | 0.852 ± 0.00245 | 0.8 ± 0.00916 | 0.853 ± 0.00632 | 0.838 ± 0.00808 |
| GROVER-Base | 0.781 ± 0.0109 | 0.886 ± 0.0132 | 0.845 ± 0.0334 | 0.865 ± 0.0241 | 0.858 ± 0.0159 |
| GROVER-Large | 0.856 ± 0.0236 | 0.975 ± 0.0595 | 0.991 ± 0.203 | 0.949 ± 0.0451 | 0.938 ± 0.0251 |
| MoleBERT | 0.886 ± 0.00477 | 1.01 ± 0.00189 | 0.924 ± 0.00851 | 0.95 ± 0.00228 | 1 ± 0.0041 |
| UniMol-G | 0.882 ± 0.0061 | 1.02 ± 0.0012 | 0.932 ± 0.00477 | 1.04 ± 0.00187 | 1.05 ± 0.0106 |
| UniMol-GA | 0.865 ± 0.0167 | 1.01 ± 0.0028 | 0.938 ± 0.00656 | 1.03 ± 0.00115 | 1.02 ± 0.00518 |
| GCC | 1.03 ± 0.0121 | 1.07 ± 0.00617 | 0.927 ± 0.0132 | 1.02 ± 0.00399 | 1.07 ± 0.00896 |
| GPT-GNN | 1.06 ± 0.0126 | 1.13 ± 0.00247 | 1.01 ± 0.00667 | 1 ± 0.00277 | 1.11 ± 0.00668 |
| GraphCL | 1.06 ± 0.0048 | 1.12 ± 0.00271 | 0.973 ± 0.00633 | 1.03 ± 0.00227 | 1.09 ± 0.00621 |
| InfoGraph | 1.32 ± 0.0253 | 1.15 ± 0.00436 | 0.982 ± 0.00914 | 1.05 ± 0.000807 | 1.14 ± 0.00736 |
| JOAO | 1.09 ± 0.0111 | 1.13 ± 0.00211 | 0.965 ± 0.00276 | 1.01 ± 0.00142 | 1.12 ± 0.00363 |
| JOAOv2 | 1.1 ± 0.014 | 1.14 ± 0.00106 | 0.977 ± 0.00162 | 0.998 ± 0.000781 | 1.1 ± 0.00354 |
| PosPred | 1.13 ± 0.0143 | 1.12 ± 0.0018 | 0.982 ± 0.00341 | 0.977 ± 0.00183 | 1.11 ± 0.000774 |
| 3D-EMGP | 1.33 ± 0.0129 | 1.13 ± 0.00415 | 1.03 ± 0.0071 | 1.08 ± 0.00549 | 1.18 ± 0.00991 |
| 3D-InfoMax | 1.33 ± 0.0129 | 1.13 ± 0.00415 | 1.03 ± 0.0071 | 1.08 ± 0.00549 | 1.18 ± 0.00991 |
| AttrMask | 1.2 ± 0.009 | 1.12 ± 0.00279 | 1.02 ± 0.00162 | 1.04 ± 0.00162 | 1.14 ± 0.00137 |
| EdgePred | 1.54 ± 0.0166 | 1.14 ± 0.00109 | 1 ± 0.00135 | 1.05 ± 0.00121 | 1.12 ± 0.00384 |
| GraphMVP | 1.72 ± 0.0364 | 1.15 ± 0.000121 | 1.03 ± 0.000743 | 1.07 ± 0.000251 | 1.14 ± 0.000143 |
| MolCLR-GIN | 1.63 ± 0.0306 | 1.15 ± 0.00215 | 1.03 ± 0.00666 | 1.06 ± 0.00271 | 1.14 ± 0.0032 |
| MolCLR-GCN | 1.59 ± 0.0115 | 1.15 ± 0.00188 | 1.04 ± 0.00953 | 1.07 ± 0.00182 | 1.15 ± 0.00899 |

Prediction is evaluated by RMSE. For each method, the mean and standard deviation are reported for the evaluation metrics.

**Supplementary Table 29.** Activity cliff prediction (cliff molecules) – Part III.

| Method | CHEMBL231<br>$K_i$ | CHEMBL233<br>$K_i$ | CHEMBL234<br>$K_i$ | CHEMBL235<br>$EC_{50}$ | CHEMBL236<br>$K_i$ |
| --- | --- | --- | --- | --- | --- |
| qcGEM | 0.794 ± 0.0129 | 0.928 ± 0.00537 | 0.714 ± 0.00451 | 0.823 ± 0.0129 | 0.847 ± 0.0073 |
| ECFP | 0.983 ± 0.0281 | 0.955 ± 0.00414 | 0.703 ± 0.00696 | 0.863 ± 0.00738 | 0.813 ± 0.00454 |
| FCFP | 0.884 ± 0.00416 | 1.01 ± 0.00716 | 0.73 ± 0.00481 | 0.844 ± 0.0066 | 0.858 ± 0.00557 |
| AvalonFP | 0.997 ± 0.01 | 0.971 ± 0.0038 | 0.731 ± 0.00699 | 0.853 ± 0.00275 | 0.878 ± 0.00257 |
| TTGEN | 0.964 ± 0.0107 | 1.03 ± 0.00617 | 0.736 ± 0.00473 | 0.934 ± 0.00352 | 0.88 ± 0.00351 |
| APGEN | 0.79 ± 0.013 | 1.03 ± 0.00543 | 0.793 ± 0.00825 | 0.869 ± 0.00857 | 0.952 ± 0.00597 |
| RDKFP | 0.825 ± 0.00363 | 1.05 ± 0.00295 | 0.788 ± 0.00322 | 0.925 ± 0.00365 | 0.985 ± 0.00275 |
| MACCS | 0.895 ± 0.0083 | 1.1 ± 0.00303 | 0.822 ± 0.00661 | 1.01 ± 0.00372 | 1.06 ± 0.0107 |
| LayeredFP | 0.886 ± 0.00685 | 1.12 ± 0.00457 | 0.844 ± 0.00258 | 0.971 ± 0.0029 | 1.02 ± 0.00501 |
| PatternFP | 0.848 ± 0.00976 | 1.08 ± 0.00453 | 0.837 ± 0.00281 | 0.951 ± 0.00763 | 1.03 ± 0.00582 |
| Molformer | 0.944 ± 0.013 | 0.972 ± 0.00553 | 0.737 ± 0.00514 | 0.75 ± 0.0039 | 0.859 ± 0.00578 |
| GROVER-Base | 0.921 ± 0.057 | 0.981 ± 0.0158 | 0.753 ± 0.00809 | 0.87 ± 0.0107 | 0.884 ± 0.00957 |
| GROVER-Large | 1.1 ± 0.295 | 0.991 ± 0.0151 | 0.772 ± 0.00758 | 0.928 ± 0.0552 | 0.947 ± 0.087 |
| MoleBERT | 0.912 ± 0.00401 | 1.13 ± 0.00182 | 0.879 ± 0.00195 | 1.06 ± 0.00949 | 1.07 ± 0.00491 |
| UniMol-G | 0.935 ± 0.00971 | 1.11 ± 0.00615 | 0.902 ± 0.00237 | 1.07 ± 0.00979 | 1.12 ± 0.00587 |
| UniMol-GA | 0.885 ± 0.0126 | 1.08 ± 0.00386 | 0.885 ± 0.00145 | 1.04 ± 0.00792 | 1.11 ± 0.00752 |
| GCC | 0.986 ± 0.0206 | 1.13 ± 0.00659 | 0.936 ± 0.00231 | 1.05 ± 0.00551 | 1.11 ± 0.00509 |
| GPT-GNN | 1.1 ± 0.0036 | 1.27 ± 0.00397 | 0.94 ± 0.000729 | 1.17 ± 0.00735 | 1.31 ± 0.00906 |
| GraphCL | 1.09 ± 0.00402 | 1.3 ± 0.00285 | 0.927 ± 0.00208 | 1.14 ± 0.0101 | 1.3 ± 0.00733 |
| InfoGraph | 1.11 ± 0.00636 | 1.28 ± 0.00405 | 1 ± 0.00164 | 1.15 ± 0.00958 | 1.34 ± 0.0063 |
| JOAO | 1.1 ± 0.000805 | 1.31 ± 0.00338 | 0.931 ± 0.00078 | 1.14 ± 0.00168 | 1.31 ± 0.00548 |
| JOAOv2 | 1.09 ± 0.0036 | 1.29 ± 0.00615 | 0.905 ± 0.00121 | 1.14 ± 0.00488 | 1.28 ± 0.00875 |
| PosPred | 1.04 ± 0.00705 | 1.19 ± 0.00473 | 0.902 ± 0.00144 | 1.13 ± 0.00639 | 1.28 ± 0.0116 |
| 3D-EMGP | 1.06 ± 0.0122 | 1.29 ± 0.0046 | 0.955 ± 0.00241 | 1.15 ± 0.00724 | 1.32 ± 0.00304 |
| 3D-InfoMax | 1.06 ± 0.0122 | 1.29 ± 0.0046 | 0.955 ± 0.00241 | 1.15 ± 0.00724 | 1.32 ± 0.00304 |
| AttrMask | 1.13 ± 0.00754 | 1.31 ± 0.00158 | 0.991 ± 0.00313 | 1.16 ± 0.00598 | 1.31 ± 0.00397 |
| EdgePred | 1.17 ± 0.000804 | 1.33 ± 0.00137 | 1.02 ± 0.000998 | 1.16 ± 0.00424 | 1.32 ± 0.0044 |
| GraphMVP | 1.23 ± 3.32e-05 | 1.33 ± 0.00021 | 1.04 ± 0.000219 | 1.16 ± 0.000894 | 1.32 ± 0.000314 |
| MolCLR-GIN | 1.2 ± 0.00925 | 1.33 ± 0.00127 | 1.05 ± 0.00222 | 1.16 ± 0.00491 | 1.32 ± 0.00379 |
| MolCLR-GCN | 1.24 ± 0.00102 | 1.33 ± 0.00105 | 1.06 ± 0.00234 | 1.16 ± 0.0048 | 1.32 ± 0.00507 |

Prediction is evaluated by RMSE. For each method, the mean and standard deviation are reported for the evaluation metrics.

**Supplementary Table 30.** Activity cliff prediciton (cliff molecules) – Part IV.

| Method | CHEMBL237<br>EC<sub>50</sub> | CHEMBL237<br>K<sub>i</sub> | CHEMBL238<br>K<sub>i</sub> | CHEMBL239<br>EC<sub>50</sub> | CHEMBL244<br>K<sub>i</sub> |
| --- | --- | --- | --- | --- | --- |
| qcGEM | 0.913 ± 0.00897 | 0.786 ± 0.00411 | 0.792 ± 0.0107 | 0.863 ± 0.00662 | 0.917 ± 0.00366 |
| ECFP | 0.864 ± 0.0077 | 0.813 ± 0.00593 | 0.863 ± 0.0115 | 0.97 ± 0.00283 | 0.934 ± 0.00513 |
| FCFP | 0.991 ± 0.00795 | 0.811 ± 0.00439 | 0.818 ± 0.00727 | 0.897 ± 0.00474 | 0.933 ± 0.00119 |
| AvalonFP | 0.925 ± 0.00558 | 0.901 ± 0.00294 | 0.757 ± 0.00538 | 0.888 ± 0.00919 | 0.922 ± 0.007 |
| TTGEN | 1.07 ± 0.011 | 0.88 ± 0.00115 | 0.77 ± 0.00287 | 1.06 ± 0.00553 | 1.01 ± 0.00693 |
| APGEN | 1.01 ± 0.00949 | 0.9 ± 0.00455 | 0.841 ± 0.0136 | 1.03 ± 0.00437 | 1.05 ± 0.00712 |
| RDKFP | 0.995 ± 0.00549 | 1.02 ± 0.00541 | 0.824 ± 0.0138 | 1.01 ± 0.00347 | 0.943 ± 0.00334 |
| MACCS | 0.979 ± 0.00558 | 1 ± 0.00563 | 0.822 ± 0.00544 | 1.02 ± 0.0081 | 1.02 ± 0.00749 |
| LayeredFP | 1.13 ± 0.00314 | 1 ± 0.00175 | 0.818 ± 0.0116 | 0.951 ± 0.00477 | 1.05 ± 0.00588 |
| PatternFP | 1.15 ± 0.00263 | 1.04 ± 0.00423 | 0.889 ± 0.00866 | 1 ± 0.00399 | 1.05 ± 0.0051 |
| Molformer | 0.888 ± 0.00576 | 0.845 ± 0.00231 | 0.753 ± 0.006 | 0.938 ± 0.00807 | 0.891 ± 0.00408 |
| GROVER-Base | 0.954 ± 0.0358 | 0.87 ± 0.0175 | 0.836 ± 0.0148 | 0.96 ± 0.0194 | 0.957 ± 0.016 |
| GROVER-Large | 1.16 ± 0.15 | 0.959 ± 0.125 | 0.988 ± 0.203 | 0.991 ± 0.0263 | 0.97 ± 0.029 |
| MoleBERT | 1.11 ± 0.00268 | 1.06 ± 0.00297 | 0.931 ± 0.00609 | 1.02 ± 0.00623 | 1.16 ± 0.00169 |
| UniMol-G | 1.29 ± 0.00237 | 1.08 ± 0.00975 | 1.02 ± 0.00915 | 1.04 ± 0.00331 | 1.12 ± 0.00225 |
| UniMol-GA | 1.26 ± 0.00235 | 1.03 ± 0.00598 | 0.987 ± 0.0195 | 1.02 ± 0.00849 | 1.09 ± 0.00194 |
| GCC | 1.19 ± 0.00831 | 1.03 ± 0.00887 | 0.914 ± 0.0174 | 1.05 ± 0.00852 | 1.18 ± 0.005 |
| GPT-GNN | 1.33 ± 0.00166 | 1.28 ± 0.00211 | 1.1 ± 0.0175 | 1.13 ± 0.00433 | 1.31 ± 0.00352 |
| GraphCL | 1.31 ± 0.00105 | 1.32 ± 0.00492 | 1.08 ± 0.0108 | 1.14 ± 0.00784 | 1.37 ± 0.00409 |
| InfoGraph | 1.31 ± 0.00331 | 1.28 ± 0.00439 | 1.17 ± 0.0124 | 1.14 ± 0.00479 | 1.38 ± 0.00208 |
| JOAO | 1.34 ± 0.0011 | 1.31 ± 0.00441 | 1.11 ± 0.0129 | 1.12 ± 0.0051 | 1.37 ± 0.00271 |
| JOAOv2 | 1.32 ± 0.00119 | 1.37 ± 0.00401 | 1.08 ± 0.0234 | 1.12 ± 0.00374 | 1.26 ± 0.00465 |
| PosPred | 1.27 ± 0.00652 | 1.19 ± 0.00654 | 1.07 ± 0.0174 | 1.1 ± 0.00301 | 1.26 ± 0.00999 |
| 3D-EMGP | 1.35 ± 0.00925 | 1.35 ± 0.00374 | 1.21 ± 0.0417 | 1.17 ± 0.0156 | 1.41 ± 0.00612 |
| 3D-InfoMax | 1.35 ± 0.00925 | 1.35 ± 0.00374 | 1.21 ± 0.0417 | 1.17 ± 0.0156 | 1.41 ± 0.00612 |
| AttrMask | 1.31 ± 0.0036 | 1.27 ± 0.00407 | 1.18 ± 0.0068 | 1.16 ± 0.00771 | 1.37 ± 0.00469 |
| EdgePred | 1.37 ± 0.00113 | 1.42 ± 0.00648 | 1.16 ± 0.0104 | 1.17 ± 0.0038 | 1.54 ± 0.000636 |
| GraphMVP | 1.36 ± 0.00057 | 1.43 ± 0.000213 | 1.2 ± 0.000419 | 1.18 ± 0.000535 | 1.55 ± 0.000118 |
| MolCLR-GIN | 1.35 ± 0.000944 | 1.44 ± 0.00484 | 1.22 ± 0.0152 | 1.18 ± 0.0226 | 1.55 ± 0.00232 |
| MolCLR-GCN | 1.36 ± 0.00153 | 1.44 ± 0.00267 | 1.21 ± 0.0106 | 1.18 ± 0.0117 | 1.56 ± 0.000713 |

Prediction is evaluated by RMSE. For each method, the mean and standard deviation are reported for the evaluation metrics.

**Supplementary Table 31.** Activity cliff prediction (cliff molecules) – Part V.

| Method | CHEMBL262<br>$K_i$ | CHEMBL264<br>$K_i$ | CHEMBL2835<br>$K_i$ | CHEMBL287<br>$K_i$ | CHEMBL2971<br>$K_i$ |
| --- | --- | --- | --- | --- | --- |
| qcGEM | 0.93 ± 0.0129 | 0.726 ± 0.00293 | 0.72 ± 0.0261 | 0.733 ± 0.0275 | 0.725 ± 0.0214 |
| ECFP | 0.961 ± 0.0197 | 0.81 ± 0.00409 | 0.831 ± 0.012 | 0.957 ± 0.0105 | 0.805 ± 0.0207 |
| FCFP | 0.864 ± 0.00738 | 0.753 ± 0.00236 | 0.771 ± 0.00596 | 0.921 ± 0.00841 | 0.873 ± 0.019 |
| AvalonFP | 0.89 ± 0.00568 | 0.771 ± 0.00301 | 0.85 ± 0.0201 | 0.898 ± 0.0124 | 0.878 ± 0.00687 |
| TTGEN | 0.952 ± 0.00708 | 0.844 ± 0.00393 | 0.828 ± 0.00915 | 0.903 ± 0.0101 | 0.815 ± 0.0104 |
| APGEN | 0.871 ± 0.0075 | 0.83 ± 0.00807 | 0.963 ± 0.0188 | 0.927 ± 0.00922 | 0.872 ± 0.0165 |
| RDKFP | 0.992 ± 0.00439 | 0.884 ± 0.00443 | 0.781 ± 0.0243 | 0.985 ± 0.00223 | 0.772 ± 0.0104 |
| MACCS | 0.993 ± 0.00717 | 0.911 ± 0.00901 | 0.797 ± 0.0111 | 0.932 ± 0.0135 | 0.678 ± 0.00748 |
| LayeredFP | 0.875 ± 0.0026 | 0.831 ± 0.0058 | 0.838 ± 0.0108 | 0.988 ± 0.00474 | 0.84 ± 0.00319 |
| PatternFP | 0.963 ± 0.00734 | 0.861 ± 0.00442 | 0.72 ± 0.0109 | 0.896 ± 0.00589 | 0.731 ± 0.00477 |
| Molformer | 0.948 ± 0.0123 | 0.751 ± 0.00447 | 0.744 ± 0.00859 | 0.854 ± 0.00635 | 0.811 ± 0.0146 |
| GROVER-Base | 0.942 ± 0.0267 | 0.785 ± 0.0146 | 0.762 ± 0.0135 | 0.805 ± 0.011 | 0.729 ± 0.0188 |
| GROVER-Large | 1.18 ± 0.392 | 0.787 ± 0.0194 | 0.902 ± 0.183 | 0.913 ± 0.114 | 0.903 ± 0.201 |
| MoleBERT | 0.922 ± 0.0031 | 0.919 ± 0.00131 | 0.791 ± 0.0046 | 0.913 ± 0.005 | 0.834 ± 0.00519 |
| UniMol-G | 0.972 ± 0.0038 | 0.969 ± 0.0032 | 0.906 ± 0.0132 | 0.916 ± 0.00826 | 0.86 ± 0.00476 |
| UniMol-GA | 0.954 ± 0.00663 | 0.933 ± 0.00526 | 0.944 ± 0.0063 | 0.867 ± 0.00787 | 0.851 ± 0.00709 |
| GCC | 0.937 ± 0.0213 | 0.953 ± 0.00675 | 1.08 ± 0.0157 | 0.955 ± 0.00718 | 0.95 ± 0.0152 |
| GPT-GNN | 1.09 ± 0.0061 | 1.03 ± 0.00526 | 0.947 ± 0.0103 | 0.978 ± 0.0103 | 0.918 ± 0.00799 |
| GraphCL | 1.05 ± 0.00421 | 1.04 ± 0.00525 | 0.814 ± 0.00774 | 0.96 ± 0.00553 | 0.875 ± 0.00627 |
| InfoGraph | 1.05 ± 0.00646 | 1.03 ± 0.00275 | 1.01 ± 0.014 | 1.04 ± 0.00742 | 0.876 ± 0.00646 |
| JOAO | 1.07 ± 0.00412 | 0.991 ± 0.00291 | 0.731 ± 0.00916 | 1.08 ± 0.00277 | 0.892 ± 0.0109 |
| JOAOv2 | 1.06 ± 0.00423 | 1.01 ± 0.00413 | 0.86 ± 0.012 | 0.996 ± 0.00394 | 0.874 ± 0.00368 |
| PosPred | 1.03 ± 0.00152 | 1.04 ± 0.00168 | 0.891 ± 0.00609 | 1.03 ± 0.00787 | 0.903 ± 0.00339 |
| 3D-EMGP | 1.05 ± 0.0169 | 1.02 ± 0.00589 | 0.876 ± 0.0177 | 1.05 ± 0.00972 | 1.04 ± 0.0148 |
| 3D-InfoMax | 1.05 ± 0.0169 | 1.02 ± 0.00589 | 0.876 ± 0.0177 | 1.05 ± 0.00972 | 1.04 ± 0.0148 |
| AttrMask | 1.16 ± 0.00455 | 1.02 ± 0.00425 | 1.2 ± 0.214 | 1.11 ± 0.00619 | 1.08 ± 0.00468 |
| EdgePred | 1.11 ± 0.00317 | 1.02 ± 0.00805 | 1.1 ± 0.00567 | 1.11 ± 0.00406 | 1.15 ± 0.00491 |
| GraphMVP | 1.16 ± 0.00276 | 1.02 ± 0.00255 | 1.52 ± 0.014 | 1.15 ± 0.00318 | 1.33 ± 0.0208 |
| MolCLR-GIN | 1.13 ± 0.0181 | 1.03 ± 0.00154 | 1.54 ± 0.00353 | 1.15 ± 0.00669 | 1.34 ± 0.0022 |
| MolCLR-GCN | 1.22 ± 0.0126 | 1.03 ± 0.00112 | 1.54 ± 0.00557 | 1.14 ± 0.00332 | 1.36 ± 0.0228 |

Prediction is evaluated by RMSE. For each method, the mean and standard deviation are reported for the evaluation metrics.

**Supplementary Table 32.**
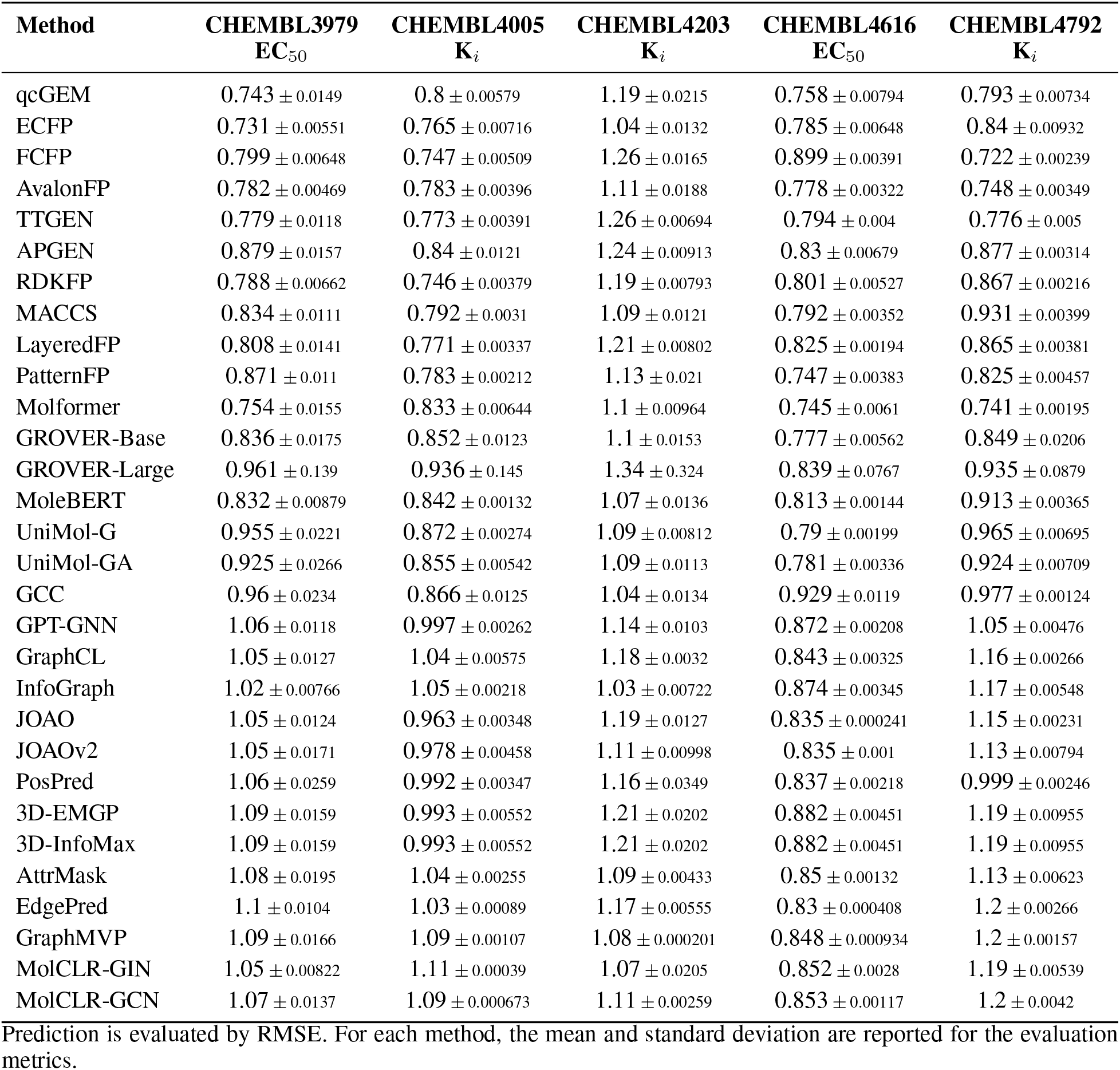
Activity cliff prediction (cliff molecules) – Part VI.

| Method | CHEMBL3979<br>EC<sub>50</sub> | CHEMBL4005<br>K<sub>i</sub> | CHEMBL4203<br>K<sub>i</sub> | CHEMBL4616<br>EC<sub>50</sub> | CHEMBL4792<br>K<sub>i</sub> |
| --- | --- | --- | --- | --- | --- |
| qcGEM | 0.743 ± 0.0149 | 0.8 ± 0.00579 | 1.19 ± 0.0215 | 0.758 ± 0.00794 | 0.793 ± 0.00734 |
| ECFP | 0.731 ± 0.00551 | 0.765 ± 0.00716 | 1.04 ± 0.0132 | 0.785 ± 0.00648 | 0.84 ± 0.00932 |
| FCFP | 0.799 ± 0.00648 | 0.747 ± 0.00509 | 1.26 ± 0.0165 | 0.899 ± 0.00391 | 0.722 ± 0.00239 |
| AvalonFP | 0.782 ± 0.00469 | 0.783 ± 0.00396 | 1.11 ± 0.0188 | 0.778 ± 0.00322 | 0.748 ± 0.00349 |
| TTGEN | 0.779 ± 0.0118 | 0.773 ± 0.00391 | 1.26 ± 0.00694 | 0.794 ± 0.004 | 0.776 ± 0.005 |
| APGEN | 0.879 ± 0.0157 | 0.84 ± 0.0121 | 1.24 ± 0.00913 | 0.83 ± 0.00679 | 0.877 ± 0.00314 |
| RDKFP | 0.788 ± 0.00662 | 0.746 ± 0.00379 | 1.19 ± 0.00793 | 0.801 ± 0.00527 | 0.867 ± 0.00216 |
| MACCS | 0.834 ± 0.0111 | 0.792 ± 0.0031 | 1.09 ± 0.0121 | 0.792 ± 0.00352 | 0.931 ± 0.00399 |
| LayeredFP | 0.808 ± 0.0141 | 0.771 ± 0.00337 | 1.21 ± 0.00802 | 0.825 ± 0.00194 | 0.865 ± 0.00381 |
| PatternFP | 0.871 ± 0.011 | 0.783 ± 0.00212 | 1.13 ± 0.021 | 0.747 ± 0.00383 | 0.825 ± 0.00457 |
| Molformer | 0.754 ± 0.0155 | 0.833 ± 0.00644 | 1.1 ± 0.00964 | 0.745 ± 0.0061 | 0.741 ± 0.00195 |
| GROVER-Base | 0.836 ± 0.0175 | 0.852 ± 0.0123 | 1.1 ± 0.0153 | 0.777 ± 0.00562 | 0.849 ± 0.0206 |
| GROVER-Large | 0.961 ± 0.139 | 0.936 ± 0.145 | 1.34 ± 0.324 | 0.839 ± 0.0767 | 0.935 ± 0.0879 |
| MoleBERT | 0.832 ± 0.00879 | 0.842 ± 0.00132 | 1.07 ± 0.0136 | 0.813 ± 0.00144 | 0.913 ± 0.00365 |
| UniMol-G | 0.955 ± 0.0221 | 0.872 ± 0.00274 | 1.09 ± 0.00812 | 0.79 ± 0.00199 | 0.965 ± 0.00695 |
| UniMol-GA | 0.925 ± 0.0266 | 0.855 ± 0.00542 | 1.09 ± 0.0113 | 0.781 ± 0.00336 | 0.924 ± 0.00709 |
| GCC | 0.96 ± 0.0234 | 0.866 ± 0.0125 | 1.04 ± 0.0134 | 0.929 ± 0.0119 | 0.977 ± 0.00124 |
| GPT-GNN | 1.06 ± 0.0118 | 0.997 ± 0.00262 | 1.14 ± 0.0103 | 0.872 ± 0.00208 | 1.05 ± 0.00476 |
| GraphCL | 1.05 ± 0.0127 | 1.04 ± 0.00575 | 1.18 ± 0.0032 | 0.843 ± 0.00325 | 1.16 ± 0.00266 |
| InfoGraph | 1.02 ± 0.00766 | 1.05 ± 0.00218 | 1.03 ± 0.00722 | 0.874 ± 0.00345 | 1.17 ± 0.00548 |
| JOAO | 1.05 ± 0.0124 | 0.963 ± 0.00348 | 1.19 ± 0.0127 | 0.835 ± 0.000241 | 1.15 ± 0.00231 |
| JOAOv2 | 1.05 ± 0.0171 | 0.978 ± 0.00458 | 1.11 ± 0.00998 | 0.835 ± 0.001 | 1.13 ± 0.00794 |
| PosPred | 1.06 ± 0.0259 | 0.992 ± 0.00347 | 1.16 ± 0.0349 | 0.837 ± 0.00218 | 0.999 ± 0.00246 |
| 3D-EMGP | 1.09 ± 0.0159 | 0.993 ± 0.00552 | 1.21 ± 0.0202 | 0.882 ± 0.00451 | 1.19 ± 0.00955 |
| 3D-InfoMax | 1.09 ± 0.0159 | 0.993 ± 0.00552 | 1.21 ± 0.0202 | 0.882 ± 0.00451 | 1.19 ± 0.00955 |
| AttrMask | 1.08 ± 0.0195 | 1.04 ± 0.00255 | 1.09 ± 0.00433 | 0.85 ± 0.00132 | 1.13 ± 0.00623 |
| EdgePred | 1.1 ± 0.0104 | 1.03 ± 0.00089 | 1.17 ± 0.00555 | 0.83 ± 0.000408 | 1.2 ± 0.00266 |
| GraphMVP | 1.09 ± 0.0166 | 1.09 ± 0.00107 | 1.08 ± 0.000201 | 0.848 ± 0.000934 | 1.2 ± 0.00157 |
| MolCLR-GIN | 1.05 ± 0.00822 | 1.11 ± 0.00039 | 1.07 ± 0.0205 | 0.852 ± 0.0028 | 1.19 ± 0.00539 |
| MolCLR-GCN | 1.07 ± 0.0137 | 1.09 ± 0.000673 | 1.11 ± 0.00259 | 0.853 ± 0.00117 | 1.2 ± 0.0042 |

Prediction is evaluated by RMSE. For each method, the mean and standard deviation are reported for the evaluation metrics.

##### 3.3.3 Protein-ligand interaction prediction

**Supplementary Figure 14.**
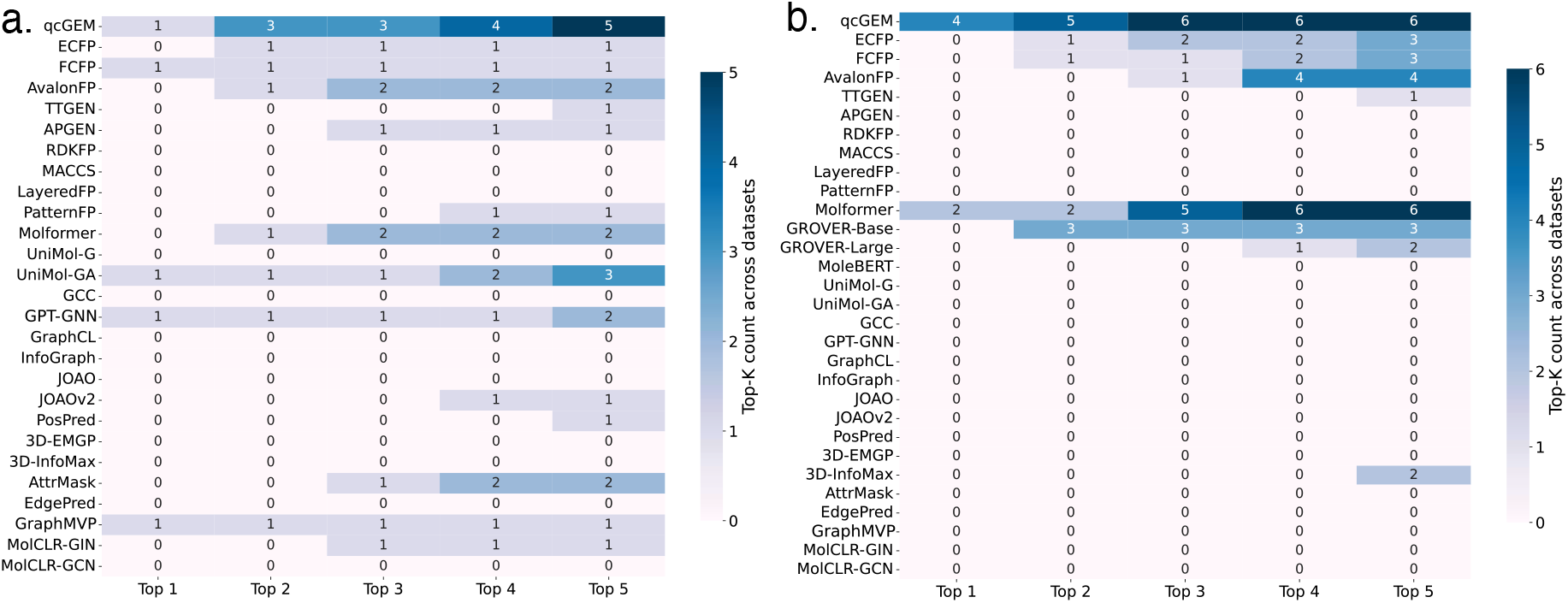
Statistics of the top 1-5 rankings for all methods in protein-ligand interaction tasks. **a)** Evaluation results of qcGEM and baseline methods on 5 protein-ligand interaction tasks. **b)** Evaluation results of qcGEM and baseline methods on 6 opioids-related tasks.

**Supplementary Figure 15.**
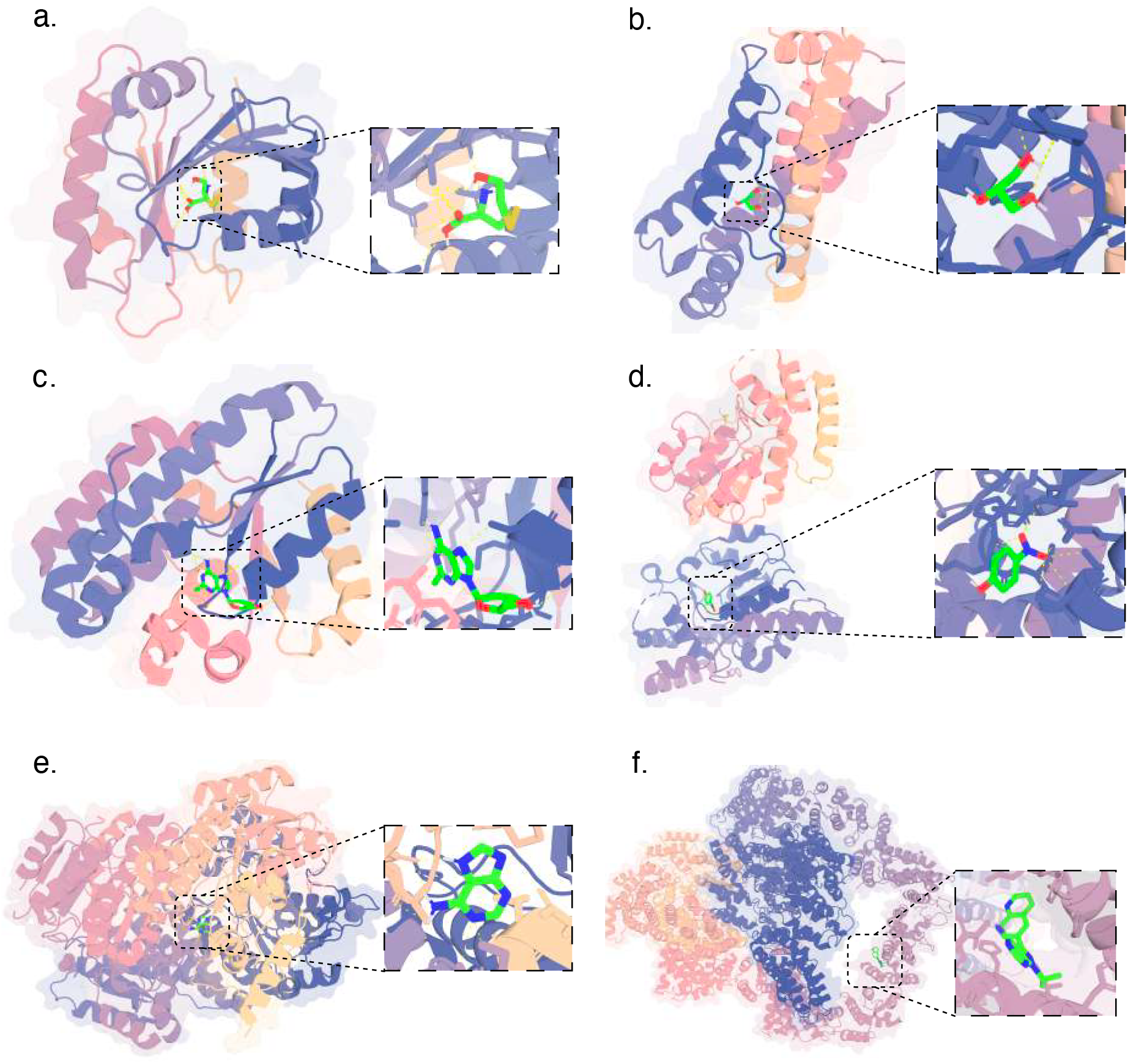
Additional docking results. This figure supplements Figure 4e, showing more examples for the atomic level interaction between the predicted positive ligand and target protein. The presented proteins include **a)** 4ILE; **b)** 1SWX; **c)** 1P5Z; **d)** 1ZD1; **e)** 1YXM; **f)** 5W1R.

**Supplementary Table 33.** Protein-ligand interaction.

| Method | Regression |  | Classification |  |  |
| --- | --- | --- | --- | --- | --- |
|  | KIBA | DAVIS | Human | C.elegans | BindingDB |
| qcGEM | 0.805 ± 0.00414 | 0.817 ± 0.000295 | 0.945 ± 0.00143 | 0.883 ± 0.12 | 0.893 ± 0.000561 |
| ECFP | 0.804 ± 0.004 | 0.821 ± 0.000234 | 0.526 ± 0.0577 | 0.521 ± 0.0463 | 0.422 ± 0.0411 |
| FCFP | 0.802 ± 0.00445 | 0.821 ± 0.000142 | 0.533 ± 0.0741 | 0.54 ± 0.0757 | 0.432 ± 0.0475 |
| AvalonFP | 0.805 ± 0.00169 | 0.82 ± 0.000351 | 0.534 ± 0.076 | 0.537 ± 0.0506 | 0.892 ± 0.00156 |
| TTGEN | 0.806 ± 0.00295 | 0.82 ± 0.000266 | 0.54 ± 0.0895 | 0.54 ± 0.0592 | 0.42 ± 0.0919 |
| APGEN | 0.809 ± 0.00272 | 0.819 ± 0.000359 | 0.541 ± 0.0926 | 0.585 ± 0.078 | 0.864 ± 0.00127 |
| RDKFP | 0.808 ± 0.00203 | 0.822 ± 0.000639 | 0.55 ± 0.112 | 0.562 ± 0.0845 | 0.521 ± 0.198 |
| MACCS | 0.814 ± 0.00218 | 0.821 ± 0.000148 | 0.5 ± 0 | 0.544 ± 0.0616 | 0.354 ± 0.0416 |
| LayeredFP | 0.81 ± 0.00117 | 0.818 ± 0.000611 | 0.547 ± 0.106 | 0.581 ± 0.0804 | 0.796 ± 0.000423 |
| PatternFP | 0.807 ± 0.00123 | 0.817 ± 0.000252 | 0.526 ± 0.0571 | 0.58 ± 0.076 | 0.858 ± 0.00097 |
| Molformer | 0.808 ± 0.00388 | 0.817 ± 0.000195 | 0.936 ± 0.0593 | 0.546 ± 0.104 | 0.811 ± 0.0913 |
| UniMol-G | 0.806 ± 0.00119 | 0.817 ± 0.000519 | 0.89 ± 0.0661 | 0.57 ± 0.0952 | 0.85 ± 0.000406 |
| UniMol-GA | 0.808 ± 0.000863 | 0.817 ± 0.000326 | 0.947 ± 0.00218 | 0.731 ± 0.119 | 0.855 ± 0.000438 |
| GCC | 0.816 ± 0.00305 | 0.818 ± 0.000179 | 0.445 ± 0.124 | 0.554 ± 0.0593 | 0.523 ± 0.0781 |
| GPT-GNN | 0.811 ± 0.00368 | 0.819 ± 0.000406 | 0.902 ± 0.00475 | 0.94 ± 0.00263 | 0.817 ± 0.00518 |
| GraphCL | 10.8 ± 3.64 | 1.23 ± 0.53 | 0.611 ± 0.0472 | 0.618 ± 0.0834 | 0.665 ± 0.0811 |
| InfoGraph | 0.819 ± 0.00245 | 0.818 ± 0.000247 | 0.792 ± 0.00954 | 0.646 ± 0.155 | 0.746 ± 0.000755 |
| JOAO | 0.807 ± 0.00254 | 0.82 ± 0.000692 | 0.792 ± 0.0182 | 0.711 ± 0.00459 | 0.744 ± 0.00292 |
| JOAOv2 | 0.812 ± 0.00141 | 0.819 ± 0.000713 | 0.892 ± 0.0133 | 0.846 ± 0.0168 | 0.774 ± 0.018 |
| PosPred | 0.821 ± 0.00224 | 0.822 ± 0.00126 | 0.851 ± 0.00416 | 0.799 ± 0.00105 | 0.763 ± 0.000709 |
| 3D-EMGP | 0.823 ± 0.00736 | 0.909 ± 0.0888 | 0.643 ± 0.015 | 0.646 ± 0.0553 | 0.725 ± 0.0201 |
| 3D-InfoMax | 0.818 ± 0.00385 | 0.819 ± 0.000159 | 0.436 ± 0.144 | 0.64 ± 0.131 | 0.642 ± 0.142 |
| AttrMask | 2.8 ± 0.998 | 0.883 ± 0.0237 | 0.911 ± 0.00865 | 0.863 ± 0.0087 | 0.618 ± 0.0653 |
| EdgePred | 1.55 ± 0.984 | 0.818 ± 0.00142 | 0.504 ± 0.0469 | 0.56 ± 0.0824 | 0.669 ± 0.105 |
| GraphMVP | 0.807 ± 0.000227 | 0.817 ± 3.5e-06 | 0.5 ± 0 | 0.5 ± 0 | 0.5 ± 0 |
| MolCLR-GIN | 0.806 ± 0.00276 | 0.817 ± 0.000314 | 0.511 ± 0.0223 | 0.506 ± 0.0137 | 0.5 ± 0.0165 |
| MolCLR-GCN | 0.807 ± 0.00248 | 0.818 ± 0.000328 | 0.523 ± 0.0344 | 0.513 ± 0.0292 | 0.498 ± 0.00438 |

Prediction on the protein-ligand binding affinity is evaluated by RMSE. For each method, the mean and standard deviation are reported for the evaluation metrics. GROVER-Base, GROVER-Large, and MoleBERT are not evaluated here due to overflow in GPU memory.

**Supplementary Table 34.** Opioid-type receptors.

| Method | CYP2D6 | CYP3A4 | DOR | KOR | MDR1 | MOR |
| --- | --- | --- | --- | --- | --- | --- |
| qcGEM | 1.62 ± 0.0887 | 1.39 ± 0.0813 | 1.68 ± 0.0689 | 1.77 ± 0.0998 | 1.48 ± 0.231 | 1.77 ± 0.067 |
| ECFP | 1.75 ± 0.114 | 1.58 ± 0.065 | 1.7 ± 0.0715 | 1.79 ± 0.0881 | 1.79 ± 0.298 | 1.85 ± 0.0547 |
| FCFP | 1.7 ± 0.153 | 1.55 ± 0.109 | 1.79 ± 0.0453 | 1.8 ± 0.119 | 1.75 ± 0.227 | 1.8 ± 0.0681 |
| AvalonFP | 1.66 ± 0.126 | 1.5 ± 0.0809 | 1.77 ± 0.0397 | 1.9 ± 0.0704 | 1.55 ± 0.275 | 1.82 ± 0.0931 |
| TTGEN | 1.74 ± 0.15 | 1.64 ± 0.0995 | 1.8 ± 0.104 | 1.85 ± 0.0641 | 1.74 ± 0.34 | 1.89 ± 0.072 |
| APGEN | 1.72 ± 0.152 | 1.61 ± 0.091 | 1.9 ± 0.116 | 1.99 ± 0.0758 | 1.87 ± 0.169 | 2.07 ± 0.114 |
| RDKFP | 1.68 ± 0.161 | 1.56 ± 0.0928 | 2.02 ± 0.0825 | 2.07 ± 0.058 | 1.71 ± 0.22 | 2.03 ± 0.11 |
| MACCS | 1.76 ± 0.165 | 1.72 ± 0.096 | 2.05 ± 0.0803 | 2.18 ± 0.0666 | 1.92 ± 0.325 | 2.2 ± 0.0874 |
| LayeredFP | 1.72 ± 0.161 | 1.81 ± 0.0908 | 2.01 ± 0.0965 | 2.2 ± 0.0975 | 1.76 ± 0.23 | 2.21 ± 0.113 |
| PatternFP | 1.73 ± 0.0975 | 1.68 ± 0.105 | 2.15 ± 0.0756 | 2.21 ± 0.0754 | 1.72 ± 0.245 | 2.16 ± 0.105 |
| Molformer | 1.6 ± 0.151 | 1.5 ± 0.111 | 1.75 ± 0.101 | 1.76 ± 0.112 | 1.57 ± 0.191 | 1.81 ± 0.0712 |
| GROVER-Base | 1.61 ± 0.109 | 1.43 ± 0.111 | 1.9 ± 0.0948 | 2.11 ± 0.151 | 1.54 ± 0.235 | 2.09 ± 0.108 |
| GROVER-Large | 1.64 ± 0.155 | 1.51 ± 0.104 | 1.95 ± 0.0928 | 2.09 ± 0.0546 | 1.64 ± 0.189 | 2.03 ± 0.149 |
| MoleBERT | 1.78 ± 0.164 | 1.87 ± 0.0971 | 2.31 ± 0.1 | 2.28 ± 0.0545 | 1.88 ± 0.163 | 2.36 ± 0.142 |
| UniMol-G | 1.8 ± 0.172 | 1.84 ± 0.0935 | 2.38 ± 0.104 | 2.44 ± 0.0536 | 2.06 ± 0.191 | 2.44 ± 0.0894 |
| UniMol-GA | 1.78 ± 0.171 | 1.79 ± 0.0939 | 2.29 ± 0.0923 | 2.37 ± 0.0484 | 1.96 ± 0.224 | 2.35 ± 0.105 |
| GCC | 1.83 ± 0.195 | 1.87 ± 0.0764 | 2.37 ± 0.058 | 2.43 ± 0.0514 | 2.08 ± 0.165 | 2.43 ± 0.167 |
| GPT-GNN | 1.91 ± 0.166 | 2.01 ± 0.106 | 2.82 ± 0.081 | 2.71 ± 0.0259 | 2.34 ± 0.251 | 2.7 ± 0.12 |
| GraphCL | 1.91 ± 0.191 | 2.02 ± 0.12 | 2.96 ± 0.119 | 2.8 ± 0.0453 | 2.46 ± 0.13 | 2.83 ± 0.143 |
| InfoGraph | 1.92 ± 0.178 | 2.02 ± 0.136 | 2.88 ± 0.1 | 2.8 ± 0.0531 | 2.7 ± 0.129 | 2.8 ± 0.13 |
| JOAO | 1.93 ± 0.183 | 2.02 ± 0.125 | 2.87 ± 0.103 | 2.74 ± 0.0365 | 2.61 ± 0.214 | 2.79 ± 0.15 |
| JOAOv2 | 1.86 ± 0.171 | 1.98 ± 0.135 | 2.82 ± 0.115 | 2.74 ± 0.0591 | 2.58 ± 0.171 | 2.74 ± 0.153 |
| PosPred | 1.87 ± 0.168 | 1.98 ± 0.121 | 2.73 ± 0.126 | 2.48 ± 0.0815 | 2.29 ± 0.0951 | 2.63 ± 0.13 |
| 3D-EMGP | 1.96 ± 0.174 | 2.01 ± 0.135 | 2.99 ± 0.137 | 2.84 ± 0.0654 | 2.64 ± 0.0995 | 2.85 ± 0.159 |
| 3D-InfoMax | 1.65 ± 0.132 | 1.59 ± 0.0899 | 1.89 ± 0.125 | 1.89 ± 0.0442 | 1.62 ± 0.222 | 2 ± 0.064 |
| AttrMask | 1.95 ± 0.183 | 2.06 ± 0.133 | 2.92 ± 0.0837 | 2.68 ± 0.0635 | 2.61 ± 0.176 | 2.77 ± 0.154 |
| EdgePred | 1.95 ± 0.187 | 2.05 ± 0.133 | 3.08 ± 0.119 | 3.02 ± 0.0423 | 2.84 ± 0.219 | 2.91 ± 0.136 |
| GraphMVP | 1.98 ± 0.183 | 2.07 ± 0.133 | 3.13 ± 0.116 | 3.07 ± 0.0532 | 2.95 ± 0.255 | 2.93 ± 0.127 |
| MolCLR-GIN | 1.98 ± 0.183 | 2.07 ± 0.135 | 3.12 ± 0.122 | 3.08 ± 0.0385 | 2.96 ± 0.25 | 2.93 ± 0.129 |
| MolCLR-GCN | 1.98 ± 0.178 | 2.07 ± 0.134 | 3.13 ± 0.109 | 3.08 ± 0.0521 | 2.95 ± 0.253 | 2.93 ± 0.128 |

Prediction on the protein-ligand binding affinity is evaluated by RMSE. For each method, the mean and standard deviation are reported for the evaluation metrics.

#### 3.4 Model fine-tuning

##### 3.4.1 Experimental setting

**Supplementary Table 35.**
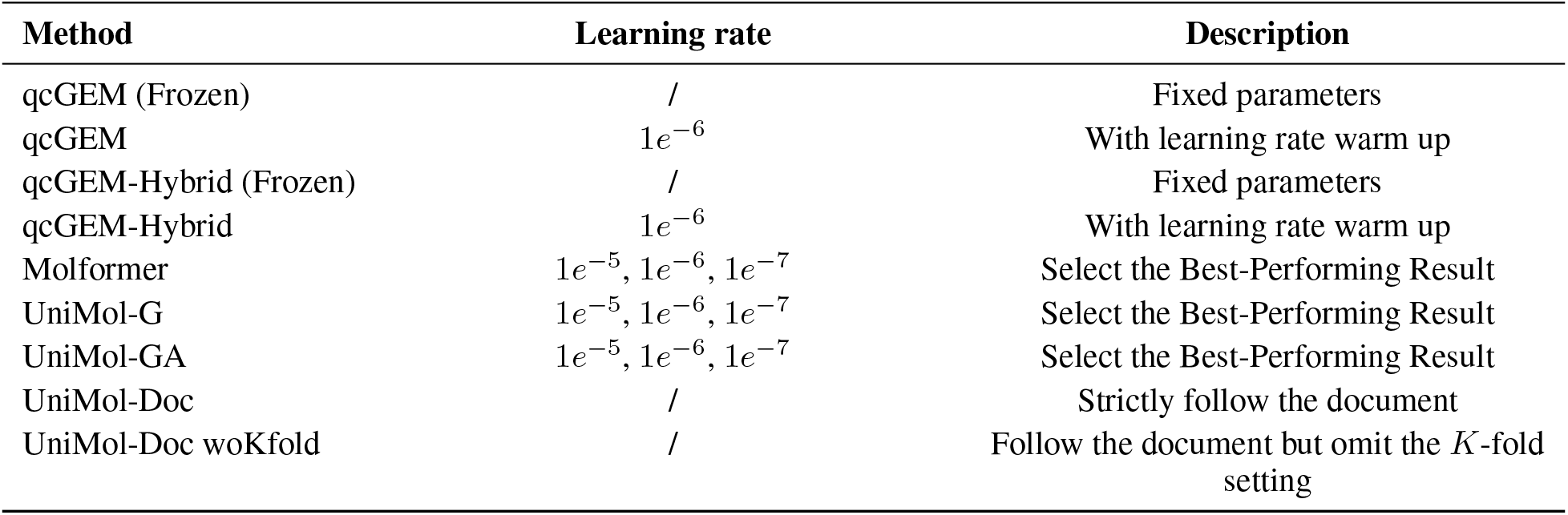
Hyper-parameter settings of fine-tuning experiments.

| Method | Learning rate | Description |
| --- | --- | --- |
| qcGEM (Frozen) | / | Fixed parameters |
| qcGEM | $1e^{-6}$ | With learning rate warm up |
| qcGEM-Hybrid (Frozen) | / | Fixed parameters |
| qcGEM-Hybrid | $1e^{-6}$ | With learning rate warm up |
| Molformer | $1e^{-5}, 1e^{-6}, 1e^{-7}$ | Select the Best-Performing Result |
| UniMol-G | $1e^{-5}, 1e^{-6}, 1e^{-7}$ | Select the Best-Performing Result |
| UniMol-GA | $1e^{-5}, 1e^{-6}, 1e^{-7}$ | Select the Best-Performing Result |
| UniMol-Doc | / | Strictly follow the document |
| UniMol-Doc woKfold | / | Follow the document but omit the $K$-fold setting |

##### 3.4.2 Experimental results

**Supplementary Table 36.** Evaluatsion results of fine-tuning experiments.

| Method | MoleculeACE (30 subsets) |
| --- | --- |
| qcGEM (Frozen) | $0.8106 \pm 0.1106$ |
| qcGEM | $0.8021 \pm 0.1158$ |
| qcGEM-Hybrid (Frozen) | $0.8110 \pm 0.1160$ |
| qcGEM-Hybrid | $0.8053 \pm 0.1172$ |
| Molformer | $0.9253 \pm 0.1066$ |
| UniMol-G | $1.0513 \pm 0.1725$ |
| UniMol-GA | $1.0304 \pm 0.1646$ |
| UniMol-Doc | $0.8281 \pm 0.1149$ |
| UniMol-Doc woKfold | $0.9805 \pm 0.1176$ |

Fine-tuning performance is evaluated by RMSE. For each method, the mean and standard deviation over the 30 datasets from MoleculeACE are reported for the evaluation metric.

### 4 Pseudo Codes and Algorithms

#### 4.1 Algorithms of Model

##### Algorithm 1: qcGEM

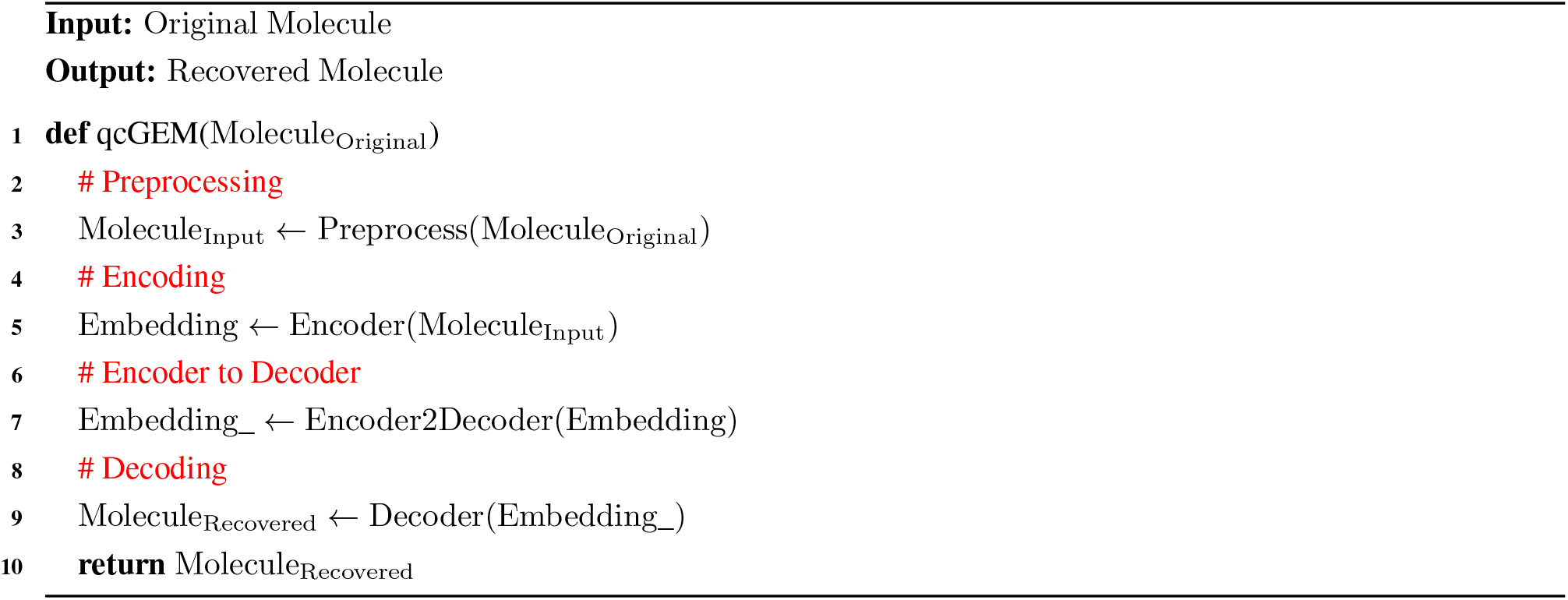

##### Algorithm 2: Preprocess

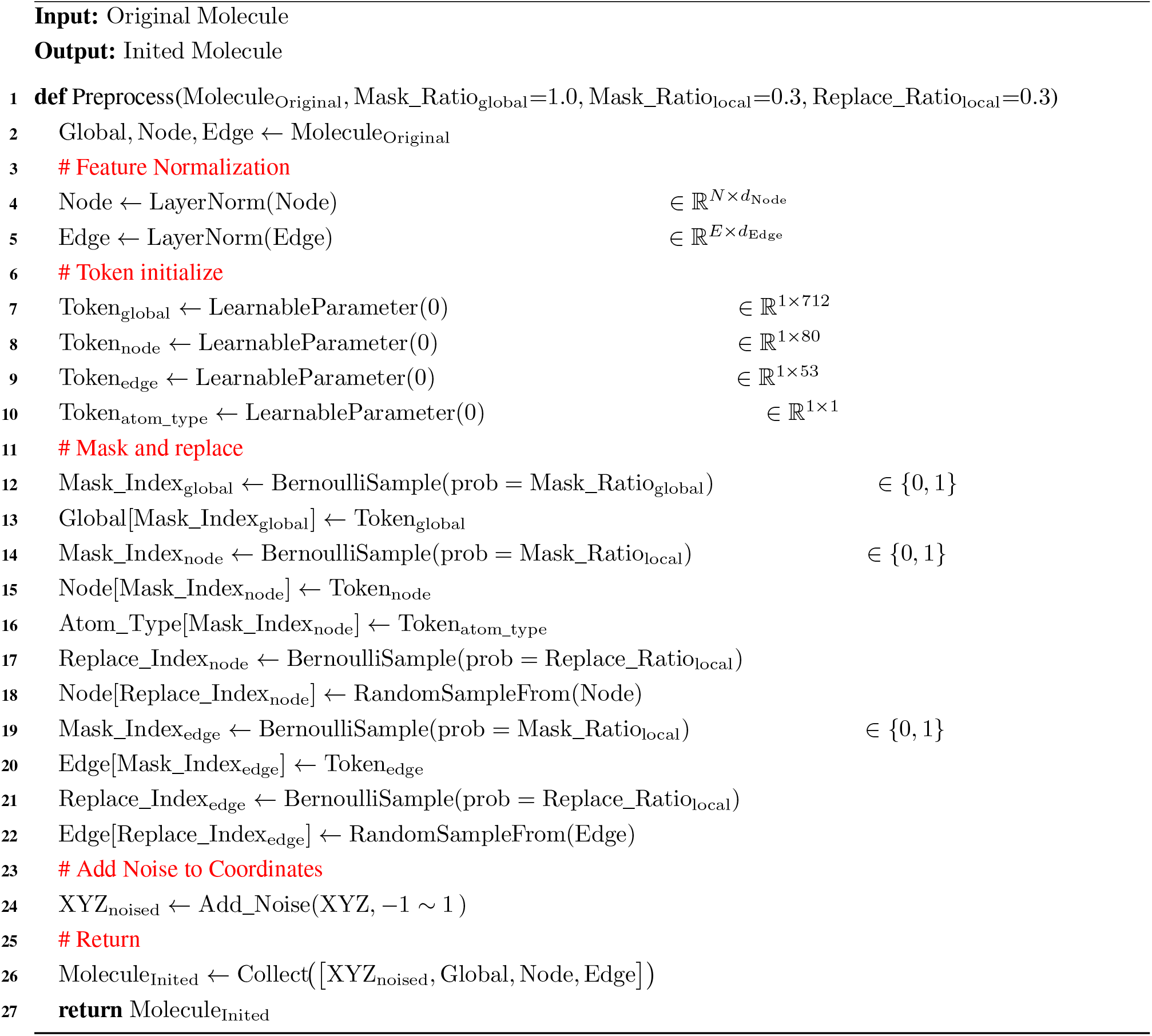

##### Algorithm 3: Encoder

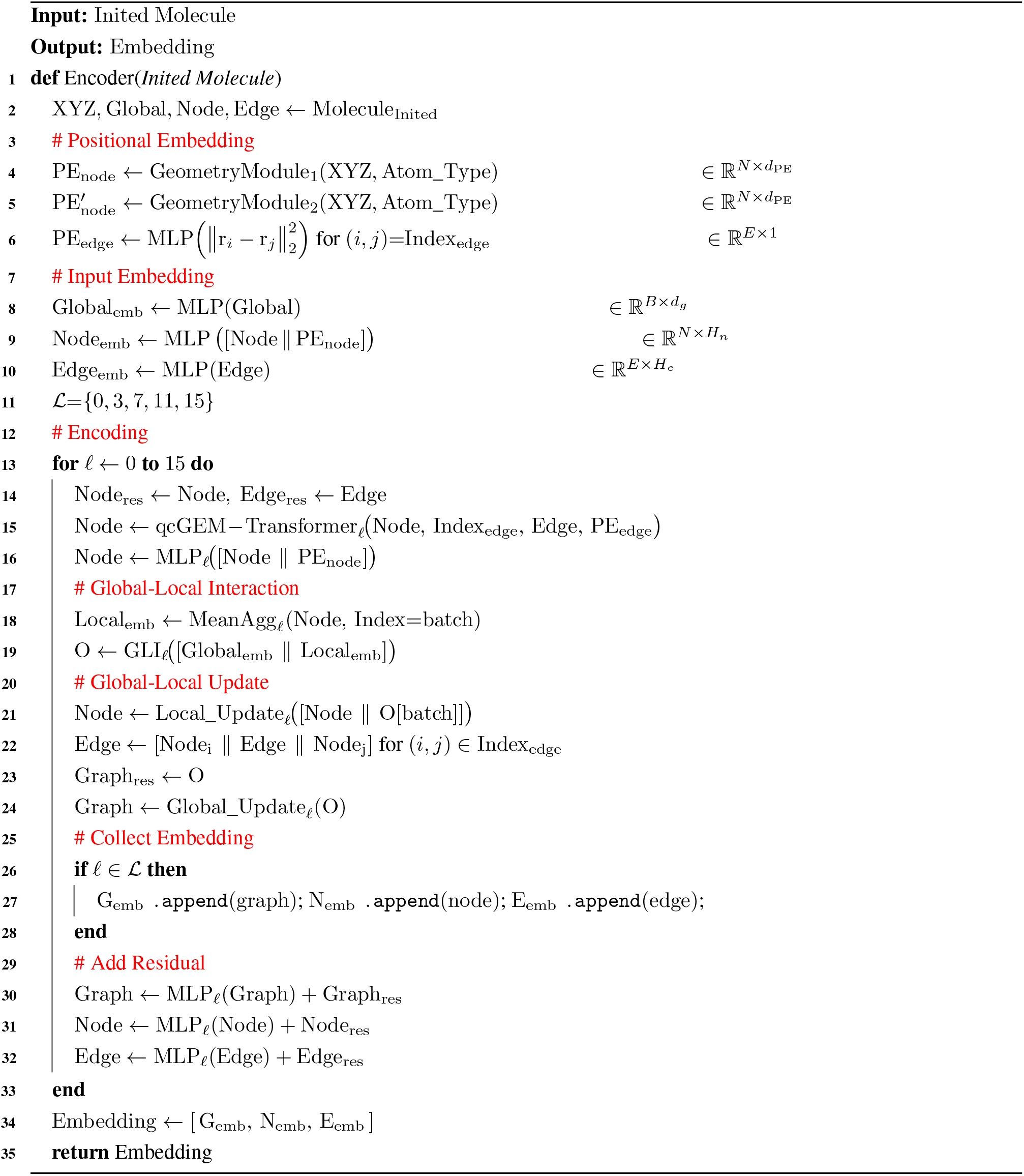

##### Algorithm 4: Encoder to Decoder

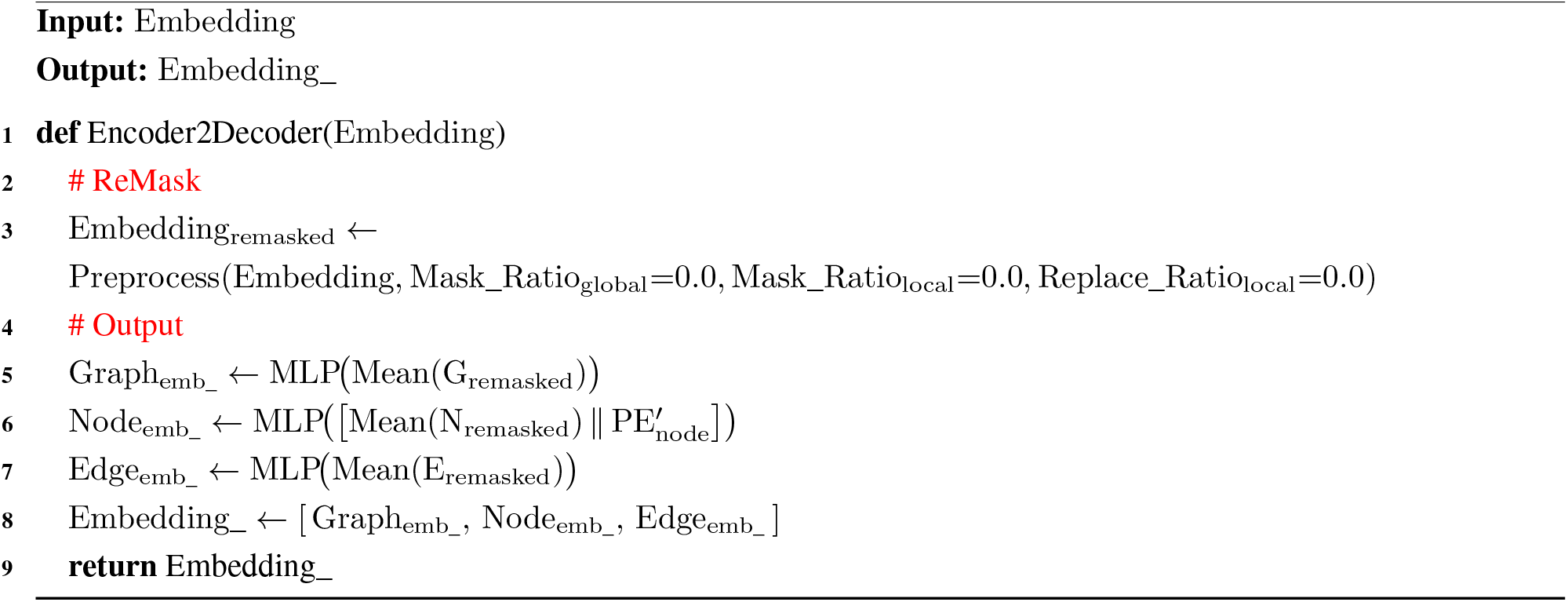

##### Algorithm 5: Decoder

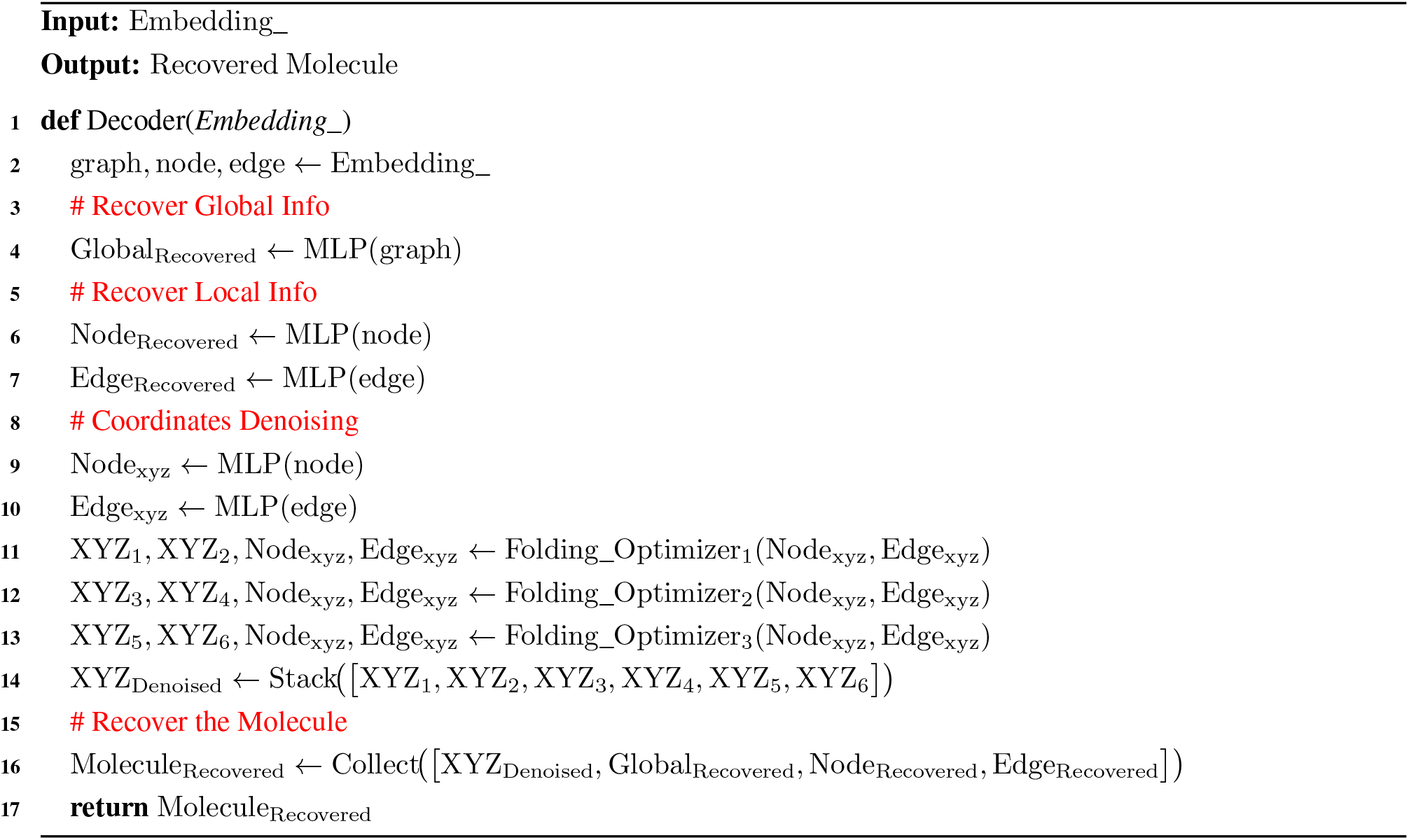

##### Algorithm 6: Geometry Module

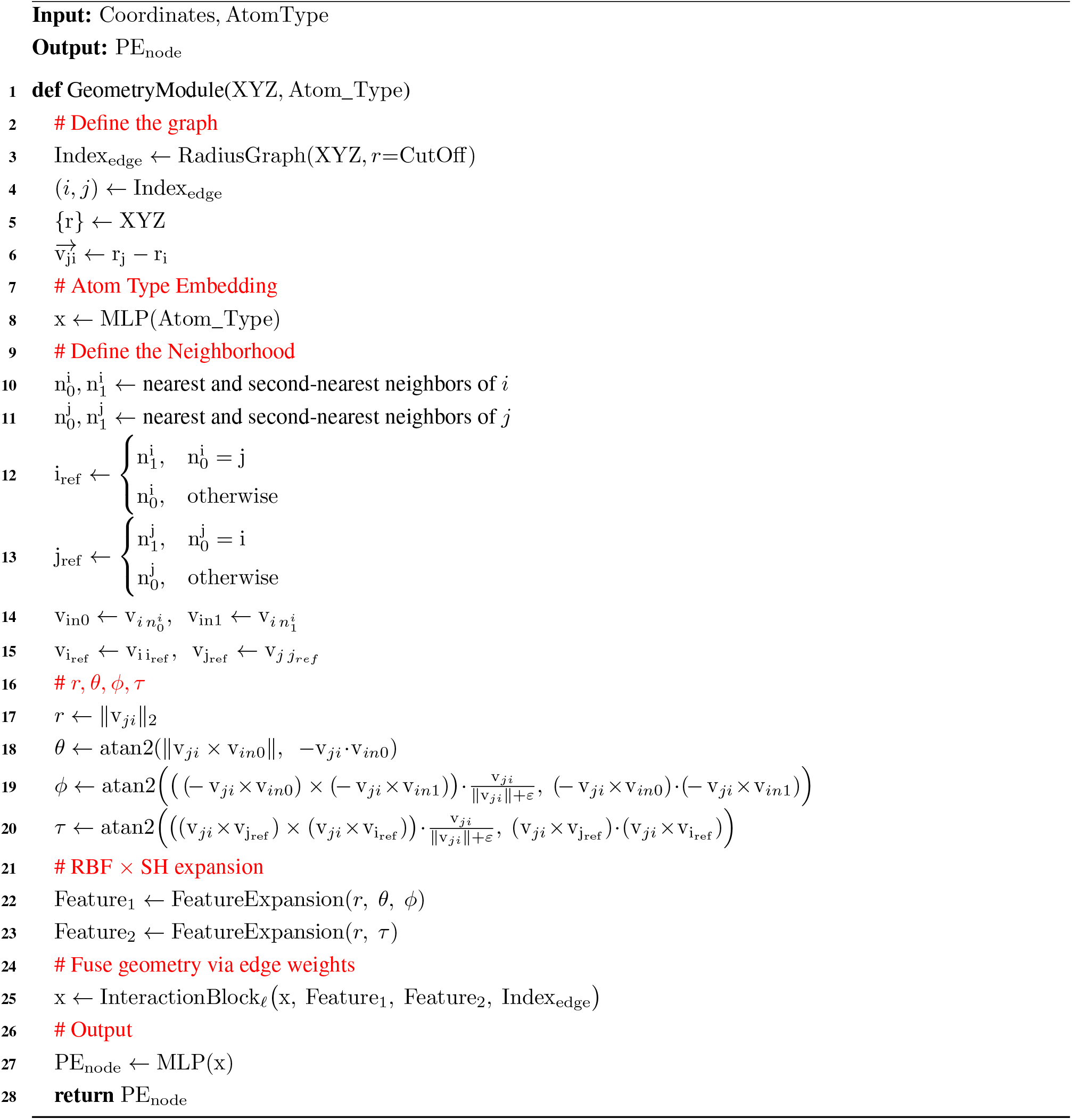

##### Algorithm 7: qcGEM-Transformer

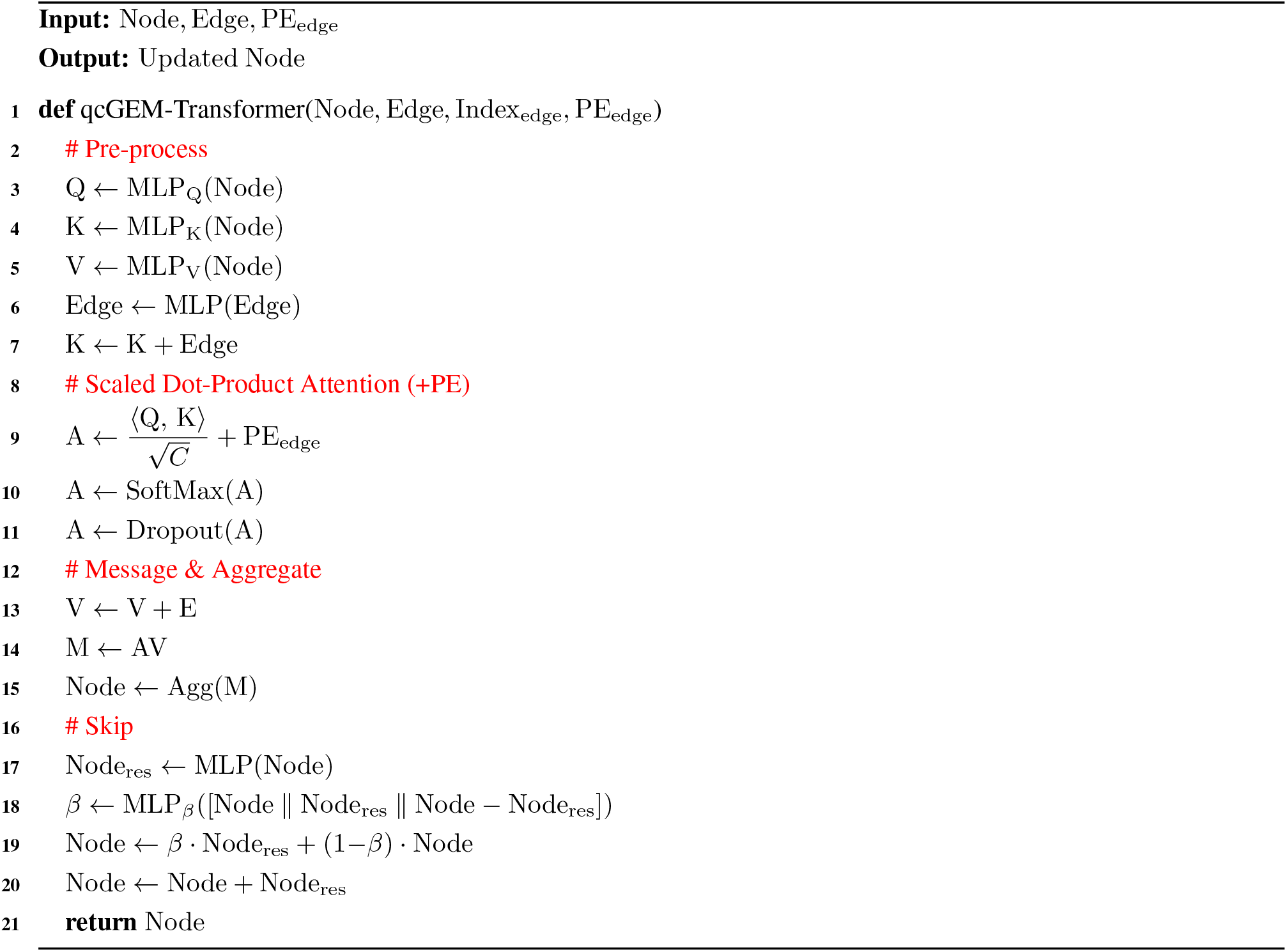

##### Algorithm 8: GLI

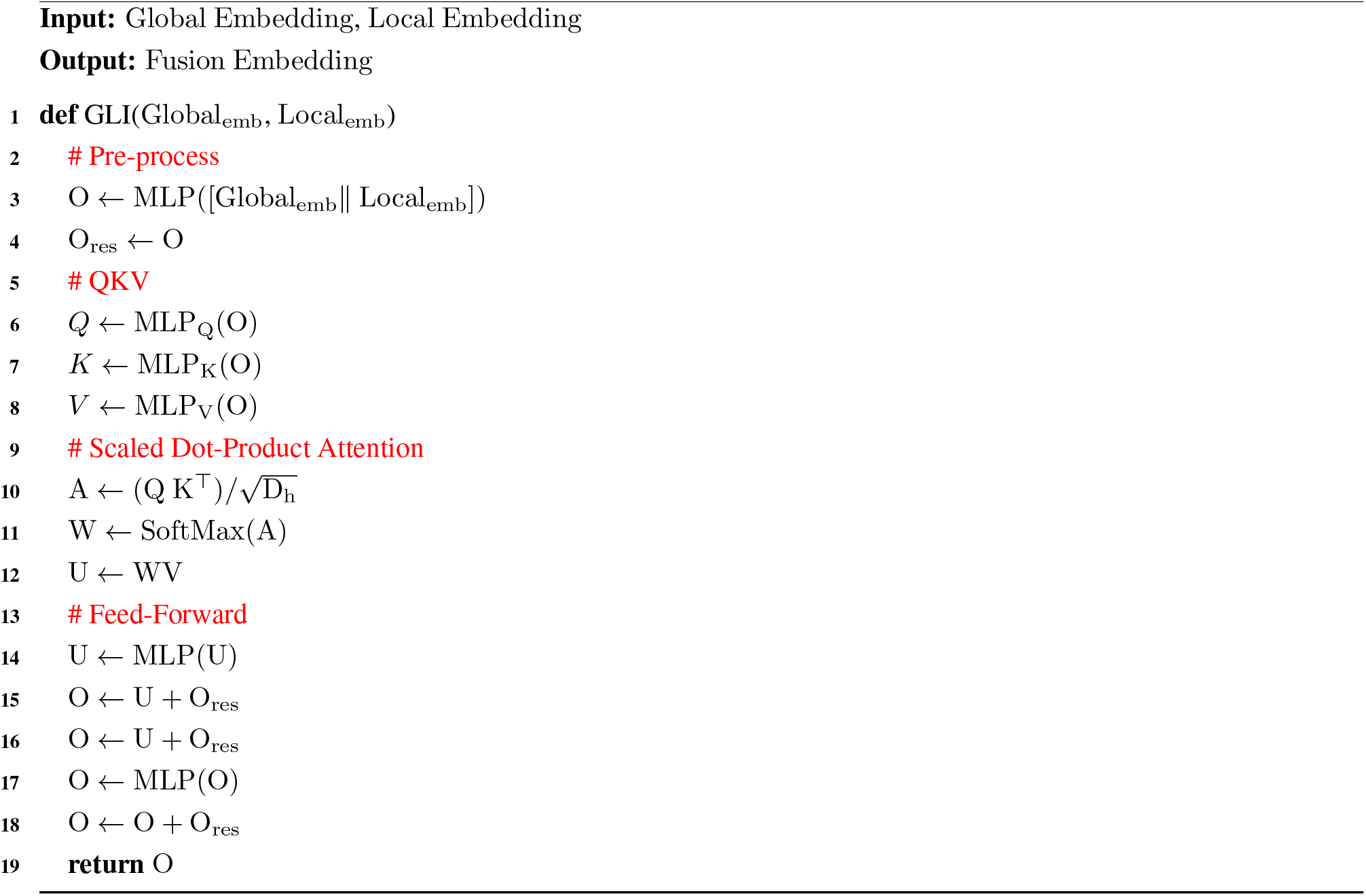

##### Algorithm 9: Global/Local Update

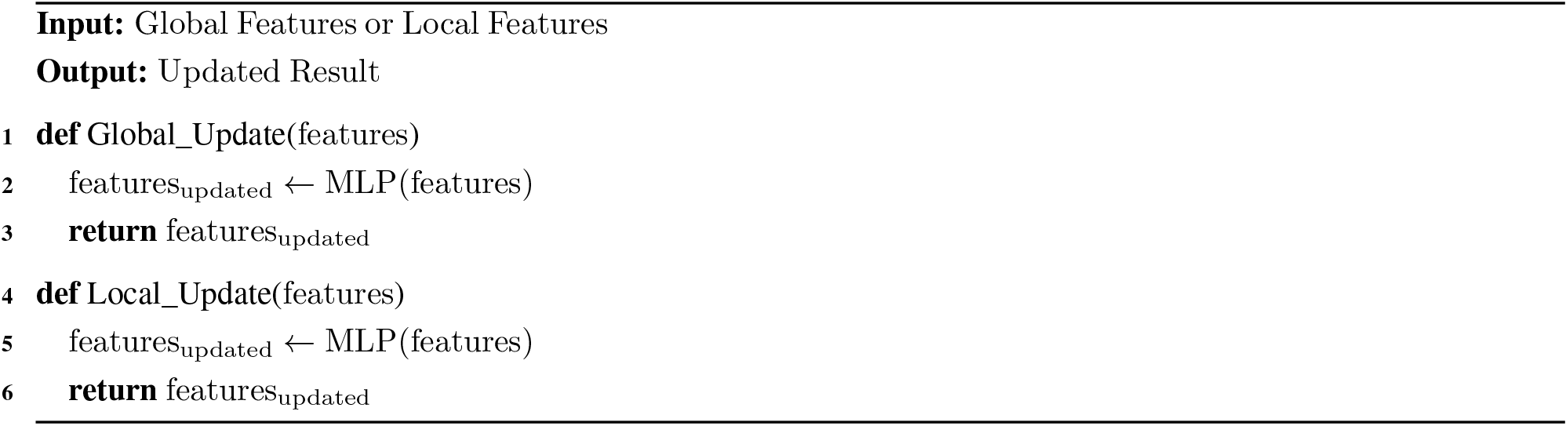

##### Algorithm 10: InteractionBlock

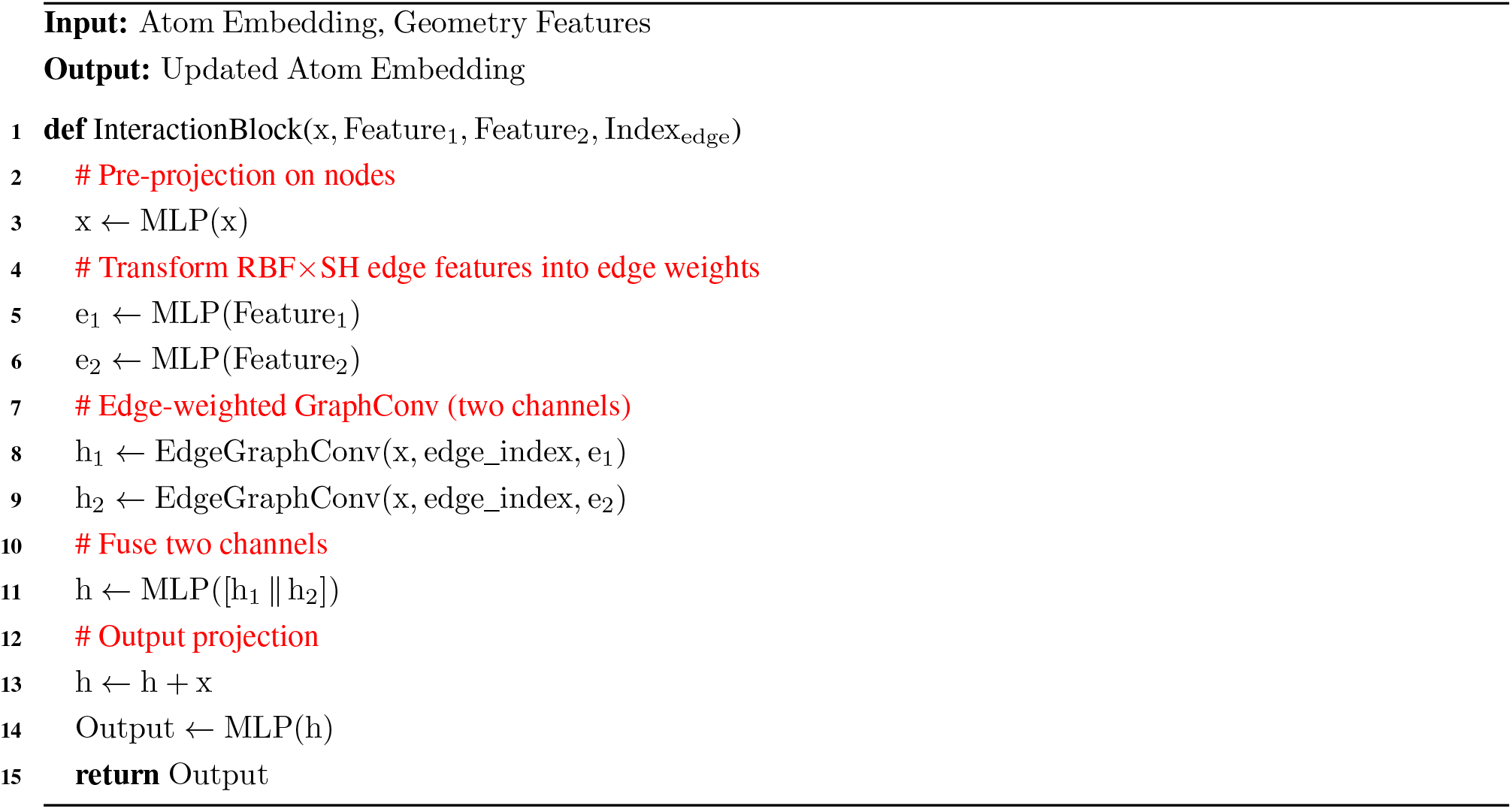

##### Algorithm 11: EdgeGraphConv

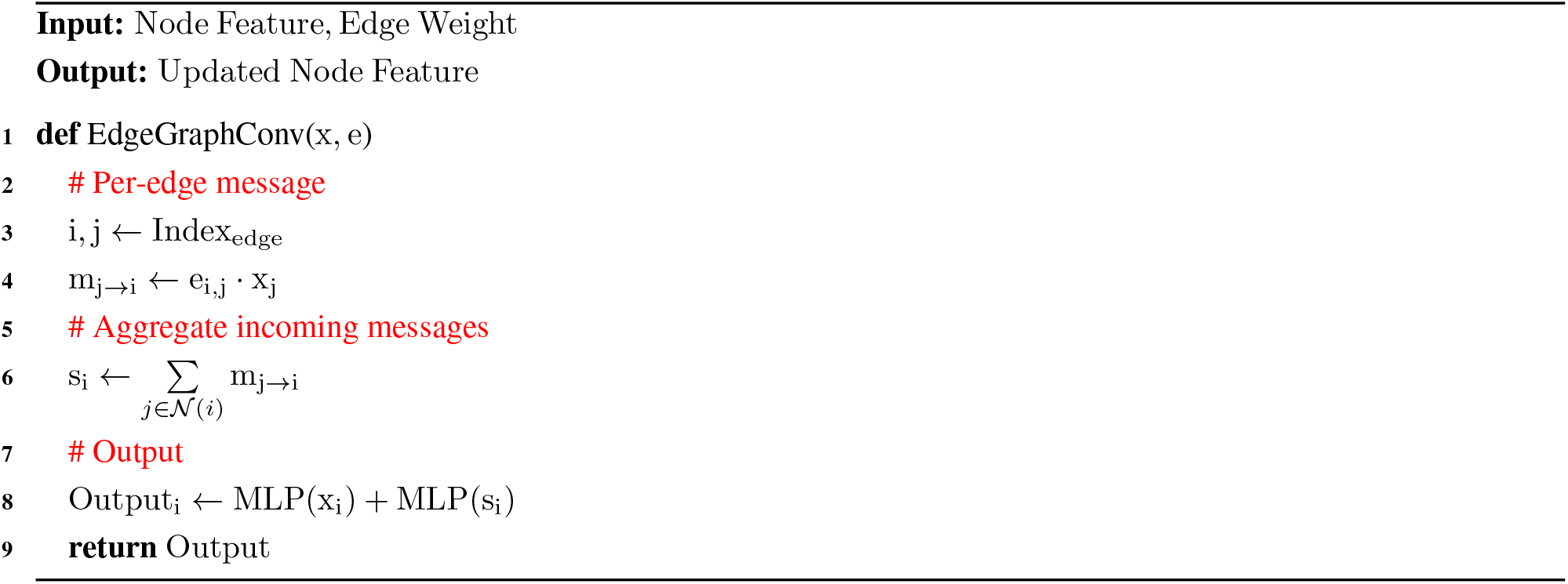

##### Algorithm 12: Folding_Optimizer

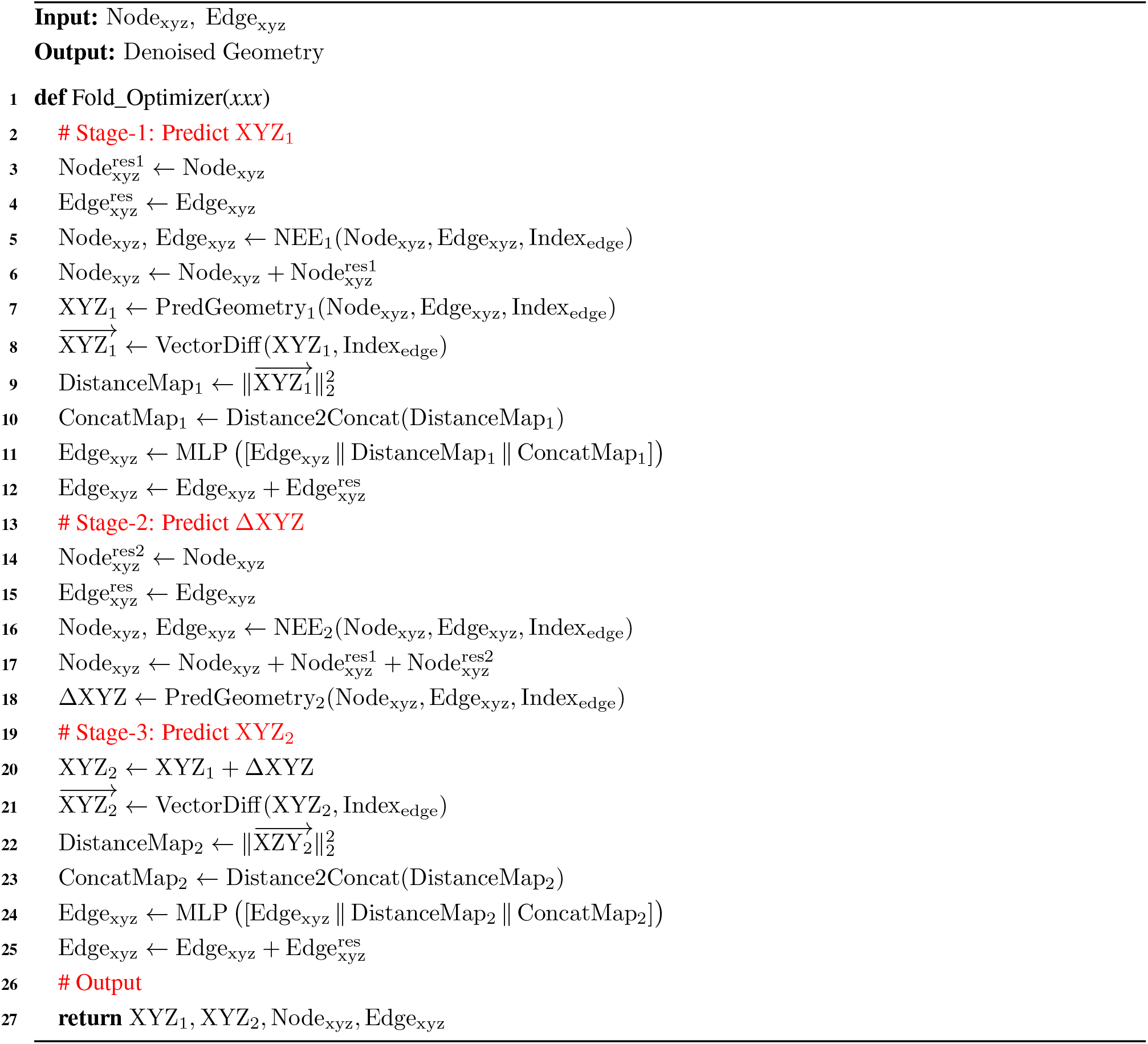

##### Algorithm 13: Distance2Concat

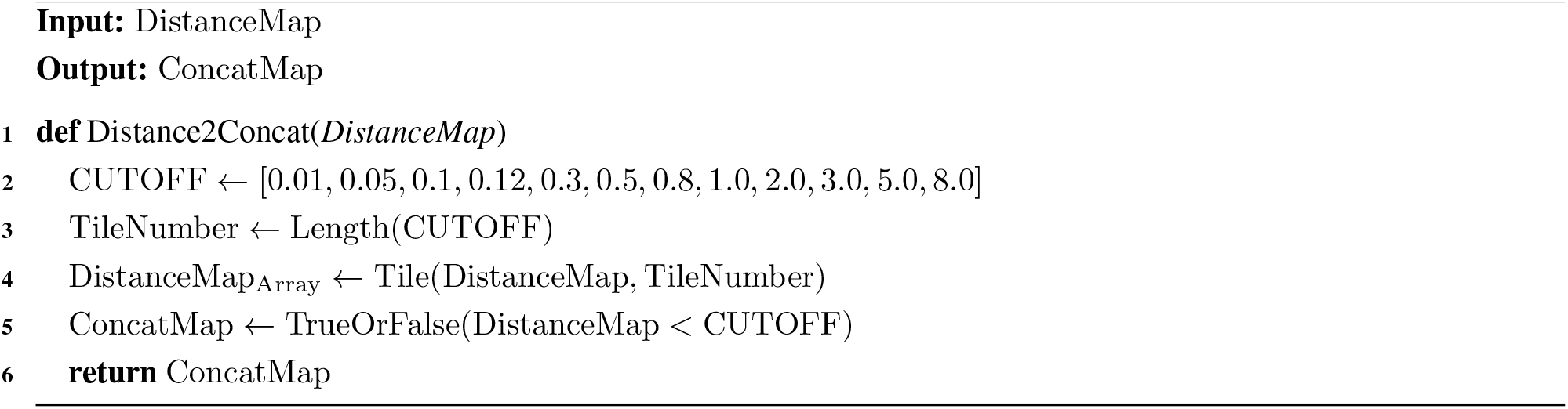

##### Algorithm 14: PredGeometry

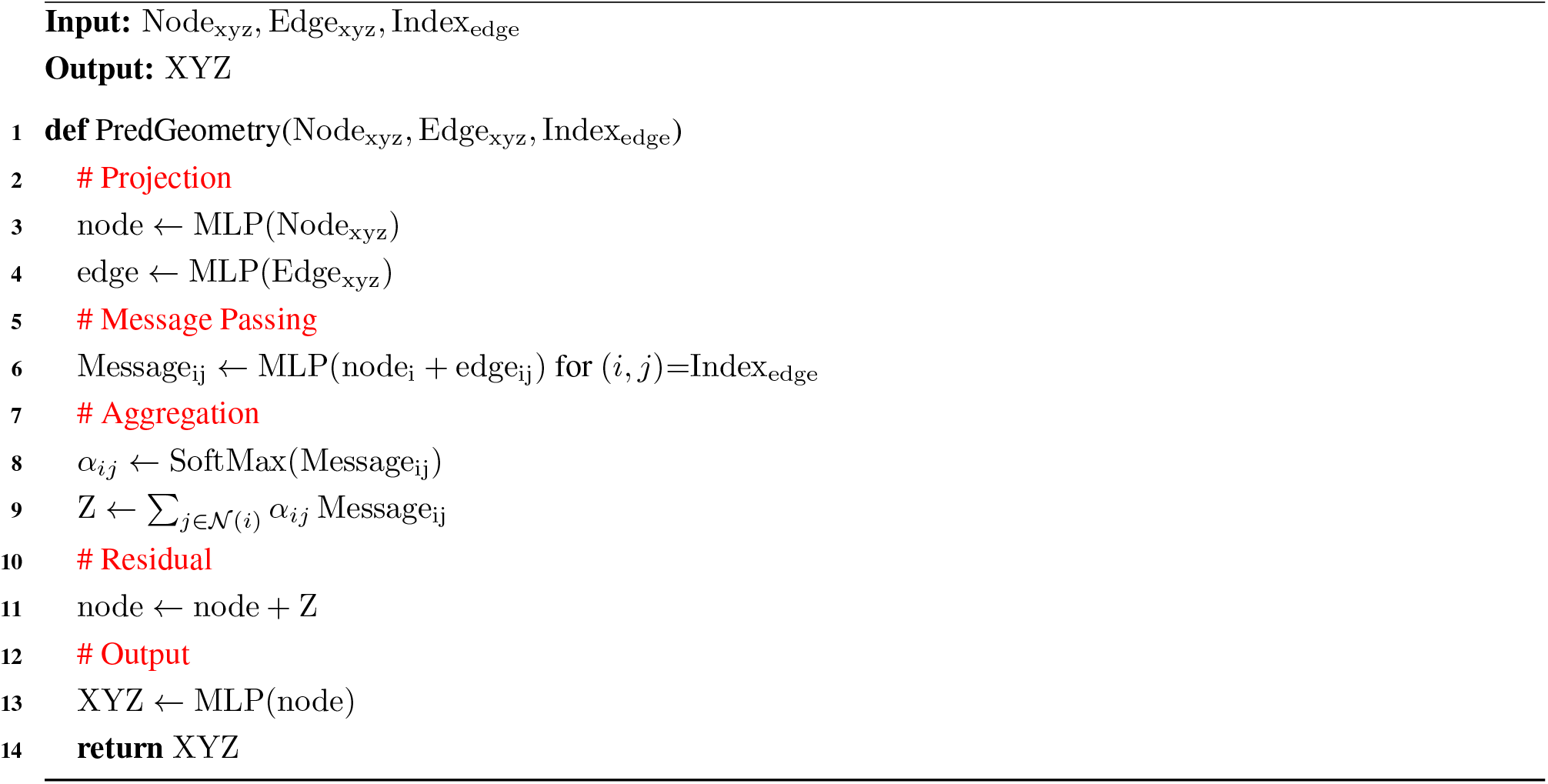

##### Algorithm 15: VectorDiff

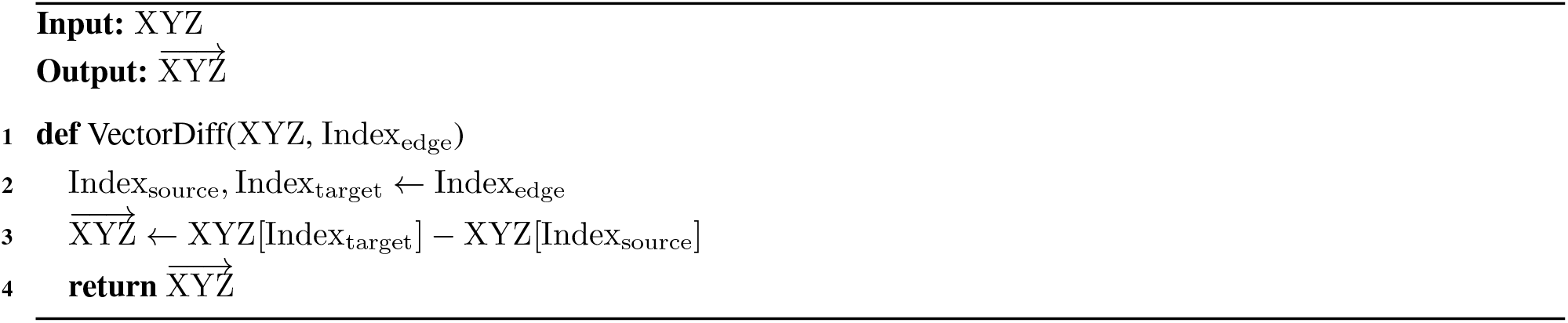

##### Algorithm 16: NEE

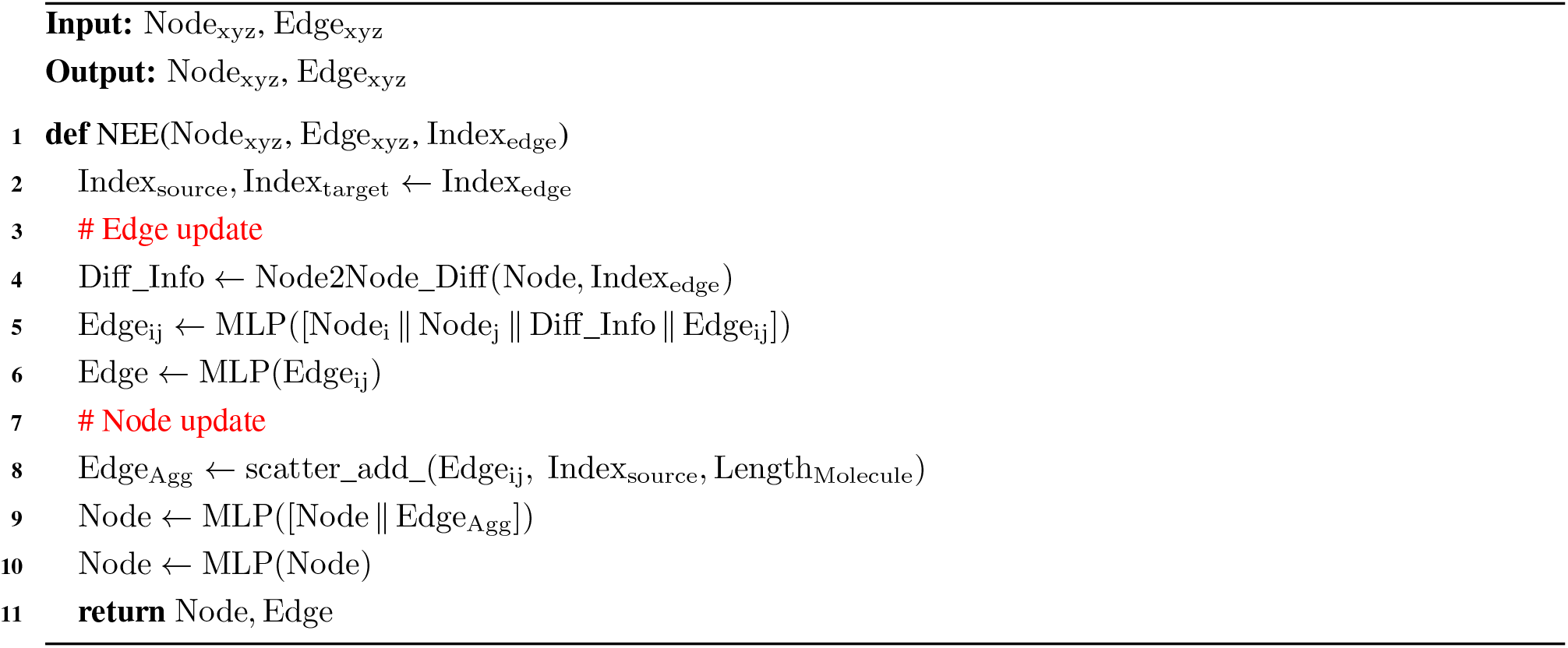

##### Algorithm 17: Node2Node_Diff

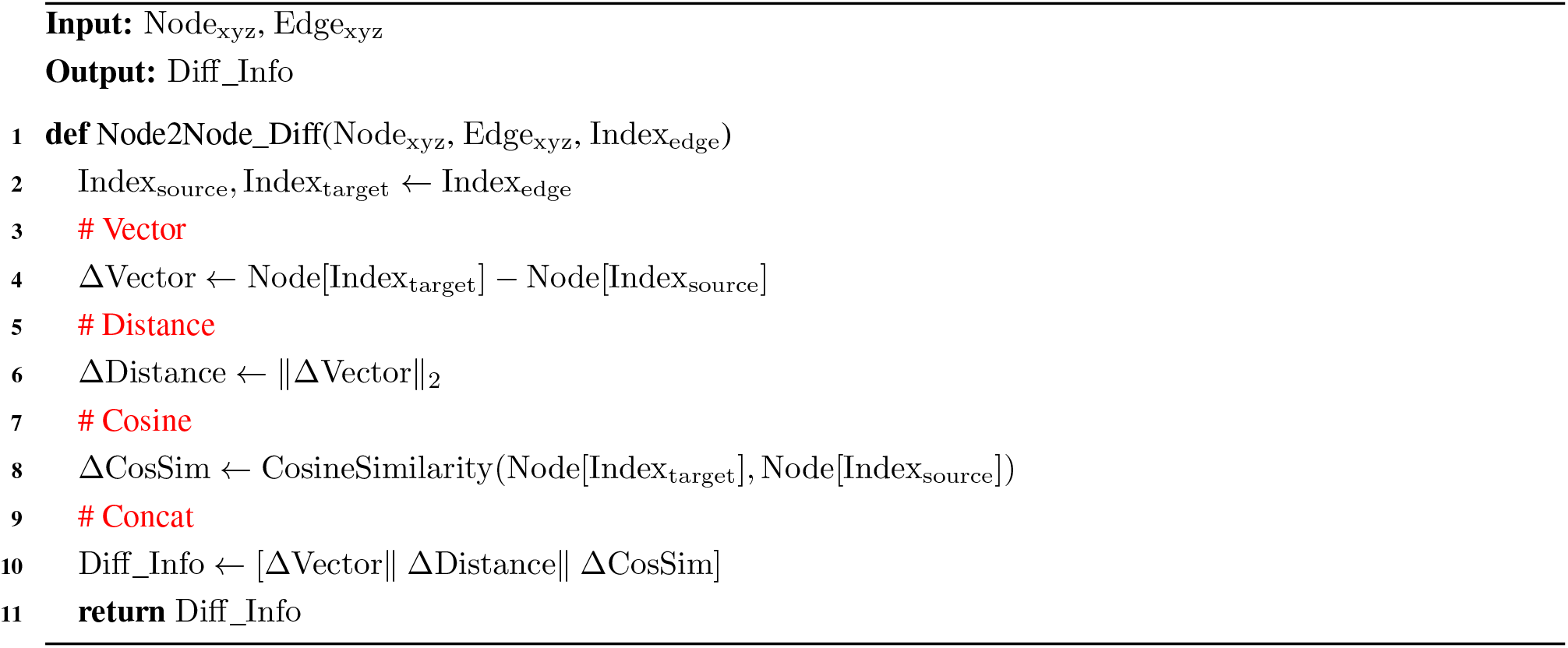

#### 4.2 Loss function for model training

##### Algorithm 18: Total Loss

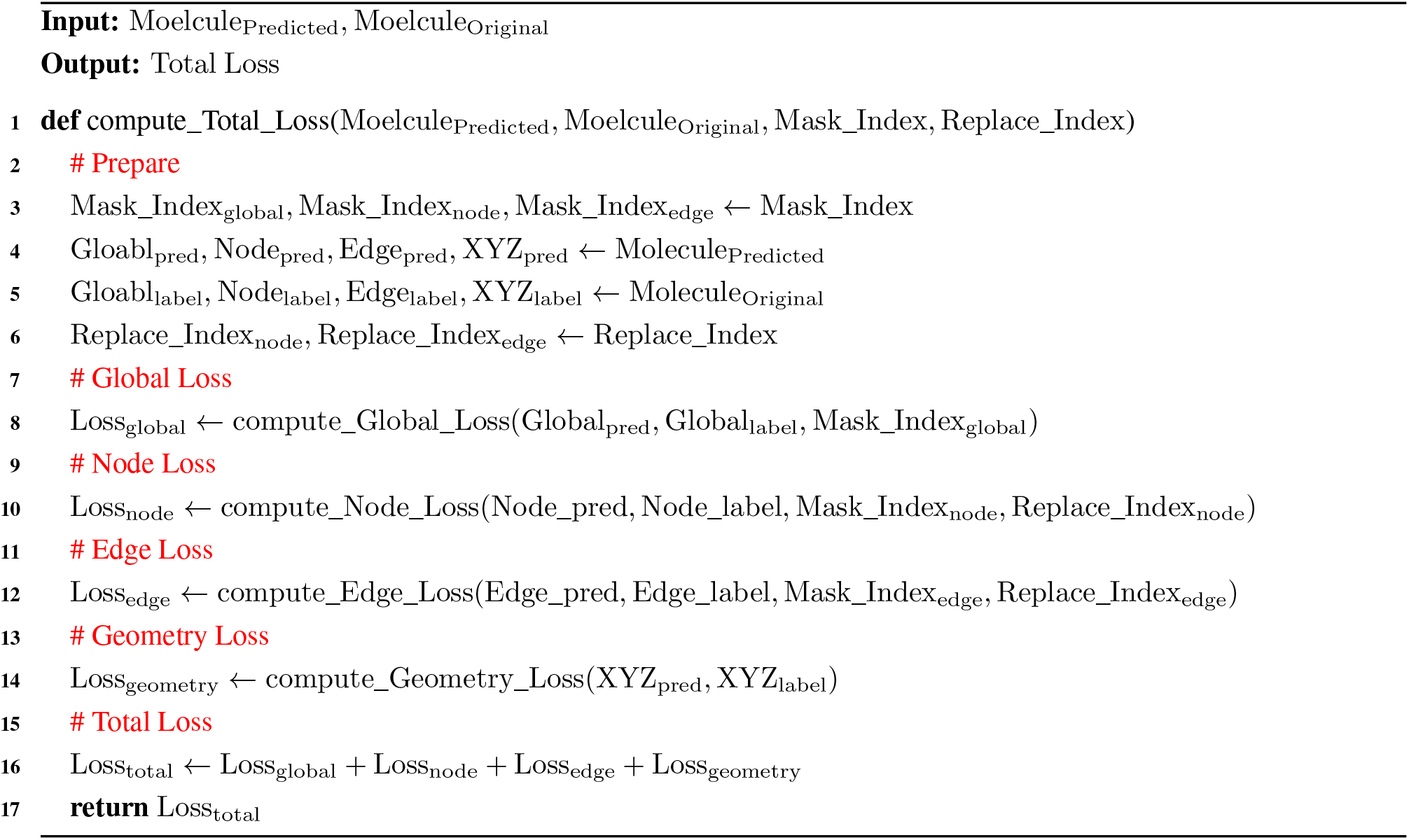

##### Algorithm 19: Global Loss

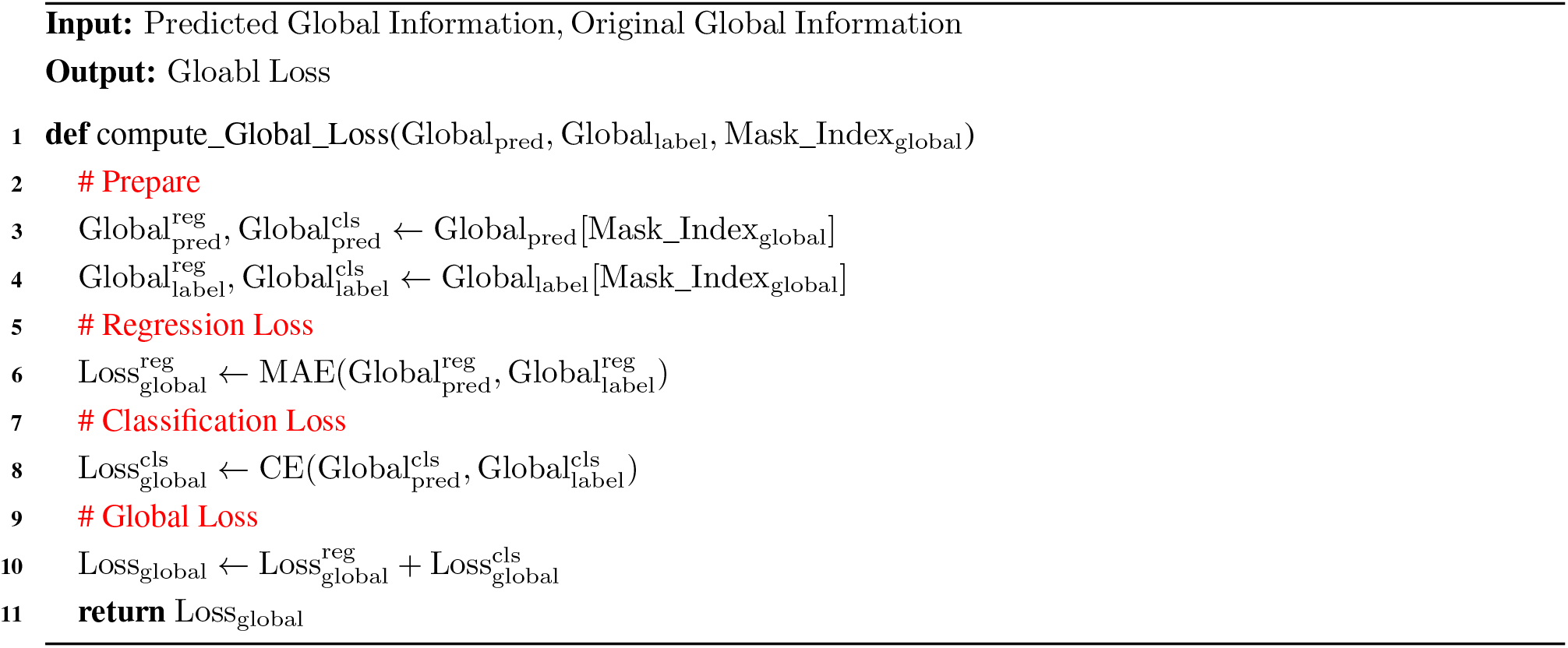

##### Algorithm 20: Node Loss

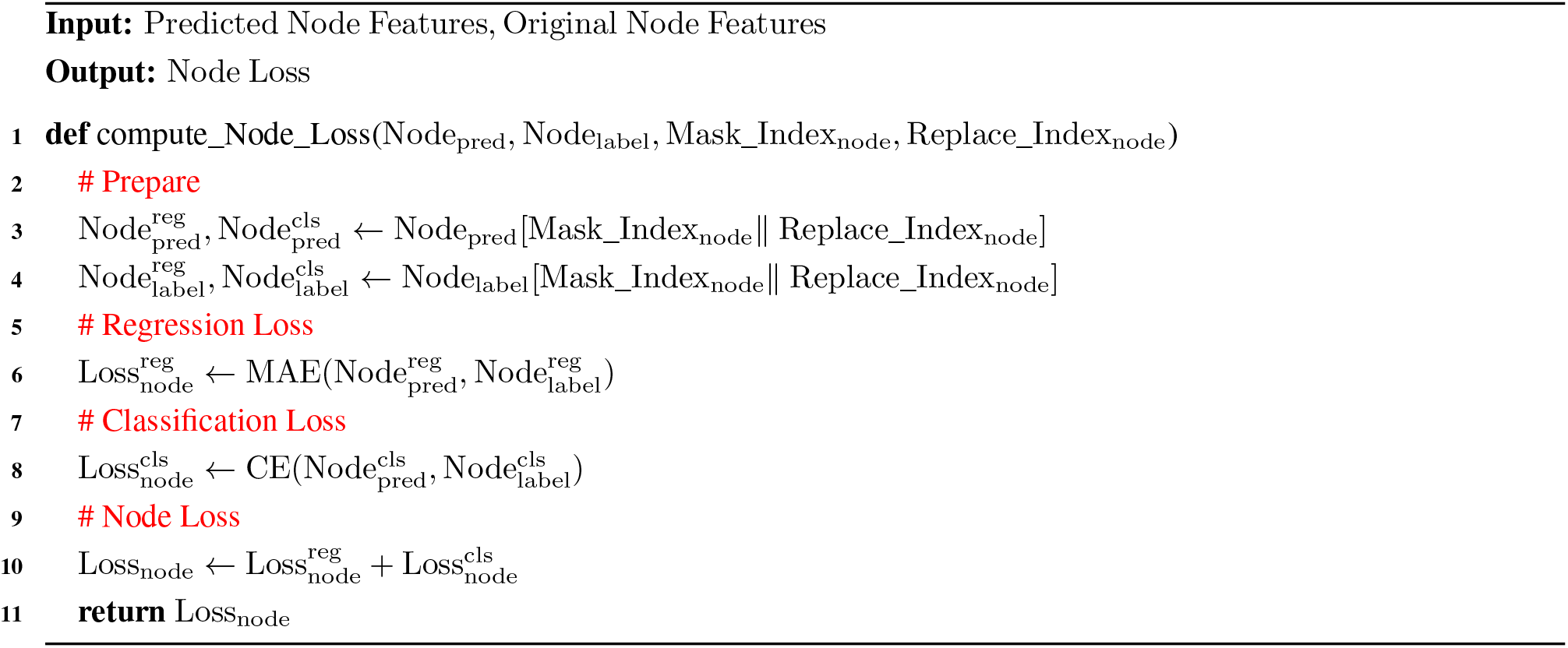

##### Algorithm 21: Edge Loss

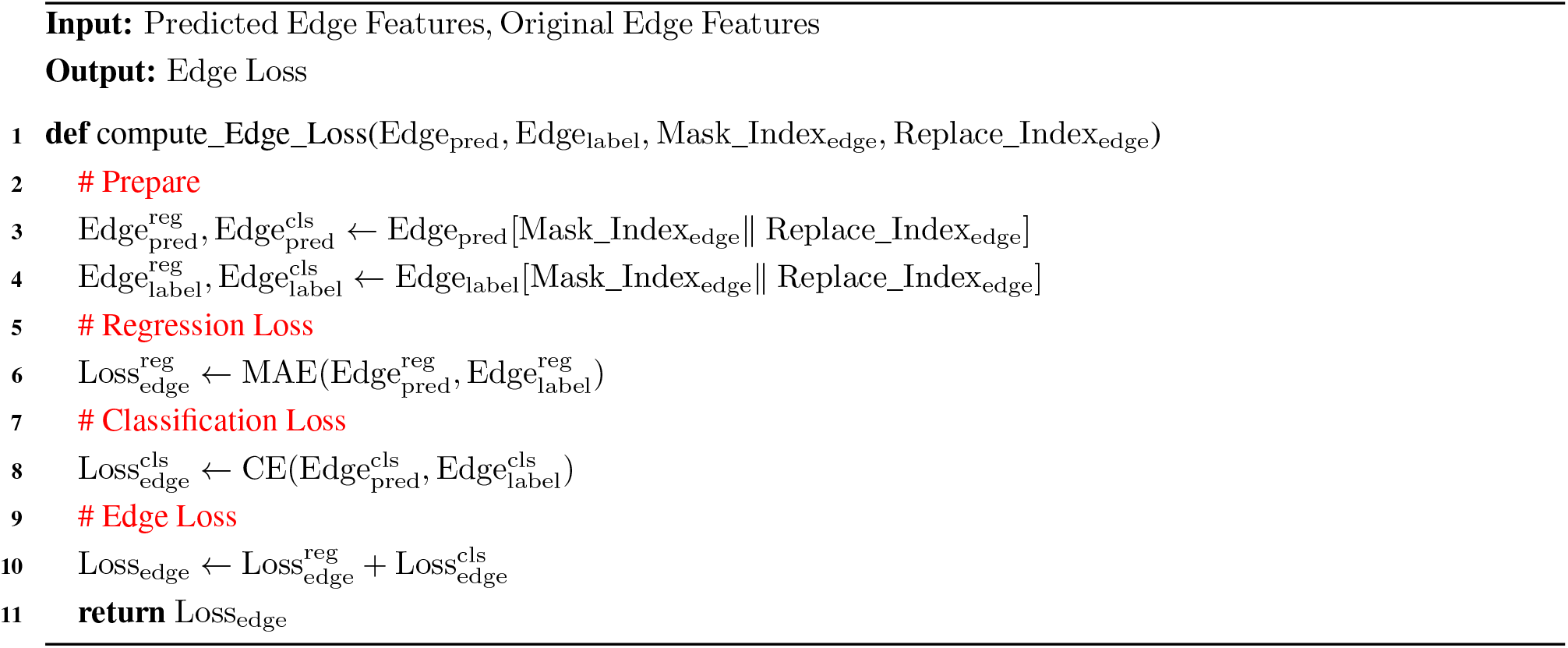

##### Algorithm 22: Geometry Loss

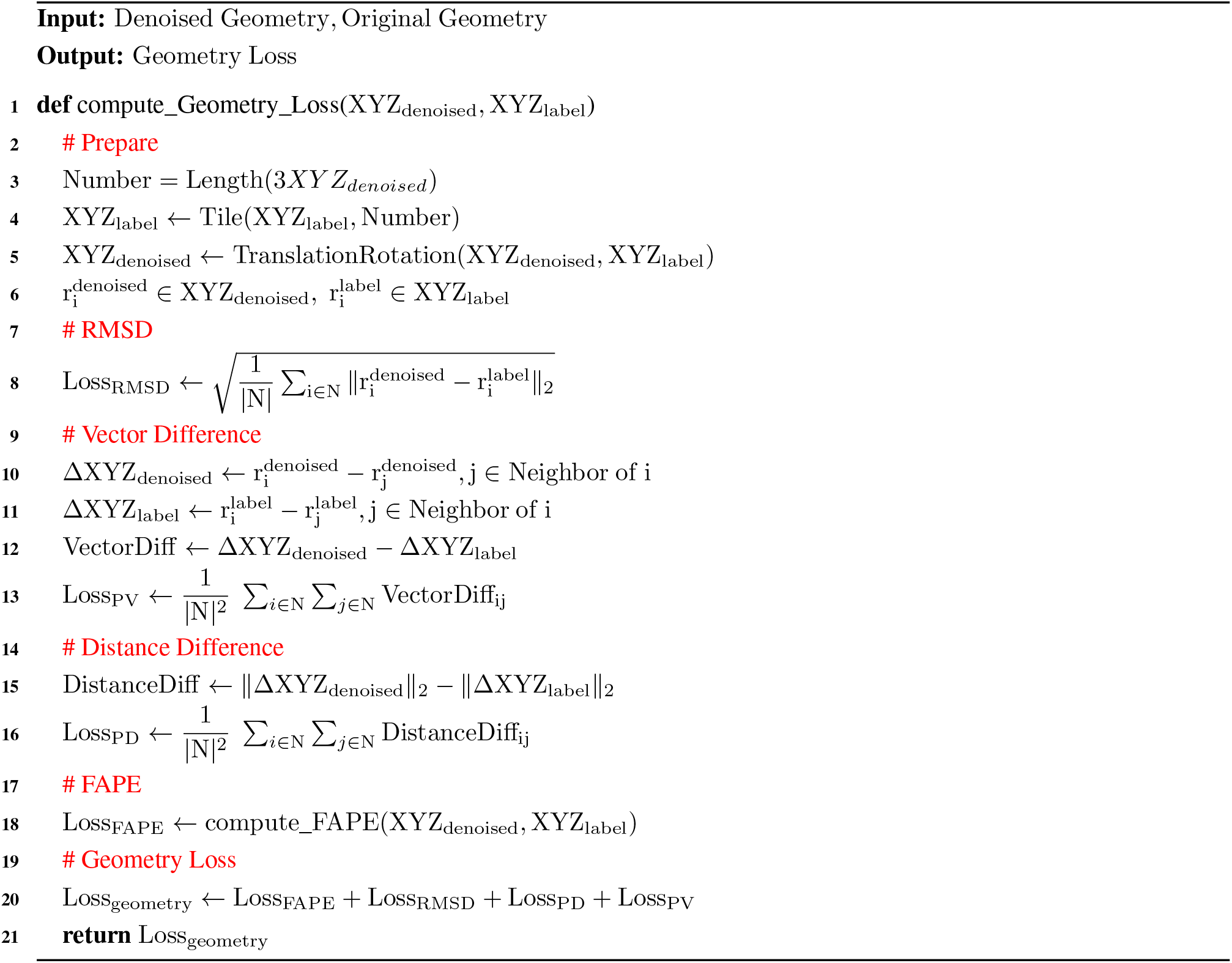

##### Algorithm 23: FAPE Loss

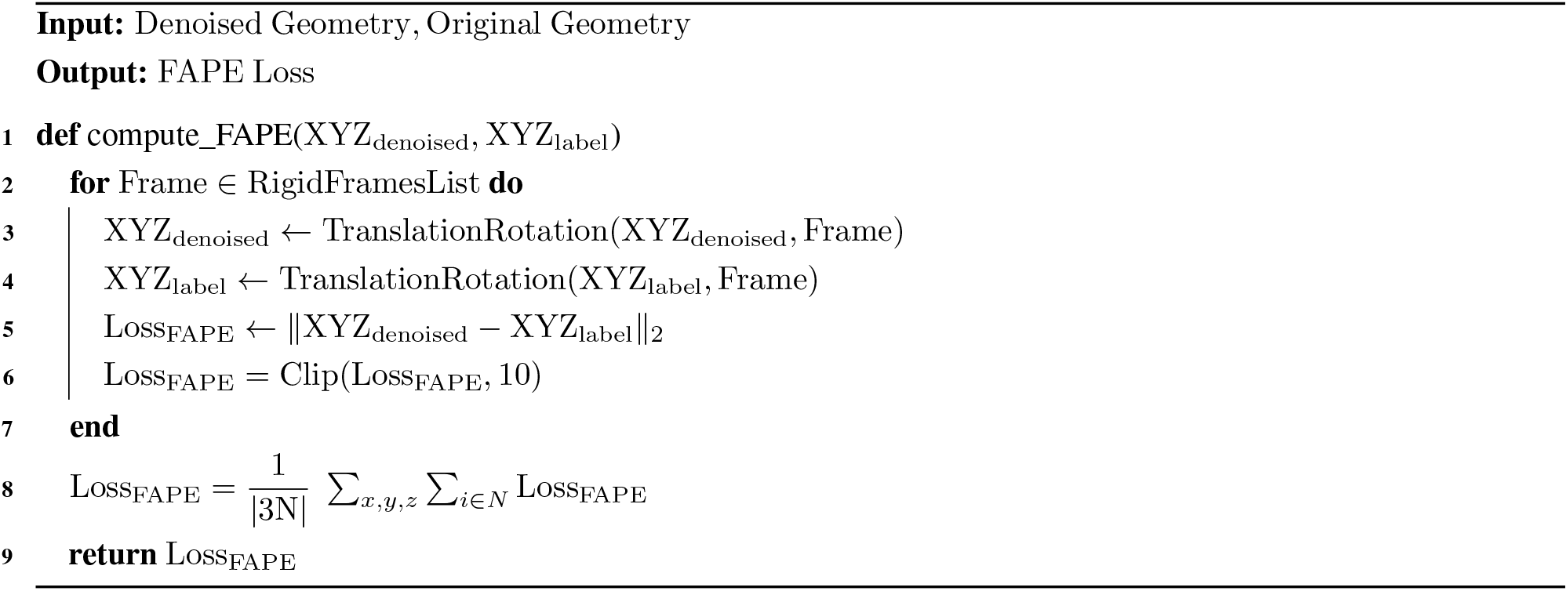

